# The snapdragon genomes reveal the evolutionary dynamics of the *S* locus supergene

**DOI:** 10.1101/2022.07.17.500290

**Authors:** Sihui Zhu, Yu’e Zhang, Lucy Copsy, Qianqian Han, Dongfeng Zheng, Enrico Coen, Yongbiao Xue

**Affiliations:** National Genomics Data Center & CAS Key Laboratory of Genome Sciences and Information, Beijing Institute of Genomics, Chinese Academy of Sciences, Beijing 100101, China; China National Center for Bioinformation, Beijing 100101, China; University of Chinese Academy of Sciences, Beijing 100049, China; State Key Laboratory of Plant Cell and Chromosome Engineering, Institute of Genetics and Developmental Biology, and the Innovation Academy of Seed Design, Chinese Academy of Sciences, Beijing 100101, China; John Innes Centre, Norwich NR47UH, UK

**Author notes:** These authors contributed equally to this work (Sihui Zhu, Yu’e Zhang). Correspondence: Yongbiao Xue.

**Keywords:** snapdragon, *S*-locus, supergene, evolutionary genomics, *SLFs*

## Abstract

The multi-allelic *S*-locus, containing a pistil *S-RNase* and dozens of *S*-locus F-box (*SLF*), underlies genetic control of self-incompatibility (SI) in *Antirrhinum hispanicum*. The genus *Antirrhinum*, harboring such a SI system has been used as a model to study self-incompatibility extensively. However, there have been limited studies on the genomic organization of the *S*-locus supergene due to a lack of high-quality genomic data. Here, we present the chromosome-level reference and haplotype-resolved genome assemblies of a self-incompatible *Antirrhinum hispanicum* line, *AhS_7_S_8_*. Alongside with the draft genome of *Misopates orontium*, comparative genomics reveals that *A.hispanicum* diverged from its self-compatible cousin 12.3 million years ago (Mya). Expanded gene families enriched in distinct functional terms implied different evolutionary trajectories of outcrossing and selfing species. For the first time, two complete *A.hispanicum S*-haplotypes spanning ∼1.2Mb and containing 32 *SLFs* were reconstructed, while most of the *SLFs* derived from retroelement-mediated proximal or tandem duplication approximately 122 Mya. Moreover, we detected a candidate *cis*-transcription factor associated with regulating *SLF*s expression, and two miRNAs may control the expression of this transcription factor. Inter-specific *S*-locus and intra-specific *S*-haplotype comparisons revealed the dynamic nature and polymorphism of the *S*-locus supergene mediated by continuous gene duplication, segmental translocation or loss, and TE-involved transposition events. Our data provides an excellent resource for future research on the evolutionary studies on *S-RNase*-based self-incompatibility system.

## Introduction

Self-incompatibility (SI) is a molecular recognition system that prevents inbreeding and promotes outcrossing in hermaphroditic flowering plants, thereby maintaining genetic diversity and helping angiosperms expand into a wide range of habitats. The molecular mechanisms of SI in eudicots have been extensively studied for decades. In Solanaceae, Rosaceae, Rubiaceae, Rutaceae and Plantaginaceae, SI is controlled by a single supergene called the *S*-locus, with a variety of haplotypes^1^. The *S*-locus generates the pistil *S* determinant (*S*-ribonuclease with cytotoxicity) and the pollen *S* determinant (dozens of pollen-specific *S*-locus F-box genes called *SLFs*) that mediate self/non-self recognition. A unique *S-*haplotype consists of specific *SLFs* and a specific *S-RNas*e. Successful fertilization only occurs between individuals harboring different *S-*haplotypes, and the S-RNase can be degraded by non-self SLF products collectively^2^. Reconstructing the evolutionary history of a supergene is necessary to understand how such a functional module originate and spread in diverse taxons^3, 4^. Supergenes controlling complex phenotypes, which include Batesian mimicry in butterfly^5^ and alternative social organization in fire ant colonies^6^, have long inspired both empirical and theoretical studies on them. However, in plant SI species, most previous reports focused on the discussions of self-incompatible genes themselves, few systematic analyses of the origin and evolution of the *S*-locus at the whole genome level, and the impact of gene duplication on the shaping process of the functional locus was not considered, presenting a critical knowledge gap.

The breakdown of SI results in a mating system change from outcrossing to self-fertilization in angiosperms. SI has been lost frequently during angiosperm diversification^7^, and the transition to self-compatibility (SC) are usually accompanied by accumulation of loss-of-mutations of the *S*-locus^8^. There has been intense interest in the role of parallel molecular changes underlying the evolutionary shifts to selfing associated with loss of SI^9^. Loss-of-function or loss-of-expression of *S-RNase* can confer self-compatibility (SC) in some Solanaceae species^10, 11^, which was also observed in snapdragon^12, 13^. Therefore, it has long been served as an ideal system to explore the impact of mating system shifts and dynamics of adaptation. With the advent of the post genome era, accurate and contiguous genome assembly, the critical basis for understanding genomic diversity and evolutionary trajectories, is available now. In this study, we assembled a chromosome-level reference genome with a total length of ∼480Mb of self-incompatible *Antirrhinum hispanicum* (*AhS_7_S_8_,* heterozygous genotype at *S-locus*, hereafter referred to as ‘SI-Ah’). Also, high-quality draft genomes of self-incompatible *Antirrhinum linkianum* and self-compatible *Misopates orontium* were present here. These sequence resources could provide us an opportunity to perform comparative genomic analyses to capture consequences of the self-incompatibility breakdown and obtain insights into the evolutionary genomics basis of the loci controlling self-incompatibility. We also took the advantage of HiFi data and Hi-C reads to assemble two haploid genomes of SI-Ah, each contains an intact *S*-haplotype. In this study, the complete structure and sequence of the *Antirrhinum S*-locus allowed us to inspect how and when the *SLF* genes proliferate, as well as variations between different *S*-haplotypes. The genomic data and analysis reported in this work are of great value for understanding the genetic bases of adaptive characteristic of species, and provide insights into the evolution of the *S*-locus in snapdragon.

## Results

### Genome assembly and evaluation

We combined different sequencing strategies to derive *A.hispanicum* reference genome assembly: Illumina pair-ended reads, PacBio single-molecule reads, BioNano optical maps, and Hi-C sequencing reads (Fig. S1 and Table S1). The self-incompatible line *A.hispanicum* of *AhS_7_S_8_* was estimated to be with genome size of ∼473.4Mb and a modest heterozygosity of 0.59% (Fig. S2) based on the 21-mer spectrum. Firstly, 36.0Gb of continue long reads (CLR) and 19.1Gb of circular consensus reads (CCS) were obtained on PacBio platform for primary assembly using different assemblers. Then, a total of 203.3Gb BioNano data and 69.9Gb Hi-C paired-end reads were produced to scaffold the contigs into 8 pseudo-chromosomes. In the end, we obtained three sets of chromosome-level genome assemblies of *AhS_7_S_8_*, a consensus assembly with mosaic sequence structure (Fig. 1) and two haplotype-resolved assemblies each contains a copy of the *S*-locus (S_7_ haplome with *S_7_*-haplotype and S_8_ haplome with *S_8_*-haplotype). Yet long stretch of gaps exists in *S_8_*-haplotype of S_8_ haplome and cannot be filled up directly with the sequencing data of *AhS_7_S_8_* due to heterochromatic characteristics of the *S*-locus^14^. Therefore, to overcome these difficulties in assembly of the *S*-locus, we applied a genetic approach using a cross between *AhS_7_S_8_* and self-compatible relative *A.majus* (*S_C_S_C_*)^15^, generating two F1 (*AhS_7_S_C_* and *AhS_8_S_C_*). Since the F1 individuals have haploid chromosome sets from *AhS_7_S_8_*, their sequencing data were used for error correction and gap filling to improve the haplome assemblies. Comparing *A.hispanicum* phased genome sequences with the consensus assembly revealed generally good alignment except some gaps and inversions in few chromosomes, which might be caused by the heterozygous region collapse in Canu assembly (Fig. S4). To facilitate comparative analyses, we also sequenced and assembled the genomes of self-incompatible *Antirrhinum linkianum* and self-compatible *Misopates orontium.* The heterozygosity and genome size of *A.linkianum* is very close to that of *A.hispanicum*, while *M.orontium* exhibits heterozygosity rate of 0.21% and approximately 24.1% reduction in genome size (Table S1).

**Fig. 1.**
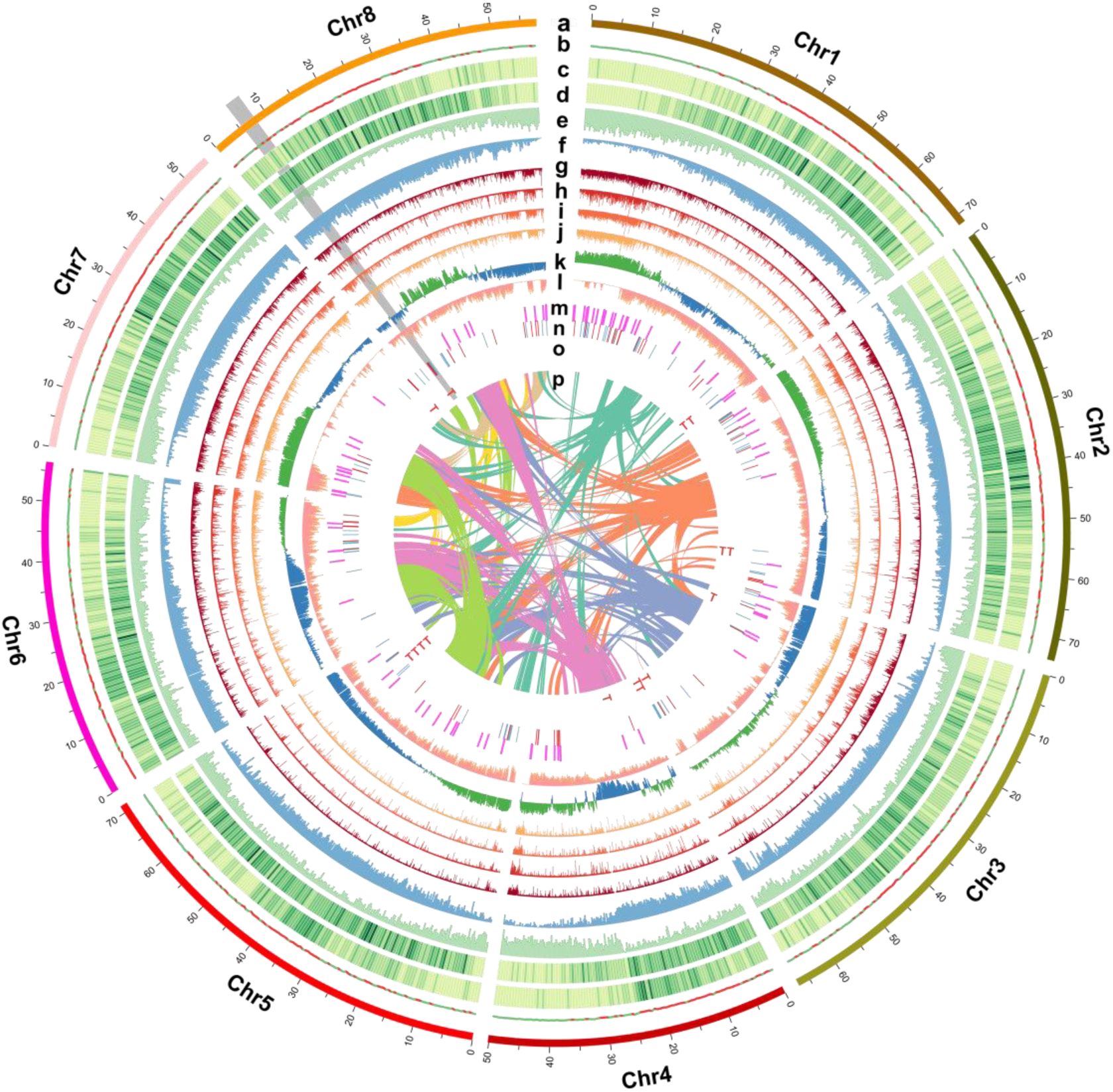
| Genomic features of *A.hispanicum*. The outermost layer is circular ideogram of the eight pseudomolecules. Track **b-j** corresponds to the distribution of the GCcontent (red denote higher than average whole-genome level, and green vice versa), Ty3/Gypsy, Ty1/Copia retrotransposons (higher density shown in darker green), gene density, methylation level, expression level of four tissues (petal, pollen, pistil, and leaf) respectively, all calculated in 250kb non-overlap window. Track **k** corresponds to A/B compartments inferred by using Hi-C data. Track **l** corresponds to SNP/InDel density distribution. Track **m** corresponds to location of miRNA genes. Track **n** corresponds to location of all F-box genes, blue and red represent forward and reverse strand respectively. Track **o** marker “T” corresponds to location of ribonuclease T2 genes. Track **p** corresponds to inter-chromosome syntenic blocks. The grey sector marked *S*-locus in Chr8.

The BUSCO assessment results indicated that 95.0-98.7% of conserved genes were completely recovered in these assemblies (Fig. S6 and Table S6). The S_7_ and S_8_ haplomes were combined and evaluated under the protein mode, and 98.6% (2,090) of BUSCO genes were complete, while 92.2% (1,955) being duplicated. A total of 30,956 *Antirrhinum* EST records downloaded from NCBI (2021-04-23) were mapped to the genome assemblies, with nearly 90% of which can be mapped (Table S5). PE reads from the DNA and RNA libraries from this study were mapped to the genome assemblies respectively. The mapping rate ranged from 70% to 94% (properly mapped or uniquely mapped), as listed in Table S5. In addition, LTR Assembly Index(LAI)^16^ based on LTR-RTs was calculated to evaluate genome assembly continuity. The LAI values of the two haplomes exceed 20, which demonstrates a ‘golden’ standard level, and the draft genome of other two species also reached ‘reference’ grade quality (19.3 and 21.9 for *A.linkianum* and *M.orontium*, respectively). Cumulatively, the results above suggested the reliability of all the nuclear genome assemblies. In addition, circular chloroplast and mitochondrial genomes were also retrieved from contigs assembly, with length of 144.8kb and 527.4kb respectively (Fig. S5). Particularly, the *S-*locus of self-incompatible snapdragon which spans more than 1.2Mb was completely assembled with a continuous BioNano molecular cmap as strong evidence (Fig. S7). The *A.hispanicum* genome assemblies are, to the best of our knowledge, one of the most contiguous and complete *de novo* plant genomes covering the complex *S*-locus supergene to date.

### Genome annotation

Repetitive elements accounted for approximately 43% of *Antirrhinum* genome (Table S4), and long terminal repeats (LTRs) being the largest member of transposons families, covering ∼23% of the nucleus genome (Fig. 1A), of which, the Ty3/gypsy (12.0%) and Ty1/copia (11.6%) are the most abundant transposable elements (TEs). DNA transposons comprises ∼12% of the *Antirrhinum* genomes (Table S4). According to theoretical modeling, if the effects of TE insertion are mostly recessive or co-dominant, increased homozygosity would result in more efficient elimination^17^, which therefore should be excepted have lesser TE in selfing species. As we observed, the total amount of TE in self-compatible *M.orontium* (38.0%) is apparently lower than self-incompatible species (Table S4). The insertion time of intact LRT-RTs in *Antirrhinum* based on the divergence of two flanking terminal repeat sequences revealed a recent burst of LTR-RTs at 0.3 Mya, and indicated that most old LTR-RTs have been lost (Fig. S9). The younger insertions have not accumulated enough substitution in two flanking repeats, making this type of genome more difficult to assemble correctly without long-range information. Gene models of all genome assemblies were predicted followed the same analysis pipeline as described in method. For consensus assembly of *A.hispanicum*, 42,667 protein-coding genes were predicted from the soft-masked genome, 89.3% of which could be annotated in at least one functional database, and other assemblies gave close genome annotation results (Table S7). The average transcript and CDS length of protein-coding genes were 3,023 bp and 1,146bp, respectively. The average exon and intron lengths were 232 bp and 479 bp, with 4.8 exons per gene on average. The length distribution of CDSs, exons, and introns of snapdragon show similar patterns to other plant species (Fig. S10), which confirmed the credibility of gene prediction results. The GC content is distributed unevenly in most pseudo-chromosomes and tends to be higher in gene-poor regions. The density of protein-coding genes on the pseudo-chromosomes was highest in distal regions and declined in centromeric regions (Fig. 1), suggesting that gene distribution is not random. As expected, the pattern of gene presence is opposite to that of LTR-RT elements and methylation level. A total of 2,112 transcription factors belong to 58 families were annotated, including bHLH, ERF, MYB, C2H2, NAC, and B3 families with largest number of members (Fig. S11). Analysis of small-RNAseq libraries from four tissues identified 108 known miRNA genes of 42 families of *A.hispanicum* and 105 miRNA genes of 42 families (details listed in Table S9). In addition, 719 transfer RNA genes (tRNAs), 311 small nuclear RNA genes (snRNAs), and 791 ribosomal RNA (rRNAs) genes for *A.hispanicum* were predicted (Table S8).

Taken together, three parts of snapdragon genome (nuclear, chloroplast, mitochondria) are of high completeness and correctness, making them a valuable resource for comparative and evolutionary studies.

### Comparative genomic analysis

We employed the consensus-assembly to represent *A.hispanicum* in comparative genomic analysis to conduct phylogenomic analysis. Proteomes from *A.hispanicum, A.linkianum*, and *M.orontium*, together with 15 other species (Table S10) were clustered into 36,295 orthogroups (Table S11) and used for phylogenetic relationship construction subsequently. In Lamiids, the family Plantaginaceae, to which *Antirrhinum* and *Misopates* belongs, form a monophyletic group that diversified recently. The species tree indicated that the divergence of Plantaginaceae from Lamiales order common ancestor occurred ∼55.8 Mya during Paleocene-Eocene transition (Fig. 2A). After that, *M.orontium* diverged from *Antirrhinum* genus and lost self-incompatibility assuming self-compatibility is ancestral state in Plantaginaceae^7^. *A.hispanicum* and *A.linkianum* split at around 3.5 Mya [95% CI=1.6-5.3 Mya], which is collided with emergence of modern Mediterranean climate and rapid speciation of *Antirrhinum* lineage, and in agreement with previously reported results using various datasets and methodologies^18^.

**Fig. 2.**
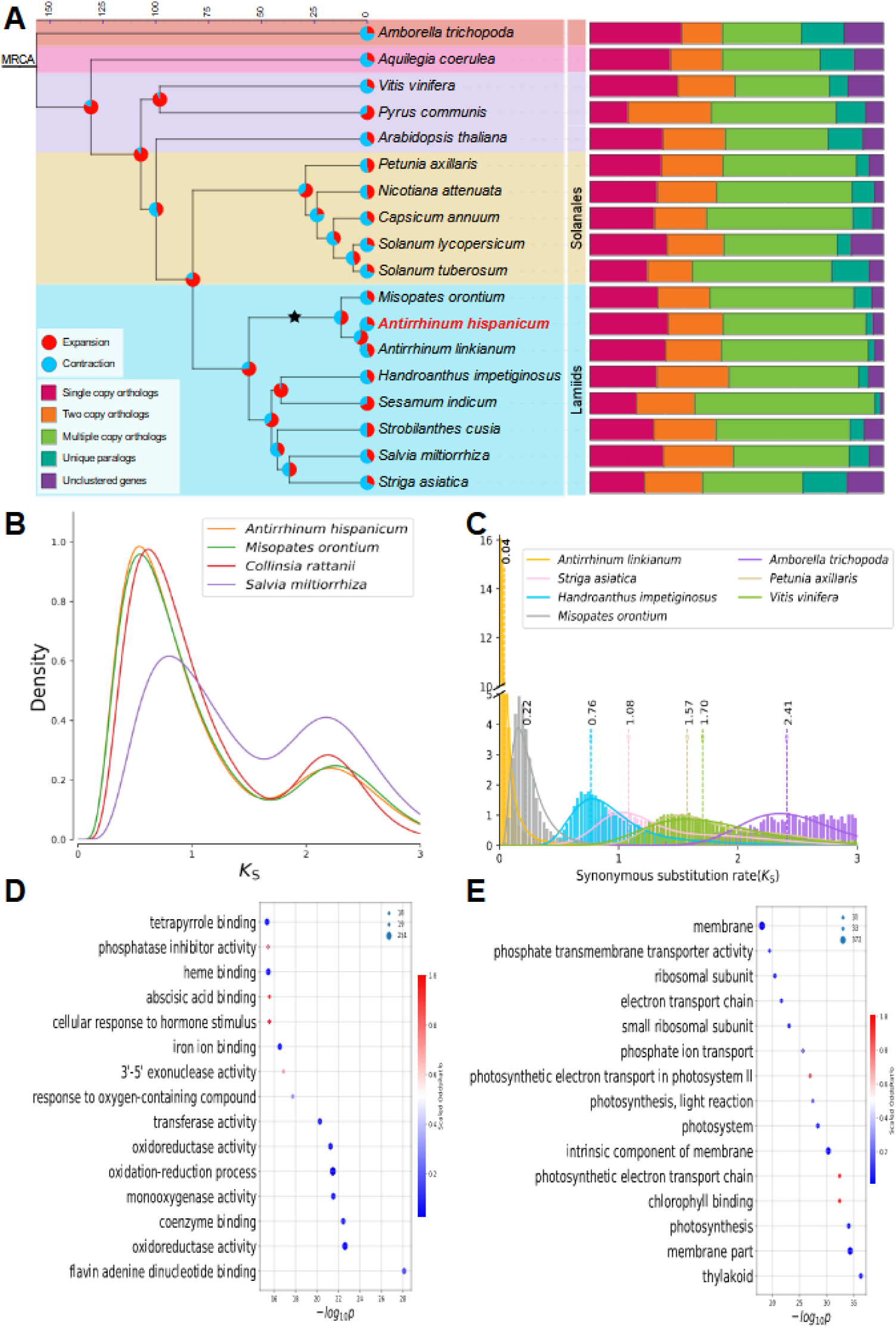
| Comparative genomics analysis and lineage expanded gene families of *Antirrhinum*. A Phylogeny position of the snapdragon and gene family evolution. The left panel shows phylogenetic tree of 18 species. The black star denotes the Plantaginaceae-specific WGD. The pie diagram on each branch of the tree represents the proportion of gene families undergoing expansion (red) or contraction (blue) events. The right panel shows the distribution of single copy, two copy, multiple copy, unique paralogs, and other orthologs in plant species. **B** Paranome *Ks* distribution of three Plantaginaceae members and *Salvia miltiorrhiza*. **C** *Ks* distribution of one-to-one orthologs identified between *A.hispanicum* and other species. The density curves were fitted using Gaussian mixture model, and the peak values marked with dotted dash lines. **D-E** Gene Ontology enrichment analysis of expanded family members in *Antirrhinum* lineage and *M.orontium* branch. The color of circle represents the scaled odds ratio. The size of circle represents the gene number of GO terms.

To identify WGD events in *Antirrhinum*, we first compared the SI-Ah genome with the reconstructed ancient eudicot (AEK)^19^ genome, and *Vitis vinifera* genome which have underwent none large-scale genome duplication after whole-genome triplication (WGT) of AEK^20^. Synteny analysis of the SI-Ah with AEK and *Vitis vinifera* genomes showed synteny depths ratio of 1:6 and 1:2 respectively (Fig. S12), and large-scale segmental duplications can be observed from the self-alignment of SI-Ah (Fig. 1). The *Ks* distribution of paranomes displayed two peaks (Fig. 2B), indicating *Antirrhinum* genus have at least experienced another one WGD after the gamma-WGT.

To approximately date the WGD in snapdragon, the distributions of synonymous substitutions per synonymous site (*Ks*) among paralogous genes within a genome were examined. The distributions plots of *Ks* for *A.hispanicum*, *C.rattanii*^21^ and *A.linkianum* paranomes showed a distinct peak at 2.21 echoing the well-reported gamma-WGT occurred in the common ancestors of all eudicots approximately 117 Mya^22^(Fig. 2B and S14), as well as an additional peak at ∼0.61 corresponding to a later duplication event shared among Plantaginaceae members. Therefore, the younger WGD was dated to occur at ∼32.3 Mya, posterior to divergence of *Antirrhinum* and other Lamiales plants, indicating a Plantaginaceae-specific WGD event. The *Antirrhinum* lineage neutral mutation rate was inferred using the formula μ=D*/2T*, where D is the evolutionary distance of two *Antirrhinum* species (peak value of *Ks* log-transformed distribution = 0.04, Fig. 2C), and T is the divergence time of the two species (3.5Mya). The neutral mutation rate for *Antirrhinum* genus was estimated to be 5.7e-9 (95% CI: 3.8e-9 -1.3e-8) substitution per site per year, which is very close to the result estimated by using genotyping-by-sequencing dataset^18^.

Loss of self-incompatibility occurred many times during angiosperms evolution history^7, 8^. Although two species have split up for millions of years, synteny block analysis revealed strong collinearity between *A.hispanicum*and *M.orontium* (Fig. S13). And to investigate the evolutionary trajectory of SI- and SC-snapdragons, gene family analysis was conducted based on orthologous relationship of 18 plant species (Fig. 2A). A total of 445 and 294 gene families were expanded and contracted respectively in *Antirrhinum* lineage. The results also revealed 704 (3,230 genes) expanded and 1,692 (2,956 genes) contracted gene families in *A.hispanicum*, 1,097 (4,787 genes) expanded and 1,392 (4,015 genes) contracted gene families in *A.linkianum* respectively. (Fig. 2A). Moreover, 702 gene families (4,305 genes) were found in expansion, while 1209 families (1,771 genes) were found in contraction in the self-compatible *M.orontium*. These rapidly evolving gene families might give us some clues about consequences of losing self-incompatibility in *M.orontium* lineage.

The GO enrichment analysis for 445 expanded gene families of self-incompatible *Antirrhinum* branch showed that the significantly enriched GO terms mostly involved ‘binding’, ‘enzyme activity’, and ‘response’ (Fig. 2D). While in *M.orontium*, enriched GO terms of expanded families mostly related to photosynthesis (Fig. 2E), and KEGG enrichment analysis revealed that the expanded genes were mainly implicated in phenylpropanoid, flavonoid, sesquiterpenoid and triterpenoid biosynthesis (Fig. S16), which may contribute to the survival of self-compatible species facing environmental changes on the way to an evolutionary ‘dead-end’^23^. For the complete list of GO enrichment analysis results, see Table S12 and S13.

### Position, structure, and flanking genes of the *S*-locus supergene

Identifying sequence details of the *S*-locus supergene would be the initial step towards understanding its evolution in snapdragon. The *Antirrhinum S*-locus was found to locate in peri-telomeric region on the short arm of chromosome 8^14^, as opposed to sub-centromeric area in *Solanaceous* species. A total of 275 genes encoding F-box protein were annotated in the entire snapdragon genome (Fig. 1). A cluster of 32 *SLF* (*S*-locus F-box) genes, which match to the structural characteristics of the candidate *S*-locus, located in chromosome 8 (Fig. 1). Gapless *S*-locus of 1.25Mb in *A.hispanicum* and pseudo *S*-locus of 804kb in *A.majus*^15^ are both reconstructed with BioNano long-range molecules as support evidence (Fig. S7 and S8), whereas the *S*-loci of *A.linkianum* and *M.orontium* were identified by synteny alignment with *A.hispanicum*. Except *SLFs*, the *S*-locus also contains genes encoding G-beta repeat, matE, zinc finger, cold acclimation, auxin responsive proteins, and dozens of genes with unknown function or TE-related function (listed in Table S16). Additionally, the *S*-locus was surrounded by two copies of organ specific protein at left side (OSP1: *AnH01G33547.01*, OSP2: *AnH01G33548.01*) and five copies of phytocyanin genes at right side (here named PC1: *AnH01G33649.01*, PC2: *AnH01G33650.01*, PC3: *AnH01G33651.01*, PC4: *AnH01G33652.01*, PC5: *AnH01G33653.01*) (Fig. 3E). The contiguity and completeness of the genome assembly, together with solid annotation, give rise to the possibility of proposing an origin model of the *S*-locus supergene. There is no doubt that gene duplication plays a key role in shaping process of such a complex supergene. It has been proved that tandem or proximal duplicates evolved toward functional roles in plant self-defense and adaptation to dramatic changing environments^24^. To inspect how multiple *SLFs* were generated and proliferated, phylogenetic tree of 275 *SLFs* was built (Fig. 3A). So, if all the *SLFs* originated via WGDs or large segmental duplication, their paralogs should colocalize elsewhere in the genome and have similar divergence with the *S*-locus genes. On the contrary, if *SLFs* were the results of stepwise local duplications, they would exhibit less differences and there should be more distant paralogs scattered elsewhere in the genome. The *SLFs* phylogenetic tree proves that the situation is the latter one. The phylogenetic tree along with expression heatmap of 275 F-box genes indicated *SLFs* not only share closer relationship, but also display similar pollen-specific expression pattern which distinguished from other F-box genes (Fig. 3A). This phenomenon has not changed much in cultivar *A.majus* after its domestication (Fig. S21). Furthermore, based on classification of gene duplication type, the two OSPs derived from the gamma-WGD followed by a recent tandem duplication (Table S20), while the PCs arise from tandem duplication asynchronously assuming constant substitution rate, both can be supported by gene phylogenetic tree topology of their paralogs (Fig. S22, S23). As for *SLFs*, 24 and 6 of them are proximal duplication and tandem duplication genes respectively, and other 2 *SLFs* are dispersed duplication derived (Table S16). Besides, 27 of 32 *SLFs* are intron-less, suggesting *SLFs* proliferated most likely through retroelement-mediated way. *Ks* distribution of pairwise *SLFs* and non-*S*-locus F-box genes were compared, and the synonymous substitutions rates of *SLF* paralogs are much lower than non-*S*-locus F-box paralogs (Fig. 3B). With the estimated *Antirrhinum* mutation rate, the peak value of *SLFs* paralogs *Ks* distribution (ranging from 1.35 to 1.40) corresponding 122 Mya^25^ (Fig. 3B), indicated that the *S*-locus structure is a very conserved and long lived supergene survived from a long-term balancing selection^26, 27^. This observation concludes that the *S*-allelic polymorphisms should be inherited from common ancestors and the differentiations began before the currently extant species formation. Dissection of *SLFs* and flanking genes indicate that the *S*-locus is still experiencing gene gaining in *Antirrhinum* (Table S20).

**Fig. 3.**
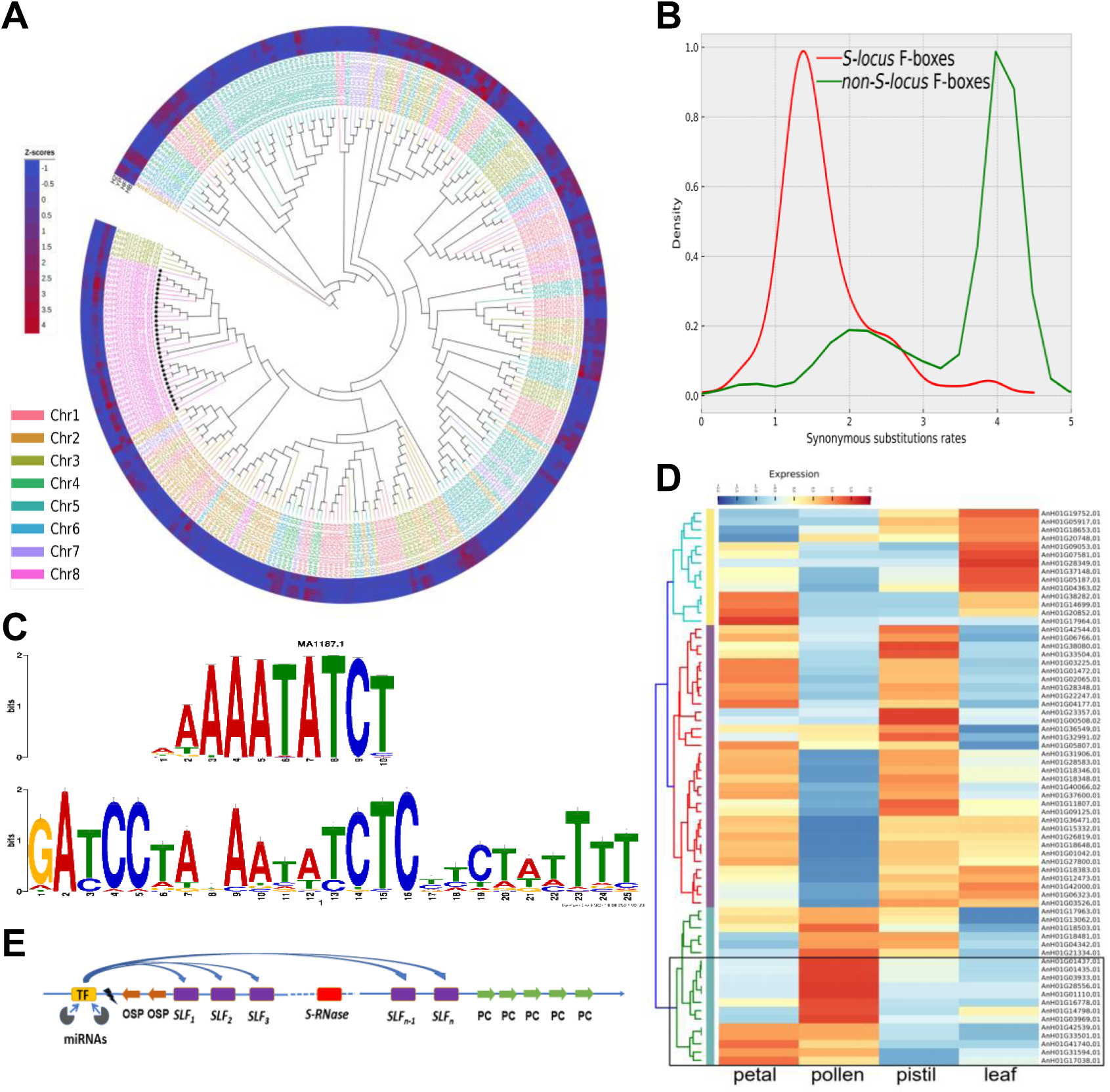
| Identification of a candidate regulator of *SLFs*. **A** Phylogenetic tree reconstructed based on the amino acid sequences of all F-box genes from *A.hispanicum* genome. The maximal likelihood phylogenetic tree was calculated using RAxML-NG with 1000 bootstrap. The color of gene ID denoted the chromosome the gene within as indicated in the legend. Heatmap color represent Z-scores derived from RNA-seq expression data for each gene. *SLF* genes are marked with black stars. **B** The red line is synonymous substitution rate distribution of 32 *SLFs*, while the green one denotes other non-*SLF* F-box genes in the genome. **C** The bottom panel represents motif found by scanning upstream 1000bp of 32 *SLFs* using MEME, and the top panel is a Tomtom alignment for annotation the putative transcription factor binding site. **D** The cluster heatmap of expression matrix of all MYB-related transcription factor family members in SI-Ah genome. The columns represent 4 tissues. The legend shows the range of log2(TPM+0.5), and TPM is mean value of two biological replicates. Genes marked in black box may be involved in controlling the expression of *SLFs*, since they are expressed in pollen but not in pistil. **E** Schematic structure and regulatory relationship of snapdragon *S*-locus.

Using MEME suit analysis, we identified a significant motif (GATCCTAXAATATCTC) located upstream of most *SLFs*, and the motif annotation implied *SLFs* are likely to be co-regulated by a MYB-related family transcription factor (Fig. 3C). By examining expression of all the MYB-related family members in the genome, *AnH01G01437.01, AnH01G01435.01, AnH01G03933.01, AnH01G28556.01, AnH01G01110.01, AnH01G16778.01, AnH01G14798.01, AnH01G03969.01, AnH01G42539.01, AnH01G33501.01, AnH01G41740.01, AnH01G31594.01,* and *AnH01G17038.01* were expressed in pollen but barely in pistil tissue (Fig. 3D). Among them, *AnH01G33501.01* is located about 1.2Mb upstream of the *S*-locus, making it a potential regulator involved in activating expression of the *SLFs*. Moreover, by examining miRNA target prediction results, two candidate miRNAs (aof-miR396b and zma-miR408b-5p) were revealed to bind with different regions of *AnH01G33501.01*, and their expression patterns exhibited negative correlation with the TF (Fig. S25). The miR396 family member, aof-miR396b, was believed to influence GRF transcription factors expression^28^, while over-expression of zma-miR408b in maize were reported to respond to abiotic stress to maintain leaf growth and improve biomass^29^. The results implied that those two miRNAs may also play an important role in self-incompatibility.

From the intraspecies similarity matrix heatmap of *SLFs* of SI-Ah, we can also observe the divergence among paralogs are much higher than interspecific orthologous pairs (Fig. S24). The pairwise sequence identity of 32 *SLF* paralogs ranged from 33.1 to 77.3%. And the closer the physical distance between two *SLFs*, the higher the sequence similarity and identity (Fig. 4A). In addition, the values of the non-synonymous/synonymous substitutions rate ratio (Ka/Ks) of nearly all *SLFs* from different allele below one is a signature of purifying selection to maintain necessary protein functions to detoxify S-RNase collectively (Fig. 4B).

**Fig. 4.**
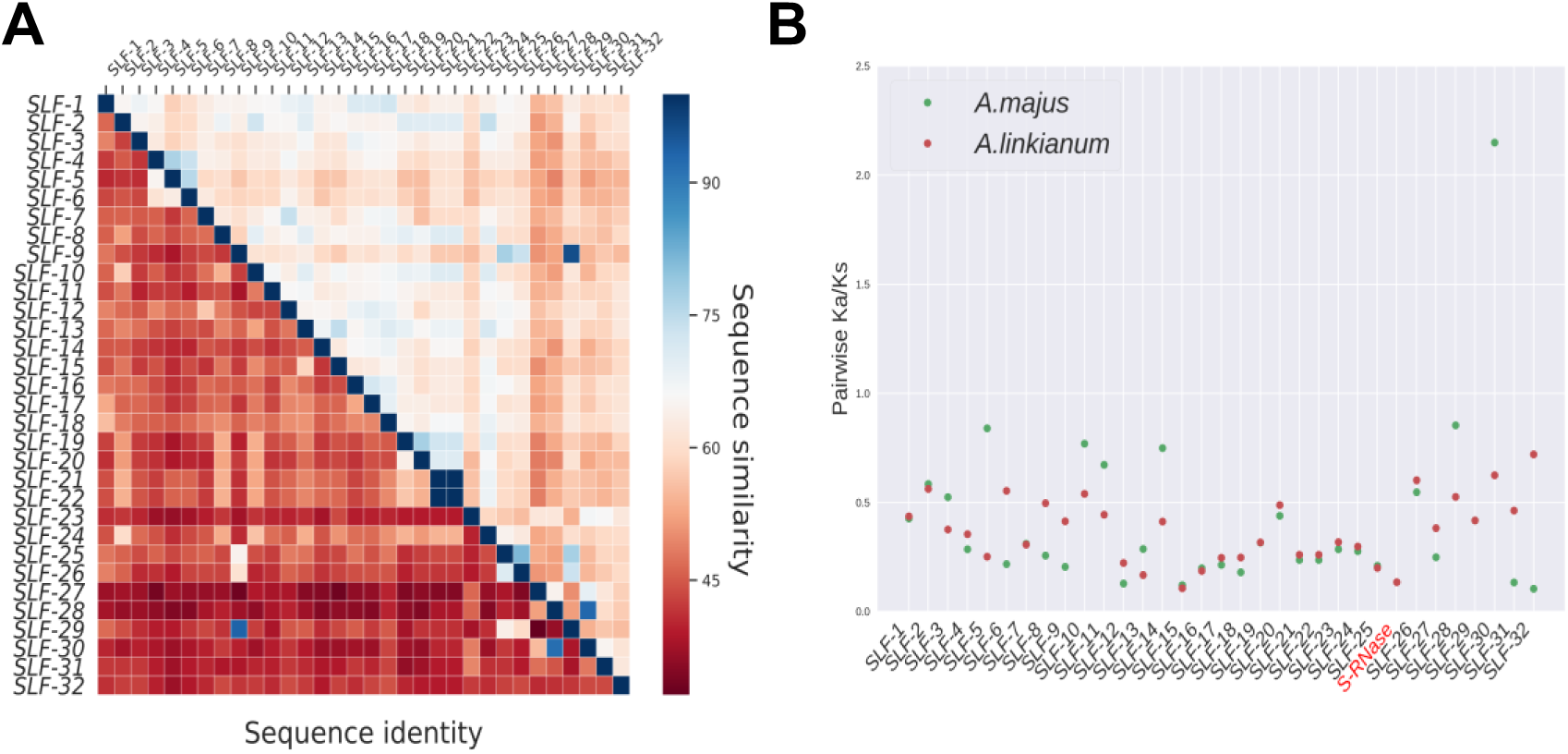
| The correlation of position and similarity of *SLFs*. **A** Heatmap showing sequence identity (lower left triangle) and sequence similarity (upper right triangle) between *SLFs* in SI-Ah. The genes in x-axis and y-axis are ordered by physical position on the chromosome. **B** Pairwise Ka/Ks analysis for all *SLF*s and ribonuclease gene in the *S*-locus between SI-Ah and *A.majus,* as well as *A.linkianum*. Genes are ordered by genomic coordinates, and label of ribonuclease gene is highlighted in red.

### Evolution history of the Antirrhinum S-locus

Comparison of microsynteny revealed the presence of the *S*-locus remnant in other *S-RNase*-based self-incompatible species, including *Chiococca alba*^30^ (Rubiaceae), *Citrus clementina*^31^ (Rutaceae), *Physalis pubescens*^32^ (Solanaceae), *Prunus davidiana*^33^ (Rosaceae), but not in the syntenic regions of species outside Eudicotyledoneae, such as *Oryza sativa* (Poaceae). The distribution pattern also suggested that the *S*-locus emerged early right after the divergence of Eudicotyledoneae from with other Mesangiosperms. In *P.davidiana*, core part of the whole *S*-locus was translocated to a different chromosome, and only four *F-boxes* are still there occupying a drastically contracted region (Fig. 5A). Among the four genes, two of them have 4 and 3 orthologous *SLFs* in SI-Ah implying that the multiple copies formed after species divergence. The same contraction was observed in *Chicocca* except with a few more F-box genes. Whereas in citrus genome, none of the homologous *SLFs* were found. However, in *Physalis pubescens*, a Solanaceae member phylogenetically closer with *Antirrhinum*, almost all *SLFs* were lined up in same order with *Antirrhinum* and well preserved. Therefore, it would be reasonable to infer that the protype of extant *S*-locus with only several *SLFs* and *S-RNase* appeared in the common ancestor of eudicots^25, 34, 35^. Then, accompanied by speciation, *SLFs* genes underwent loss and gain at different rate in different lineages (Fig. 5F). If the new *SLFN* copy can recognize more non-self S-RNase^12^, then it is advantageous for carrying this new duplication and spreading through the population by genetic drift^36^. Gene acquisition was most likely through tandem or proximal duplications from ancestor copies, and the losses mainly involved large fragment deletion.

**Fig. 5.**
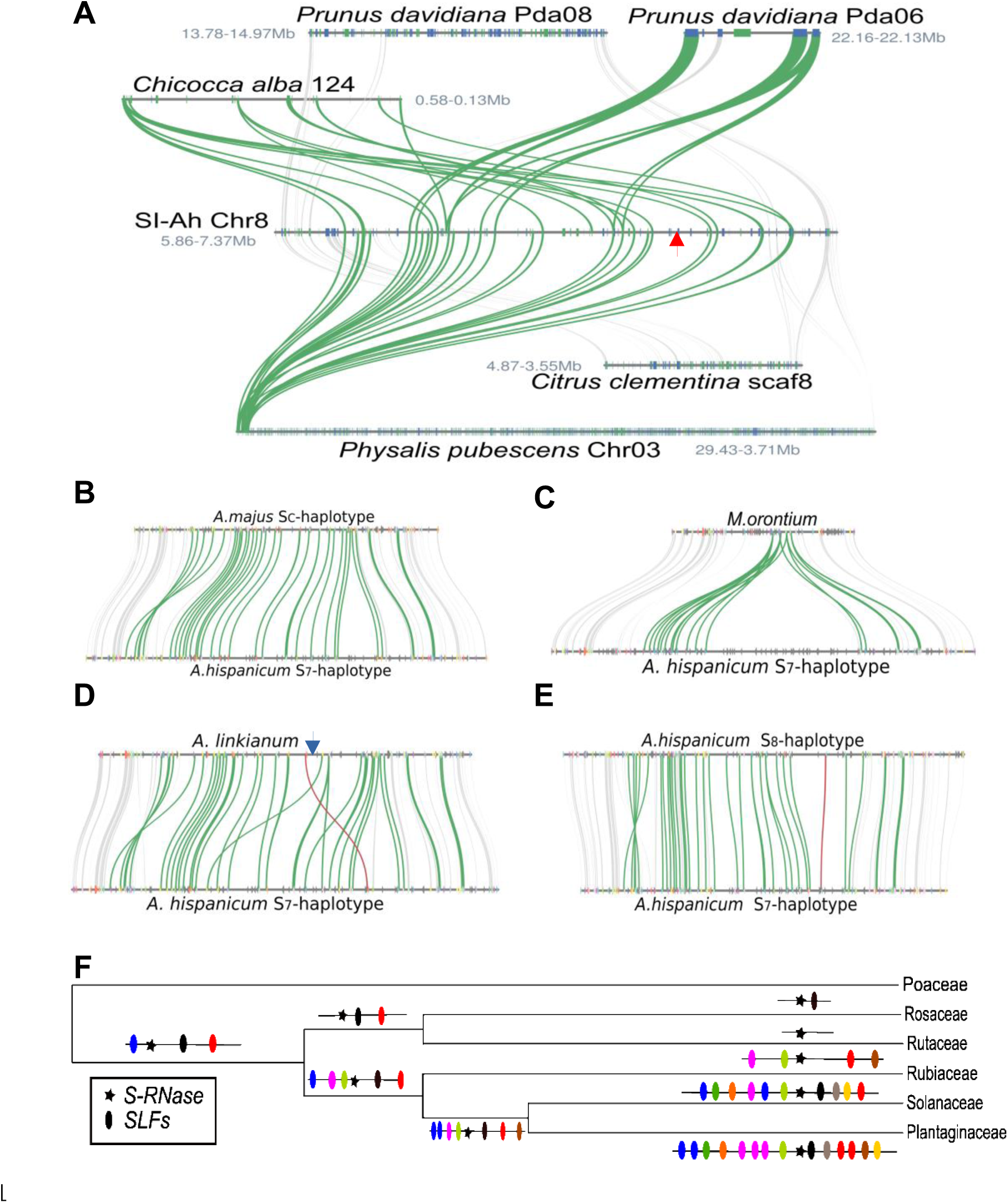
| Evolution of the *Antirrhinum S*-locus supergene. **A** Microsynteny of the S-locus between *A.hispanicum* with other four species (*Chiococca alba*, *Citrus clementina*, *Physalis pubescens*, and *Prunus davidiana*). The red arrow denotes the *S-RNase* of *A.hispanicum*. **B-D** The micro collinearity patterns of *S*-haplotype in *A.majus*, *M.orontinum*, and *A.linkianum*, with *S_7_*-haplotype from *A.hispanicum*. **E** The microsynteny of *S_7_*-haplotype vs. *S_8_*-haplotype. The green curve lines and the red lines denote *SLFs* and *RNase* gene respectively, other genes are colored grey. The Tam downstream 11kb of *S-RNase* is marked by a blue arrow, and the TE is on the same strand with *S-RNase*. **F** Possible formation process of the extant *Antirrhinum S*-locus. The color of *SLFs* represents newly evolved ones to deal with the evolution of the *S-RNase* with new specificity.

The continuity of genome assemblies in this study also provides an opportunity to make comparison of *S*-haplotypes from same or very close species. For the sake of simplicity, we use ‘*S_7_*-locus’ and ‘*S_8_*-locus’ to refer the *S*-locus from the *S_7_*- and *S_8_*-haplotypes containing *S-RNase* and *SLF*s respectively, and their *SLF* genes as ‘*S_7_*-*SLF*’ and ‘*S_8_*-*SLF*’, while Al-*S*-locus refers the *S*-locus of self-incompatible *A.linkianum*. Am-*S_C_*-locus and Mo-*S_C_*-locus which contain no *S*-*RNase* denotes ψ*S*-locus of self-compatible *A.majus* and *M.orontium*. There are 32 *SLFs* annotated in the region of *S_7_*-locus (*S_7_*-*SLF1* ∼ *S_7_*-*SLF32*) and *S_8_*-locus (*S_8_*-*SLF1* ∼ *S_8_*-*SLF32*) respectively. Their position in chromosome and sequence information were listed in Table S16. We compared different *S*-haplotypes to identify polymorphism of the supergene.

Synteny plot showed structure and gene content of the flanking ends of different *S*-haplotypes were well conserved. The *S_7_*-locus has high collinearity with Am-*S_C_*-locus, *S_8_* locus and Al*-S-*locus except a small inversion at the start of alignment (Fig. 5B-E). From the comparison between Mo-*S_C_*-locus and *S_7_*-locus, a large deletion with a length of about 556kb could be observed, which involved *S-RNase* and more than ten *SLFs* as well as dozens of functional genes in *S_7_*-haplotype. These genes include ATPase, Cytokinin, DUF4218, other ones mostly are uncharacterized proteins. This structural variation can be responsible for the loss of self-incompatibility in *M.orontium*. Whereas between the *S_7_* and *S_8_* locus, a small inversion could be observed, which may involve with recombination suppression, and serval neighbor genes of *S_7_*-*RNase* have no corresponding orthologs in *S_8_* haplotype (Fig. 5E), suggesting a divergent evolution history in different regions of the *S*-locus. The relative position of *S*-*RNase* in Al-*S*-locus being different from *A.hispanicum* (Fig. 5D), indicated that the changing physical position *S-RNase* does not impact its function and the dynamic change of the polymorphic *S*-locus in *Antirrhinum* has been very active. And we did observe a nearby CACTA transposable element in the vicinity of *A.linkianum S-RNase*, *Tam*, which may lead to the gene transposition.

Together, these results revealed the dynamic nature of the *S*-locus supergene during evolution and speciation in Eudicotyledoneae, which involves continuous gene duplication, segmental translocation or loss, and TE-mediated transposition.

### Allelic expression and DNA methylation profiling

Since haplotype-resolved assembly is available and gene order is highly consistent between two haplomes (Fig. S27-34), we can determine two alleles of a physical locus and investigate allelic gene expression directly. Base on synteny and homology relationship, 79,398 genes (90.6% of all predicted genes) were found to have homologs on the counterpart haplotype (see “Method”). Most allelic genes displayed low level of sequence dissimilarity (Fig. 6A). To understand the expression profile of allelic genes, RNA-seq datasets were analyzed using allele-aware quantification tools with combined coding sequences of two haplomes as reference. In the survey across four different tissues, most expressed loci did not exhibit large variance, only 2,504 gene pairs (3.2% of total genes, see Fig. S36) displayed significant expression imbalance between two alleles (|fold-change| ≥ 2, p ≤ 0.01), suggesting that most alleles were unbiasedly expressed in the SI-Ah genome. We also analyzed four WGBS libraries to profile methylation state of two haplomes. About 51.6-59.5 million PE reads were generated for each sample (Table S2). About 92.4-92.9% of total reads can be properly mapped to the haplomes. Reads for each sample were mapped to the snapdragon chloroplast genome too, the conversion rate higher than 99% confirming reliability of the experiments. Methycytosines were identified in CpG, CHH, and CHG contexts. A large fraction of cytosines in CpG (∼80.6%) and CHG (∼55%) contexts were methylated, but a smaller fraction of cytosines in CHH (∼8%) context were methylated. No significant methylation level variations were observed between two haplomes (Fig. S27-34), similar to that reported in potato^37^.

**Fig. 6.**
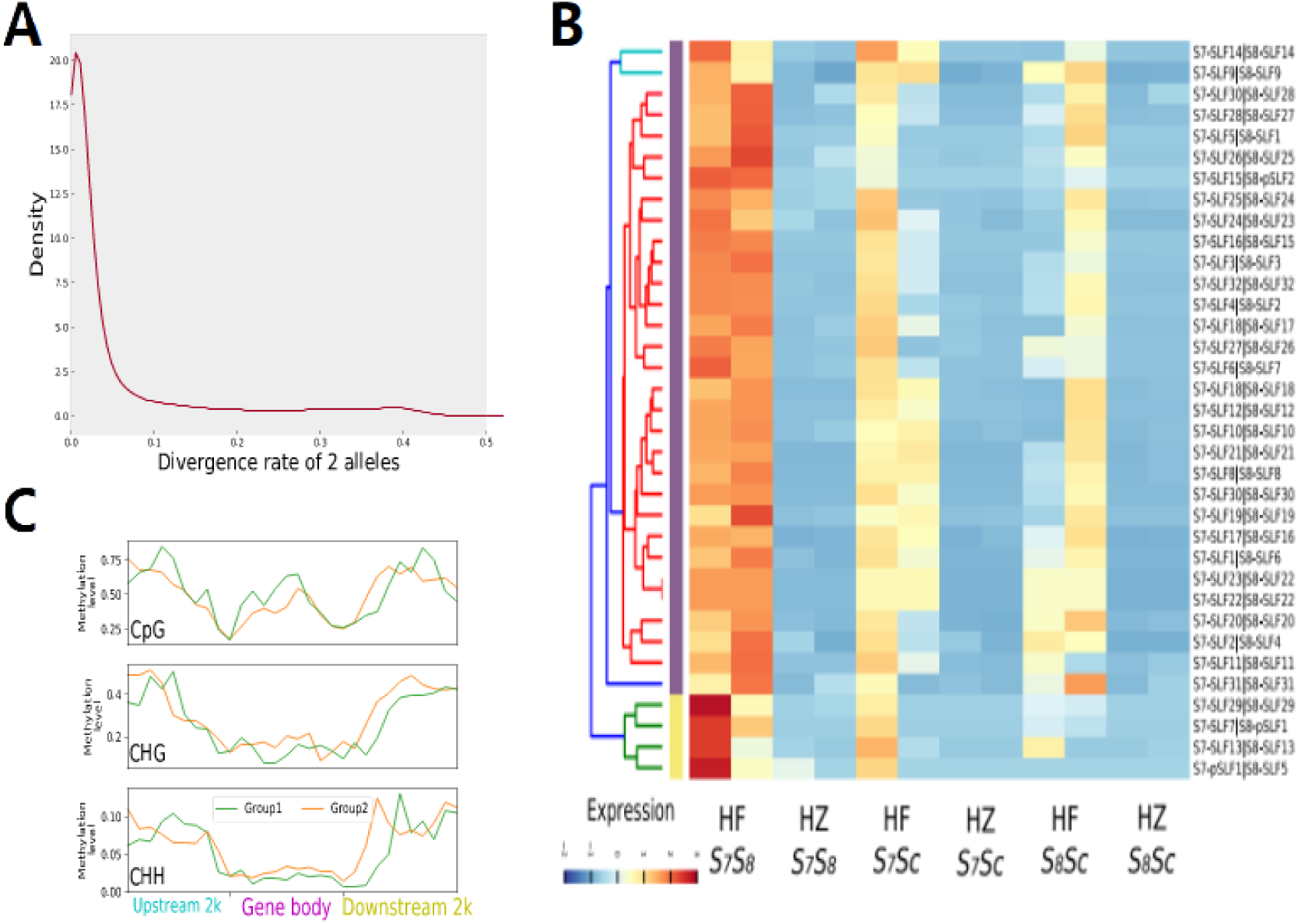
| Allelic expression and methylation profile of *SLFs* for *AhS7S_8_ of A.hispanicum*. **A** The distribution of allelic divergence of genes. The divergence was measured using Levenshtein edit distance. **B** Allelic expression heatmap of *SLFs* in three genotypes. Every two columns represent allelic gene pairs of two *S*-haplotypes (HF: pollen, HZ: pistil). *SLF* starts with ‘p’ indicated pseudo-F-box gene lack of F-box domain or F-box protein interaction domain, but they are the best hit of the counterpart F-box allele. **C** Methylation level within group1 (higher expressed) and group2 (lower expressed) *SLFs* and their 2kb flanking (upstream and downstream) regions in CpG, CHG, and CHH context. Each region was divided into 10 bins of equal size and the methylation level in each bin is displayed in the line graph.

After examining *SLFs* and *S*-*RNase* expression of two haplotypes, we found that *S_7_*-*RNase* and *S_8_*-*RNase* expressed indiscriminately and fairly high in *AhS_7_S_8_* pistil (Table S18), which is consistent with the situation in crab cactus^38^, and they are nearly undetectable in leaf and petal. Those two genes only expressed in pistils of *AhS_7_S_C_* and *AhS_8_S_C_* respectively as expected, yet at higher level than that of *AhS_7_S_8_* pistil. Since the gene sequences did not change after one crossing experiment, it is very likely that RNase expression was changed through other mechanisms. As for the expression profiles of *SLFs* in pollen tissue of three different genotypes, *S_7_*-*SLFs* and *S_8_*-*SLFs* did show an overall decline in two offspring samples but tend to be reduced generally. Those genes could be divided into two groups. Group1 consists of 4 genes, were highly expressed in pollen of *AhS_7_S_8_* with obvious tissue-specificities, but barely expressed in two hybrid progenies, and these genes displayed allelic bias. Another 28 *SLFs* in cluster 2 maintain expression patterns in three genotypes, but the expression level in two hybrid progenies are slightly lower than that of *AhS_7_S_8_* (Fig. 6B). We assumed that group2 are necessary for degrading toxic S-RNase.

Epigenetic modifications are believed to be crucial in controlling gene expression^39^. To determine whether methylation level is responsible for the difference between two groups, we compared the DNA methylation in the pollen from *AhS_7_S_8_*. We calculated the average DNA methylation rates for the *SLFs* gene regions and found that DNA methylation in gene body are higher than flanking regions and the highest methylation rate of mCs was in the CpG context, followed by CHG and CHH (Fig. 6C). The average methylation rates of two groups in the CpG, CHG, and CHH are comparable in gene body and flanking regions. Thus, the similar levels and patterns of methylation of two groups of *SLFs* may not significantly explain the variance in expression. Therefore, the difference may be caused by other kinds of factors. Thus, our data show that some genes are grouped into one cluster can be regulated differentially, irrespective of their relative position within the locus.

## Discussion

Assembling genomes of self-incompatible plants has always been a challenge due to inevitable heterozygosity and large portions of repetitive sequences. In current study, we utilized multiple sequencing technologies and the genetic strategy to obtain chromosome-level genome of self-incompatible *A.hispanicum*. Benefiting from the robust assembly, we can determine the genomic landscape of the first self-incompatible snapdragon. Along with high quality genomes of other two Plantaginaceae species, *A.linkianum* (2n=16) and *M.orontium* (2n=16), these assemblies would be valuable resources and enabled us to understand the genome evolution of *Antirrhinum* lineage, gaining knowledge of the proximate and ultimate consequences of losing self-incompatibility^8^. Apart from the shared gamma WGT in all eudicots, *A.hispanicum* underwent a Plantaginaceae-specific WGD event dated back to the Eocene, which may have been involved in large scale chromosomal rearrangement and TEs proliferation. The phylogenomic analysis clearly reveals that *A.hispancium* split from its relative *A.linkianum* during the Pliocene Epoch, coinciding with establishment of modern Mediterranean climate in the Mediterranean basin^40^. Mating system shift is ubiquitous throughout angiosperms, the evolutionary consequences of transition have attracted extensive attention for a long time^41, 42^. The comparative genomic analysis indicated that after losing self-incompatibility, though the genomic architecture of SC-*M.orontium* has not changed much, the gene contents have undergone certain fluctuations. To increase environmental tolerance, the self-compatible organism recruits alternative biological metabolic pathways to improve survival rate, and produced more plant compounds to cope with various external stimuli, since it is no longer possible for self-compatible species through recombination to generate new variants to adapt to the volatile environments.

Genetic architecture and evolutionary history of the supergenes have long fascinated biologists. Owning to the long-range sequencing information, the complete *S*-locus consisting of dozens of genes was successfully reconstructed, and our results indicated that *SLF* members within this gene cluster share common *cis*-regulatory elements, ensuring a spatially coordinated expression of protein products to degrade *S-RNase*. Additionally, the physical position and expression pattern of *AnH01G33501* suggested that it may be a candidate transcription factor regulating downstream *SLFs* expression. Moreover, two miRNAs were considered to act as regulators to control expression of this transcription factor^43^. Based on calculation of Ks of *SLFs*, we found that the ancestral *SLF* had an ancient origin. During the long evolutionary process, different ancestral *SLFs* faced different fate (deletion, expansion, contraction, pseudogenization) in different lineages. Frequent and rapid acquisition of new *SLFs* were adapted to the allelic diversity of *S-RNase*^12^ and contributed to the complexity of the extant *Antirrhinum S*-locus supergene. Duplication type analysis of all the extant *SLFs* revealed that most of them originated earlier than two WGDs in Plantaginaceae through tandem or proximal duplication, yet the flanking genes of the *S*-locus are still expanding gradually. Thus, the snapdragon *S*-locus supergene has charm of both old and young.

Structural variations such as translocations and inversion are considered drivers of genome evolution and speciation^44, 45^. These rearrangements have likely contributed to the divergence of these species because they can lead to reproductive isolation if they are large enough to interfere with chromosome pairing and recombination during meiosis^46^. By comparing sequence of two *S*-haplotypes, we speculated such an inversion at this locus may be involved in recombination suppression, thus maintain the integrity and specificity of every single haplotype. Inter-specific and intra-specific *S*-haplotype comparison revealed continuous dynamic changes exist in the snapdragon *S*-locus supergene, and the hyper-divergence may reveal the cause of reproductive isolation and speciation. These findings further enriched our understanding of the *S*-locus from a genome-wide perspective. Ultimately, our study provides unprecedented insights into the genome dynamics of this ornamental flower and offers abundant genomic resource for studies on *Antirrhinum*. A combination of advanced sequencing technologies, comparative genomics, and multi-omics datasets will help further decipher the genetic mechanisms of the *S*-locus evolution, thereby expediting the processes of horticulture and genetic research in the future.

## Materials and methods

### Sample collection and library construction

All plant samples were grown and collected from green house. For *de novo* assemblies, fresh young leaf tissues of 3 *Antirrhinum hispanicum* lines (*AhS_7_S_8_, AhS_7_Sc,* and *AhS_8_Sc)*, *Antirrhinum linkianum,* and *Misopates orontium* were collected for DNA extraction using cetyltrimethylammonium bromide (CTAB) method. PCR-free paired-end DNA libraries of ∼450bp insert size for each individual were prepared and then sequenced on an Illumina HiSeq4000 platform, following the manufacturer’s instructions. For PacBio Sequel II sequencing, high molecular weight (HMW) genomic DNA was extracted to construct SMRTbell libraries with insert size of 20kb, using SMRTbell Express Template Prep Kit 2.0 (PacBio, #100-938-900) and sequenced on PacBio Sequel II system. Another library of 15kb size was sequenced to generate high-qualitied consensus reads (HiFi reads) for *AhS_7_S_8_*. For BioNano optical maps sequencing, HMW genomic DNA of *AhS_7_S_8_* were extracted using BioNanoPrepTM Plant DNA Isolation Kit (BioNano Genomics), following BioNano Prep Plant Tissue DNA Isolation Base Protocol (part # 30068). Briefly, fresh leaves were collected and fixed with formaldehyde, then followed by DNA purification and extraction. Prepared nuclear DNA was labeled using BspQ1 as restriction enzyme with BioNano Prep Labeling Kit (Berry Genomics, Beijing). The fluorescently labelled DNA was stained at room temperature and then loaded onto a Saphyr chips to scan on the BioNano Genomics Saphyr System by the sequencing provider Berry Genomics Corporation (Beijing, China).

To obtain a chromosome-level genome, Hi-C library was constructed for *AhS_7_S_8_*. Young leaves were fixed with formaldehyde and lysed, then cross-linked DNA samples were digested with restriction enzyme DpnII overnight. After repairing sticky-ends and reversing the cross-linking, DNA fragments were purified, sheared, enriched and sequenced on Illumina NovaSeq6000 under paired-end mode. Four representative tissues, including leaf, petal, pollen, and stamen, of three species, plus pollen and pistil of the lines *AhS_7_S_C_* and *AhS_8_S_C_* were collected in the green house for RNA-seq. Two biological replicates were prepared for pistil and pollen. Strand-specific RNA-seq libraries were prepared following the manufacturer’s recommended standard protocol, and then sequenced on Illumina platform under paired-end mode. At least 20 million paired-end reads were generated for each library. Small RNAs were separated from total RNA described above using PAGE gel, then followed adapter ligation, reverse transcription, PCR product purification, and library quality control. Samples were sequenced using Nextseq500 (50 bp single-end) to yield 20 million reads per sample. Same tissues were also used for whole-genome bisulfite sequencing. Genomic DNA was extracted and fragmented into ∼450bp, then treated with bisulfite using the EZ DNA Methylation-Gold Kit according to manufacturer’s instructions. The libraries were sequenced on a HiSeq4000 system and at least 50 million PE-reads for each library were generated.

### Genome assembly and evaluation

To estimate the genome size independent of assembly, we characterized the basic genome feature using high-quality Illumina short reads. Firstly, PE reads were trimmed using Trim_galore (v0.6.1)^47^ with parameters “-q25 --stringency 3” to remove low-quality bases and adapters. Based on the 21-mer spectrum derived from clean data, heterozygosity rate and haploid genome length of three species were evaluated using software GenomeScope2^48^ .

For line *AhS_7_S_8_*, a consensus assembly consisting of collapsed sequences, and haplotype-resolved assemblies consisting of scaffolds from phased allele, were generated. Firstly, we removed reads of poor quality (RQ < 0.8) and short length (< 1kb) in PacBio subreads from the 20kb library before assembly. The contig-level assembly was performed on filtered subreads using Canu package (v1.9)^49^ with parameters “genomeSize=550m corMinCoverage=2 corOutCoverage=200 ‘batOptions = -dg 3 -db 3 -dr 1 -ca 500 -cp 50’”. After that, alternative haplotig sequences were removed using purge_dups^50^, only primary contigs were kept for further scaffolding. The draft genomes of *A .linkianum* and *M.orontium* were assembled using the same procedure. On the other hand, to obtain haplotype-resolved genome assemblies of *AhS_7_S_8_*, ∼40X PacBio HiFi reads along with Hi-C data were utilized to separate parental allele using FALCON-phase^51^. The resulting two phased contig-level haplotigs were used for downstream analysis independently. The contig assemblies were scaffolded into pseudo-chromosomes following the same pipeline described below (Fig. S1).

Low-quality BioNano optical molecular maps with length shorter than 150kb or label density less than 9 per 100kb were removed. After that, the BioNano genome maps combining with contigs were fed into BioNano Solve hybrid scaffolding pipeline v3.6 (BioNano genomics) to produce hybrid scaffolds under non-haplotype aware mode. When there were conflicts between the sequence maps and optical molecular maps, both were cut at the conflict site and assembled with parameters “-B2 -N2”. Followed by BioNano scaffolding, Hi-C data was utilized to cluster contigs into 8 groups subsequently. Clean Hi-C reads were then mapped to the BioNano-derived scaffolds using BWA^52^. The chromosome-level assembly was furtherly generated using the 3D-DNA analysis pipeline to correct mis-join, order, orientation, and translocation from the assembly^53^. Manual review and refinement of the candidate assembly was performed in Juicerbox Assembly Tools (v1.11.08)^54^ for interactive correction. Then the long pseudo-chromosomes were regenerated using script “run-asm-pipeline-post-review.sh -s finalize --sort-output --bulid-gapped-map” in 3D-DNA package with reviewed assembly file as input. Three chromosome-level assemblies of *AhS_7_S_8_* were constructed, and we named them as consensus-assembly SI-Ah (the mosaic one), S_7_ haplome and S_8_ haplome encompassing *S_7_* haplotype and *S_8_* haplotype, respectively). The sequences of two haplomes were named as Chr*.1 and Chr*.2 to distinguish homologous chromosomes. After manual curation, we used ultra-high-depth PacBio dataset of two offsprings and PBJelly^55^ to fill gaps in haplome assemblies, followed by two rounds of polish with gcpp program (https://github.com/PacificBiosciences/gcpp) in pbbioconda package. The organellar genome of *Antirrhinum* were identified using BLASTn against the *A.hispanicum* contigs with plant plastid sequences as reference (https://www.ncbi.nlm.nih.gov/genome/organelle/). The organellar genome sequences were submitted to CHLOROBOX^56^ website to annotate automatically and visualize genome maps.

The continuity and completeness of genome assemblies were benchmarked using long terminal repeat (LTR) Assembly Index (LAI)^16^ and BUSCO embryophyta_odb10^57^ core genesets, which include 2121 single-copy orthologous genes. Also, the PacBio reads and Illumina PE reads were mapped to respective assemblies using minimap2 and BWA^52^, to evaluate correctness. MUMmer package^58^ was used to inspect the difference between pair-wise genome assemblies. SNV (SNPs and InDels) were called using show-snps with parameters ‘-rlTHC’, then parsed and collapsed adjacent mutation/variation with our custom python script. While structural variation (SVs) including duplication, deletion, inversion, and translocation between two phased assemblies were called based on reciprocal alignment from nucmer and show-coords.

### Annotation of repetitive DNA elements

To identify repetitive elements within each genome sequences, firstly we used RepeatModeler (open-1.0.11) to build a *de novo* repetitive elements library from the assembled genome sequences independently. GenomeTools suite^59^ was used to annotate LTR-RTs with protein HMMs from the Pfam database. In addition, LTR_retriever^60^ was used to utilize output of GenomeTools and LTR_FINDER^61^ to generate whole-genome LTR annotation. Parameters of LTR_FINDER and LTRharvest were set requiring minimum and maximum LTR length of 100 bp and 7 kb, respectively.

Each of these libraries were classified with RepeatClassifier, followed by combining and removing redundancy using USERACH (https://www.drive5.com/usearch/). Then the unclassified sequences library was analyzed using BLASTX to search against transposase and plant protein databases to remove putative protein-coding genes. We submitted unknown repetitive sequences to CENSOR (https://www.girinst.org/censor/index.php) to get further classified and annotated. *De-novo* searches for miniatures inverted repeat transposable element (MITEs) used MITE_Hunter^62^ software with parameters “-w 2500 -n 5 -L 80 -I 80 -m 2 -S 12345678 -P 1”. Finally, all genome assemblies were soft-masked in company with the repeat library using RepeatMasker (open-4-0-9)^63^ with parameters “-div 22 -cutoff 220 - xsmall”.

### Gene prediction and functional annotation

The gene prediction was performed using BRAKER2^64^ annotation pipeline with parameters ‘--etpmode --softmasking’, which integrates RNA-seq datasets and evolutionary related protein sequences, as well as *ab initio* prediction results. Clean RNA-seq reads were aligned to the repeat-mask genomes using STAR^65^, then converted and merged to BAM format. The hint files were incorporated with *ab initio* prediction tool AUGUSTUS^66^ and GeneMark-ETP^67^ to train gene models. The predicted genes with protein length < 120 or TPM < 0.5 in any RNA-seq sample were removed. The *SLF* gene models included in the *S*-locus were examined in Genome browser and manually curated.

The tRNA genes were annotated by homologous searching against the whole genome using tRNAscan-SE(v2.0)^68^, rRNA and snRNA genes were identified using cmscan program in INFERNAL(v1.1.2)^69^ package to search from the Rfam database. Four small-RNA sequencing datasets were collapsed, combining with known miRNA sequence download from mirBase(v22)^70^ as input of miRDP2(v1.1.4)^71^ to identify miRNA genes across the genome. Next, miRNA mature sequences were extracted to predict potential target gene using miRANDA (http://www.microrna.org) by searching against all predicted full-length cDNA sequences.

To perform functional analysis of the predicted gene models, protein sequences were search against the InterPro consortium databases including PfamA, PROSITE, TIGRFAM, SMART, SuperFamily, and PRINTS as well as Gene Ontology database and pathways (KEGG, Reactome) databases using InterProScan (v5.51-85.0)^72^. The protein sequences were also submitted to plantTFDB online server^73^ to identify transcription factors and assign TFs families. In addition, predicted genes were also annotated biological function using diamond^74^ to search against NCBI non-redundant protein and UniProtKB (downloaded on 21^st^ Mar 2021, UniProt Consortium)^75^ database with an e-value of 1e-5. Genes were annotated to the eggnog database using eggNOG-mapper^76^.

### Whole-genome duplication and intergenomic analysis

Syntenic blocks (at least 5 genes per block) of *A.hispanicum* were identified using MCscan (Python version)^77^ with default parameters. Intra-genome syntenic relationships were visualized using Circos^78^ in Fig 1. We also compared *A.hispanicum* genome with several plant genomes, including *S.miltiorrhiza*, *S.lycopersicum*, *A.coerulea*, and *V.vinifera.* Dotplots for genome pairwise synteny was generated using the command ‘python -m jcvi.graphics.dotplot --cmap=Spectral --diverge=Spectral’.

For the *Ks* plots, we used wgd package^79^ with inflation factor of 1.5 and ML estimation times of 3 for reciprocal best hits search and MCL clustering. The synonymous substitution rate (*Ks*) for paralogs and orthologs were calculated using codeml program of PAML package^80^. We plotted output data from result files using custom python scripts with Gaussian mixture model function to fit the distribution (1-5 components) and determined peak *Ks* values. Estimated mean peak values were used for dating WGD events.

### Comparative genomics and divergence time estimation

Orthologous genes relationships were built based on the predicted proteomes deprived from consensus assembled *A.hispanicum* and other angiosperm species listed in (Table S11) using OrthoFinder2^81^. Only longest protein sequences were selected to represent each gene model. Rooted species tree inferred by OrthoFinder2 using STRIDE^82^ and STAG^83^ algorithm was used as a starting species tree for downstream analysis.

The species divergence time was estimated using MCMCtree in PAML^80^ package with branch lengths estimated by BASEML, with *Amborella* as outgroup. The Markov chain Monte Carlo (MCMC) process consists of 500,000 burn-in iterations and 400,000 sampling iterations. The same parameters were executed twice to confirm the results were robust. Species divergence time for *Amborella trichopoda-Arabidopsis thaliana* (173-199 Mya), *Vitis vinifera-Petunia axillaris* (111-131 Mya) and *Solanum lycopersicum– Solanum tuberosum* (5.23-9.40 Mya) which were obtained from TimeTree database^84^ were used to calibrate the estimation model, and constrained the root age <200 Mya.

To determine the gene family expansion and contraction, orthologous genes count table and phylogenetic species tree topology inferred by OrthoFinder2 were taken into latest CAFE5^85^, which employed a random birth and death model to determine expansion and contraction in gene families of given node. Corresponding p value was provided for each node and branch of phylogeny tree and cutoff 0.05 was used to identify gene families undergo expanded or contraction significantly at a specific lineage or species. KEGG and GO enrichment analysis of expanded gene family members were performed with Fisher’s exact test and adjusted by the Benjamini-Hochberge method to identify significantly enriched terms.

### Gene expression analysis

Apart from aiding gene model prediction, RNA-seq datasets were also used for transcripts quantification. For expression analysis, we used STAR^65^ to map clean RNA-seq reads to reference genomes with parameters ‘--quantMode TranscriptomeSAM --outSAMstrandField intronMotif --alignIntronMax 6000 -- alignIntronMin 50’. The transcripts per million (TPM) values were obtained using expectation maximization tool rsem^86^. The reproducibility of RNA-seq samples was assessed using spearman correlation coefficient. Samples from same tissue display strong correlation with a Pearson’s correlation coefficient of ***r*** > 0.85 (Fig. S29), indicating good reproducibility. Owing to haplotype-resolved assembly and the gene structure annotation of *A.hispanicum* genome, allelic genes from a same locus can be identified using a synteny-based strategy along with identity-based method. Reciprocal best hit between two haplotypes were identified using MCSCAN^77^ at first, and genes not in synteny block were searched against coding sequence counterparts to fill up the allelic relationship table. We used MUSCLE^87^ to align coding sequences of two allelic genes, and then calculated Levenshtein edit distance to measure allelic divergence. The divergence rate was defined as the number of edit distance divided by the total length of aligned bases.

Allelic transcripts quantification was conducted using cleaned RNA-seq datasets and allele-aware tool kallisto v.0.46.0^88^. We applied this software to obtain the expression levels read counts and TPM of genes of both haplotypes. Differential expression analysis of allelic genes was performed using R package edgeR^89^. Cutoff criteria of allele imbalanced expressed genes were set as adjusted *P* value < 0.01, false discovery rate < 0.01 and |log_2_(FC)|>1.

### Phylogenetic analyses of genes

F-box genes in whole genome were identified by searching Interproscan annotation results. Sequences annotated with both PF00646 (F-box domain) and PF01344 (Kelch motif) or PF08387 (FBD domain) or PF08268 (F-box associated domain) were considered as F-box genes. And the F-box genes located in the *S*-locus region of snapdragon were considered as potential *SLFs*. Organ specific protein and paralogs were selected using PF10950 as keyword, while PF02298 for plastocyanin gene. Protein sequence alignments were constructed using MUSCLE^87^ and manually checked. The maximum likelihood phylogenetic gene trees were constructed using raxml-ng^90^ with 100 replicates of bootstrap and parameter ‘-m MF’. And duplicate gene were classified into different categories using DupGen_finder^24^ with parameters ‘-d 15’.

### Cis-regulatory element analysis of predicted SLFs promoters

*Cis*-regulatory elements are specific DNA sequences that are located upstream of gene coding part and involved in regulation of genes expression by binding with transcription factors (TFs). We explored the upstream 2000bp sequences of 32 *SLFs* of *A.hispanicum* to discover TF binding sites using MEME^91^, and Tomtom was used for comparison against JASPAR database^92^ of the discovered motif.

### Whole-genome bisulfite sequencing analysis

The raw WGBS reads were processed to remove adapter sequences and low-quality bases using Trim_galore with default parameters. The cleaned reads were then mapped to the two haplomes using abismal command from MethPipe^93^ package with parameters ‘-m 0.02’. All reads were mapped to the chloroplast genome (normally unmethylated) of snapdragon to estimate bisulfite conversion rate. The non-conversion ratio of chloroplast genome Cs to Ts was considered ad a measure of error rate.

Each cytosine of sequencing depth ≥5 were seen as true *methylcytosines* sites. Methylation level at every single *methylcytosine* site was estimated as a probability based on the ratio of methylated to the total reads mapped to that loci. Methylation level in genes and 2kb flanking regions was determined using custom Python scripts. Gene body refers to the genomic region from start to stop codon coordinates in the gff file. Each gene and its flanking regions were partitioned into ten bins of equal size and average methylation level in each bin was calculated by dividing the reads indicating methylation by total reads observed in the respective bin.

### Data availability

All raw genome sequencing datasets, assembled genome sequences, and predicted gene models have been deposited at the GSA database^94^ in the National Genomics Data Center, Beijing Institute of Genomics (BIG), Chinese Academy of Sciences and China National Center for Bioinformation, under accession numbers <u>PRJCA006918</u> that are publicly accessible at http://ngdc.cncb.ac.cn/gsa. RNA-seq datasets have been deposited under accession id <u>CRA005238</u>.

## Acknowledgements

This work was supported by the National Natural Science Foundation of China (32030007) and the Strategic Priority Research Program of the Chinese Academy of Sciences (XDB27010302).

## Author contributions

Y.X. designed and supervised the research; Y.E.Z. and Q.Q.H performed the experiments and prepared plant samples for sequencing; L.C. provided plant seeds; S.Z. analyzed the data; D.Z. contributed data visualization; S.Z., Y.X. and E.C. wrote the manuscript. All authors discussed the results and commented on the final manuscript.

