## Supplementary material for "The snapdragon genomes reveal the evolutionary dynamics of the *S* locus supergene": Fig. S

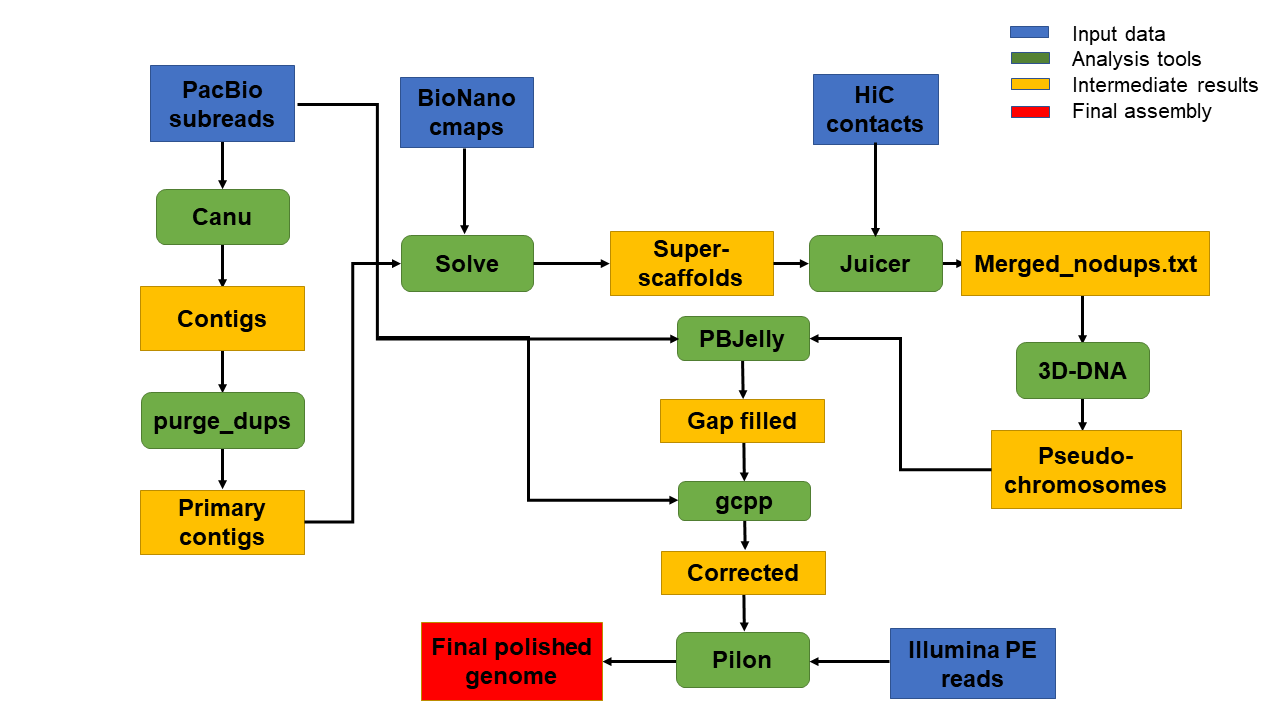

**Fig. S1** Analysis flowchart of heterozygous *AhS_7_S_8_*.

| 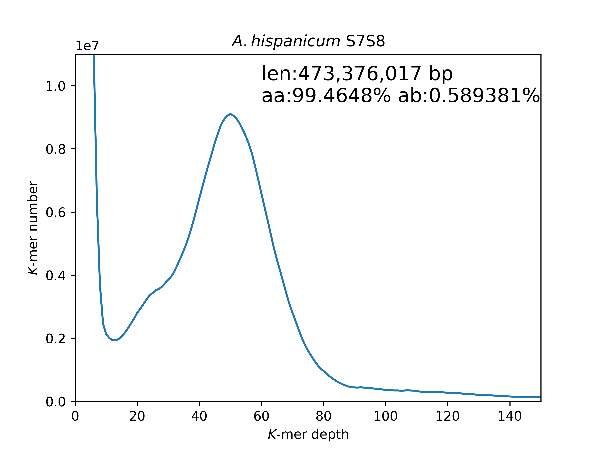 | 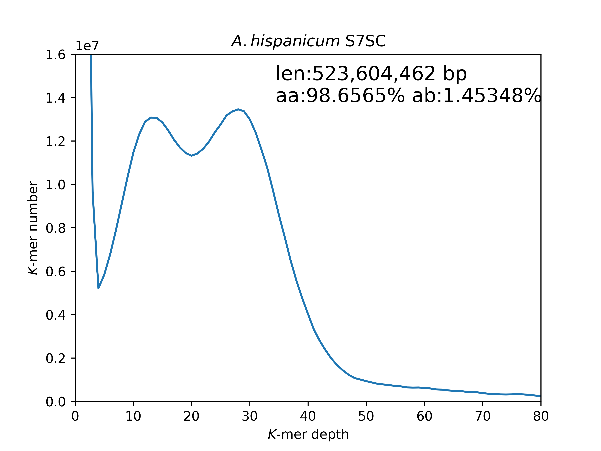 |
| --- | --- |
| 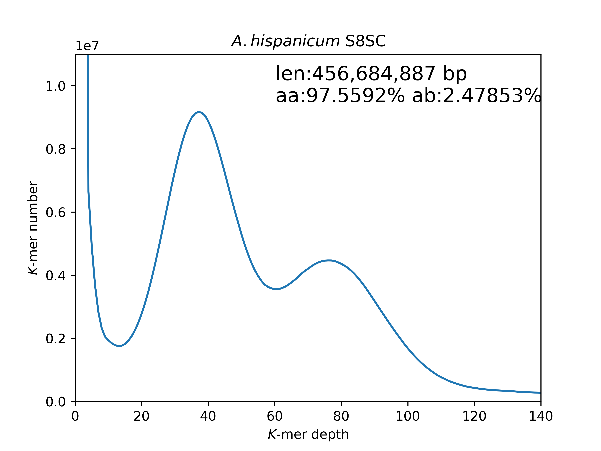 | 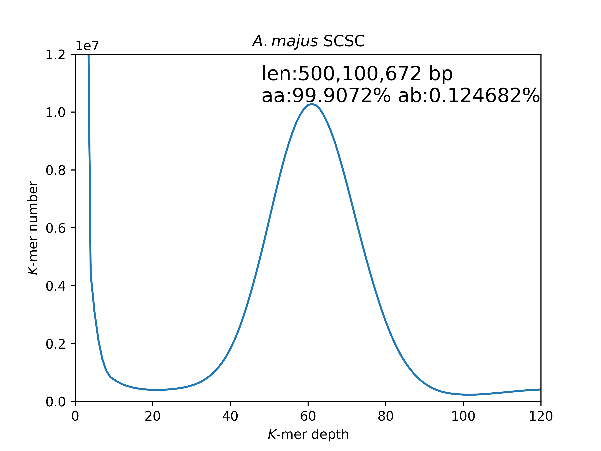 |
| 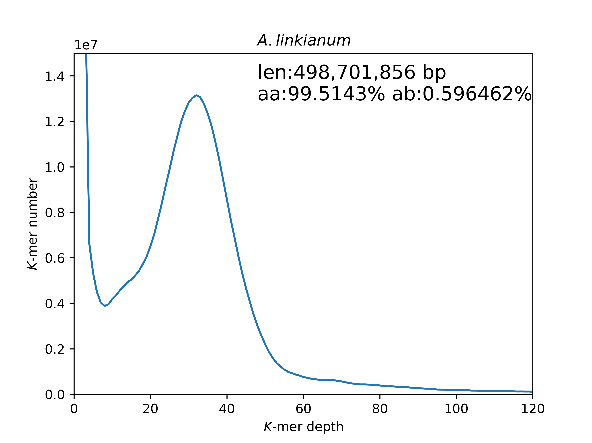 | 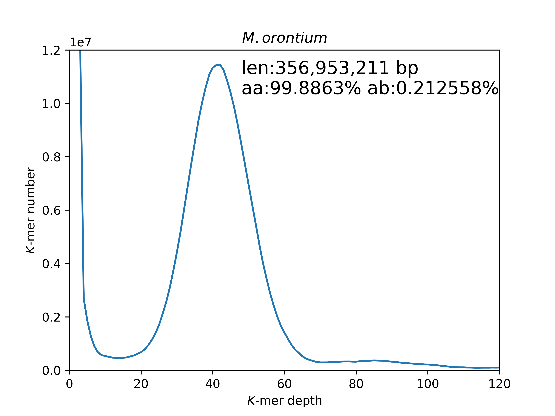 |

**Fig. S2** Estimation of genome size using k-mer spectrum analysis. The frequency histograms of 21-mers were generated from whole genome Illumina reads of 3 *A.hispanicum* lines (*AhS_7_S_8_*, *AhS_7_S_C_*, *AhS_8_S_C_*), self-compaptible *A.majus* (*AmS_C_S_C_*), *A.linkianum* and *Misopates orontium*. The first and second peak represent the heterozygous and homozygous regions in genome respectively.

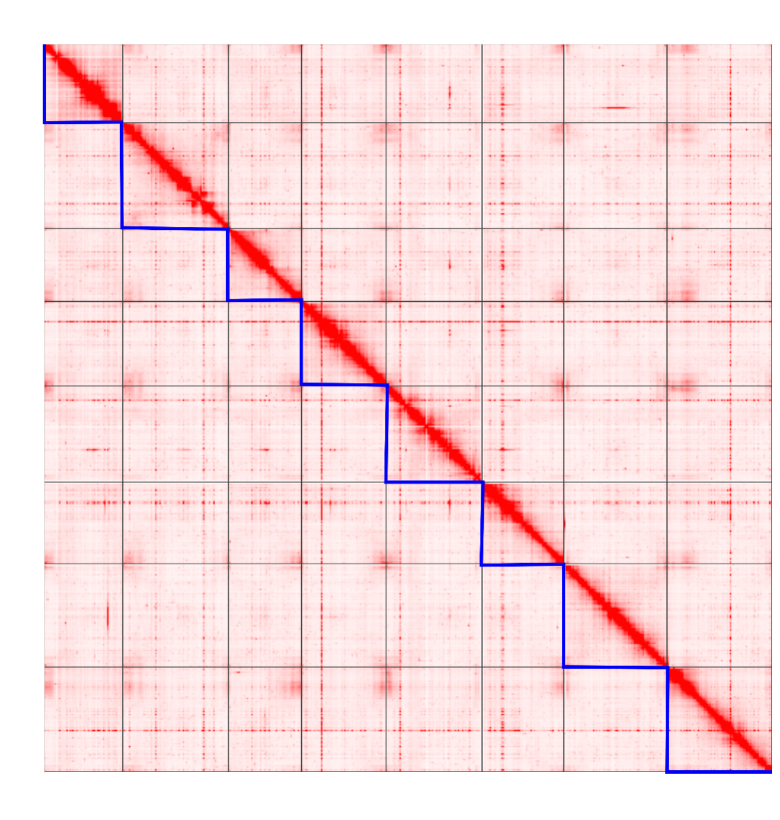

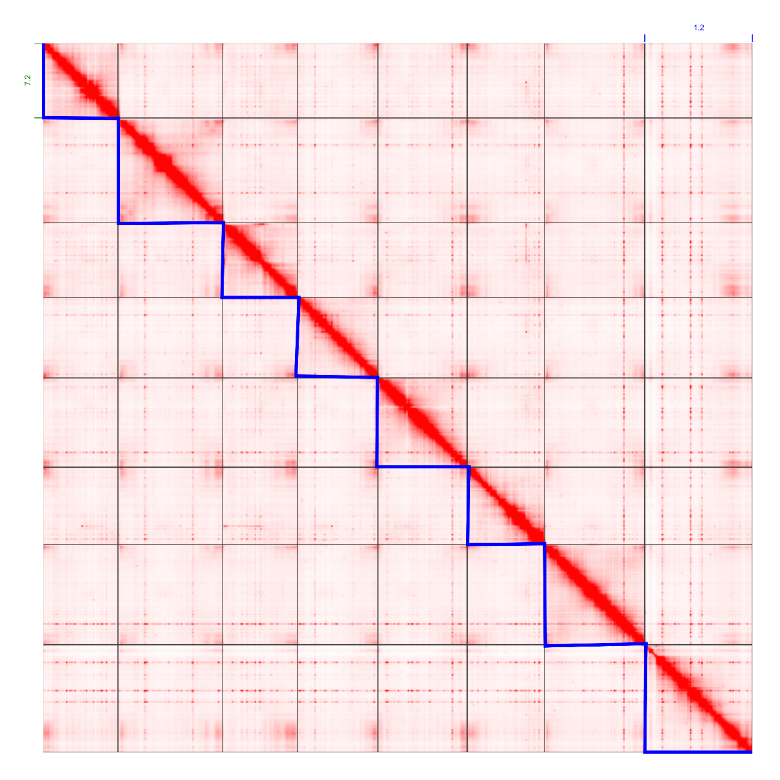

**Fig. S3** Hi-C contact map representing chromosome-scale phased genomes of *Antirrhinum hispanicum*. Hi-C contact matrix visualization for the phased genomes, S_7_ haplome and S_8_ haplome. The pixel intensity represents the Hi-C links at 100 Kb size-windows at logarithmic scale. The blue bold lines separate chromosomes for each phased genome.

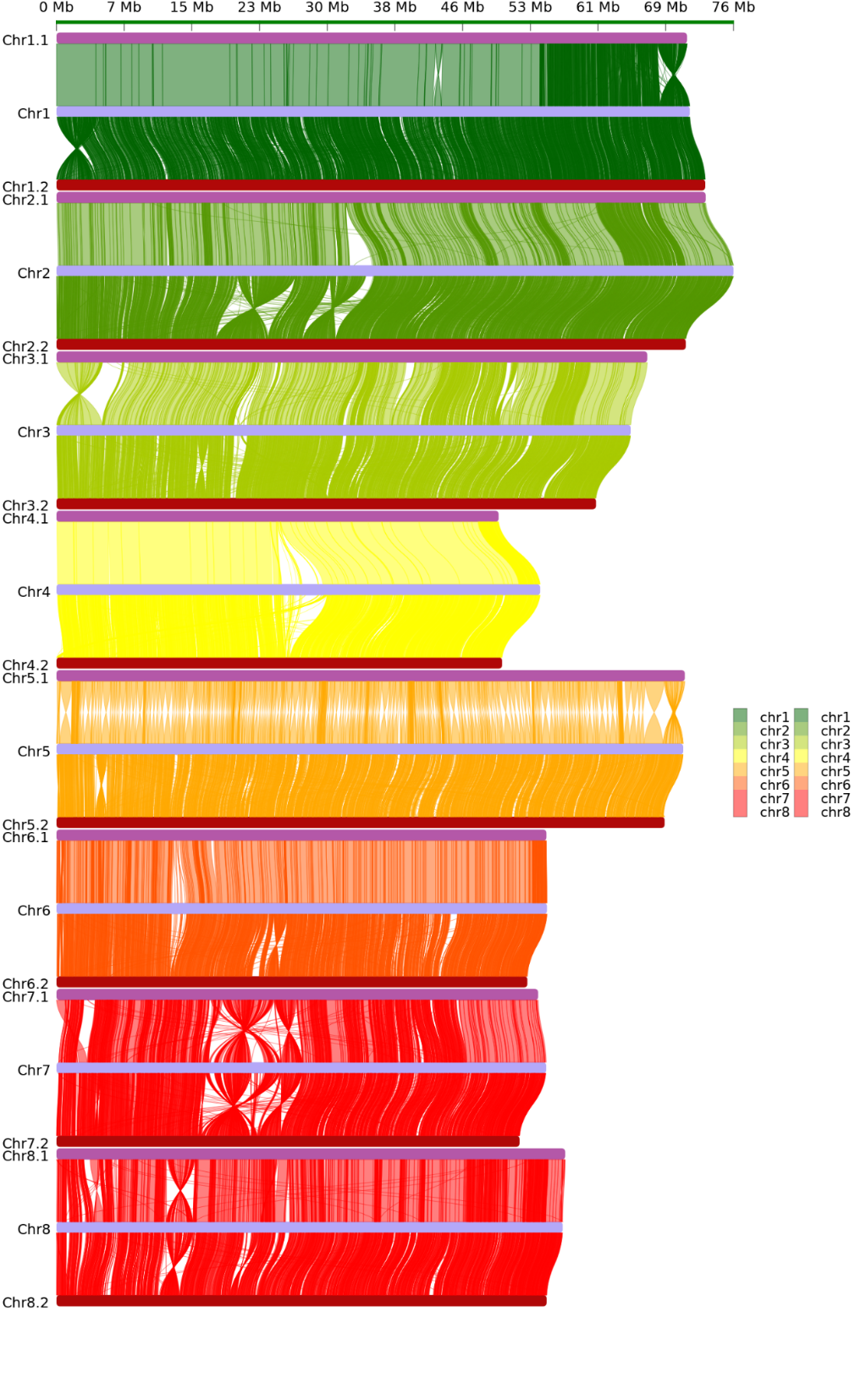

**Fig. S4** DNA colinear alignment between two haplomes and consensus assembly of *A.hispanicum*.

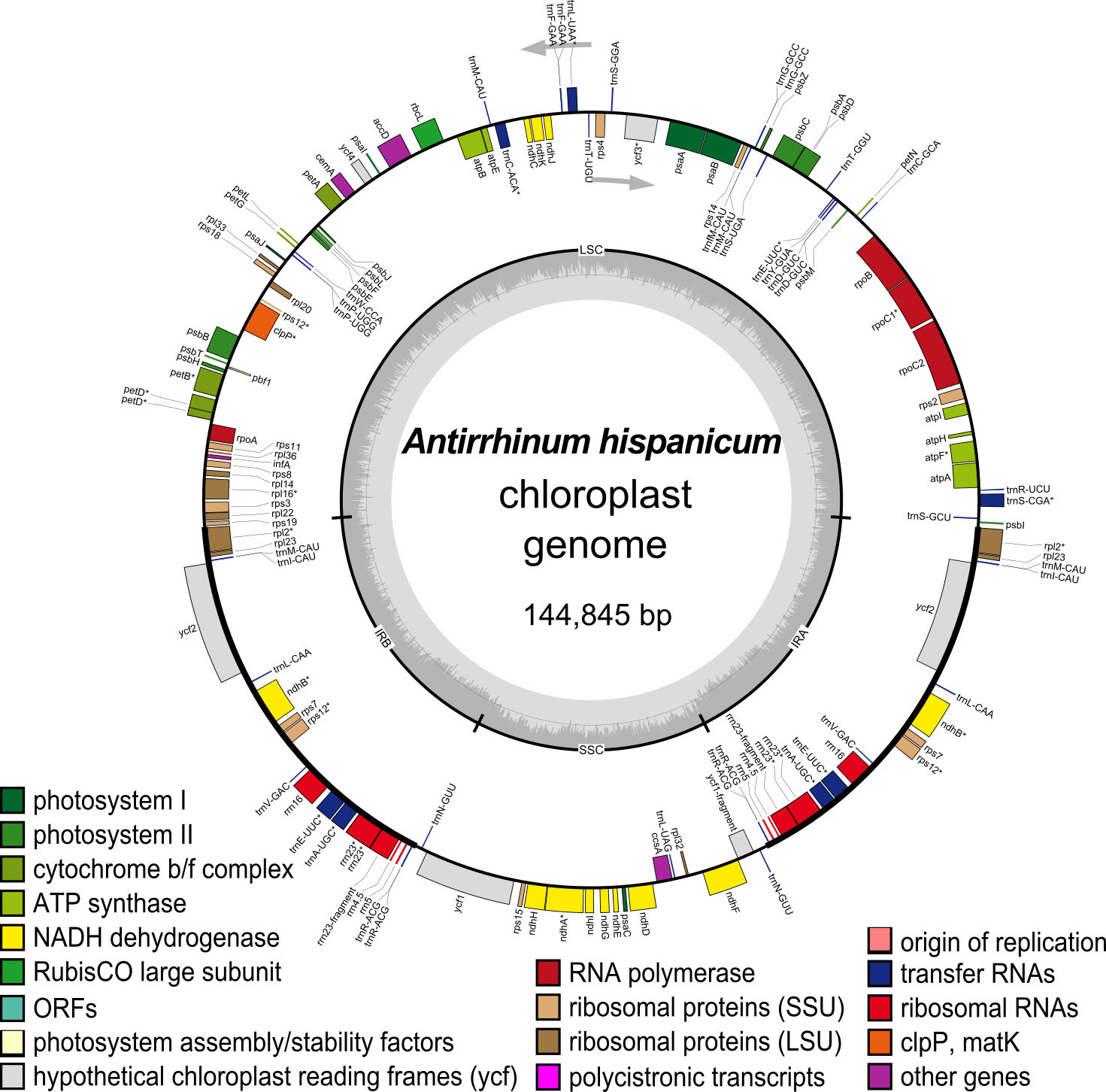

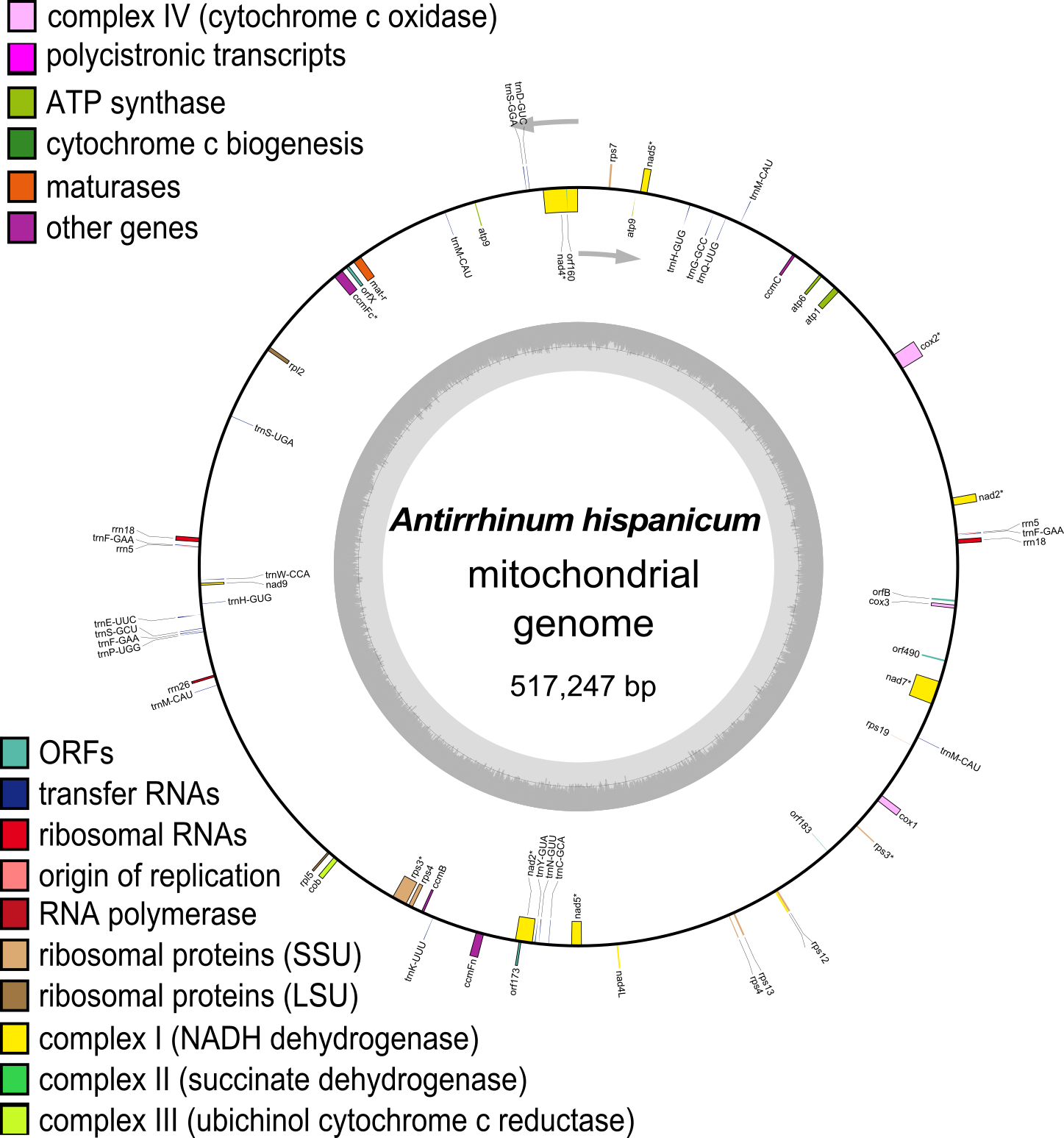

**Fig. S5** The circular map of *A.hispanicum* chloroplast genome and mitochondrial genome. The legend describes the family of genes in the genome. The inner circle depicts nucleotide content (GCcontent in dark gray and ATcontent in light gray). Inverted repeats (IRA and IRB) along with large (LSC) and small (SSC) single copy regions are indicated on the inner circle. The outer circle shows annotated genes that are color-coded by function. Genes located on the outside of the circle are transcribed in a clockwise direction, and those on the inside of the circle are transcribed counterclockwise.

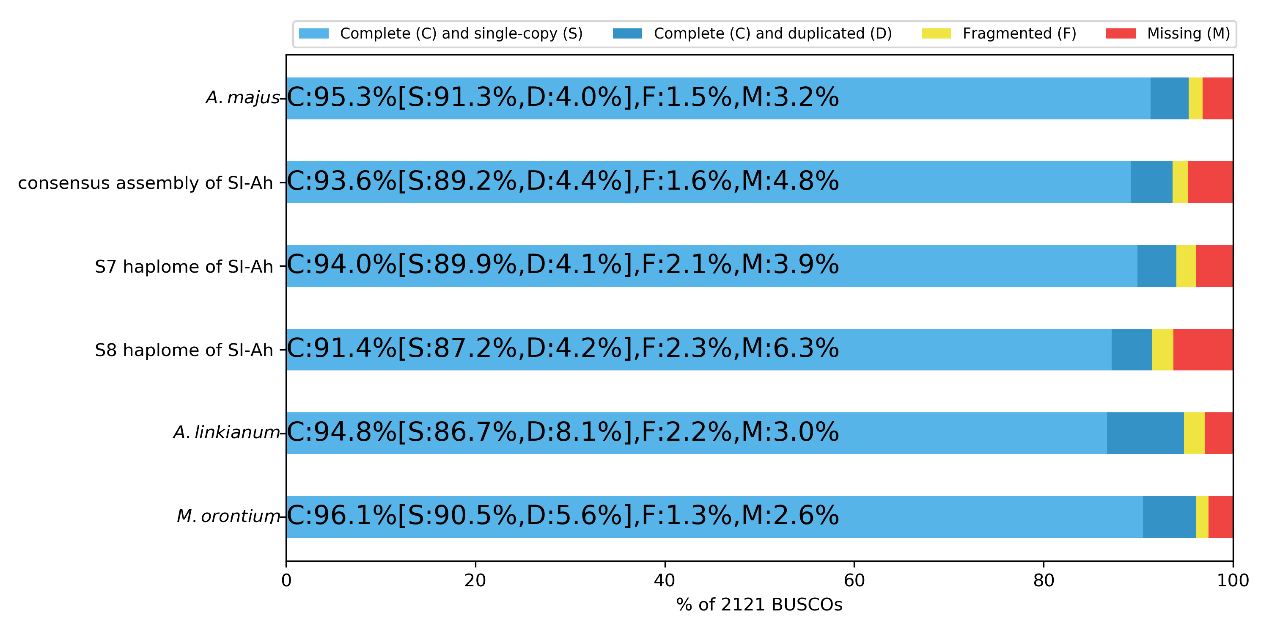

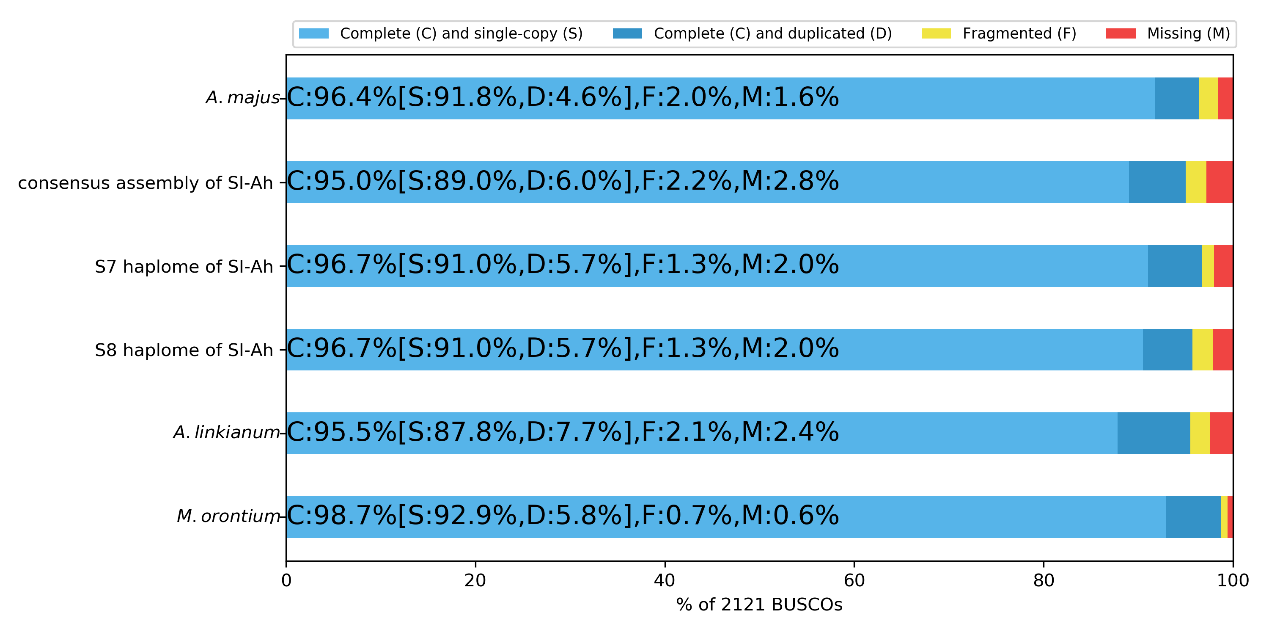
 **Fig. S6** BUSCO assessment results for *A.majus*, consensus-assembly and two haplomes of SI-Ah, *A.linkianum*, *M.orontium*. The different panels correspond to different running modes:(A) assembled genomes, (B) predicted proteins.

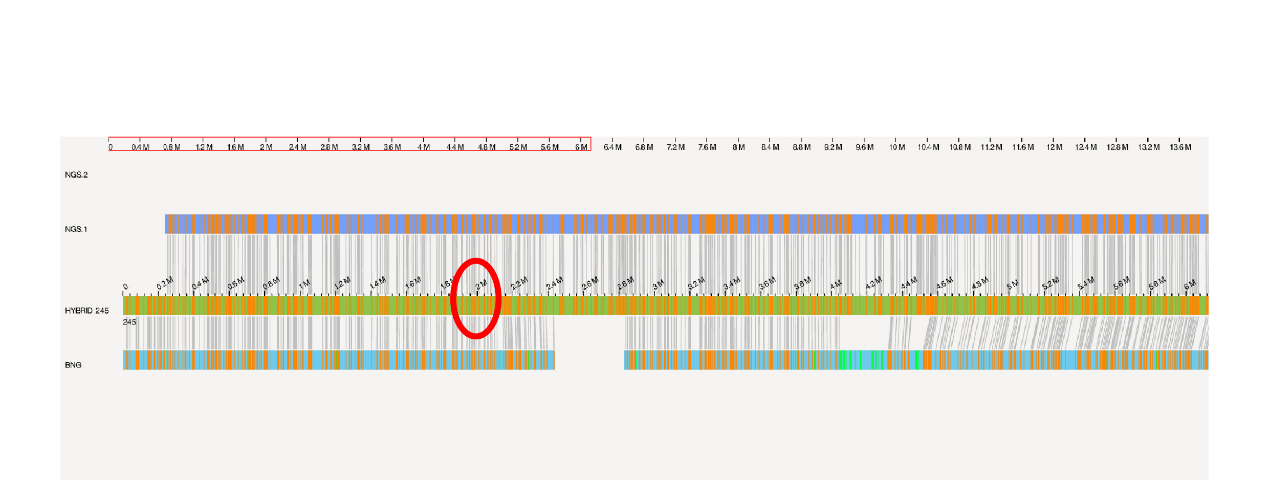

**Fig. S7** BioNano evidence for *S*-locus correctness of S_7_ haplotype. The top blue panel represents pacbio-assembly reference cmap. The middle panel represents hybrid scaffold consensus cmap. The bottom represents BioNano *de novo* assembly cmap. The red circle denoted *S-RNase*. The grey lines between bars represent consensus restriction enzyme cutting sites.

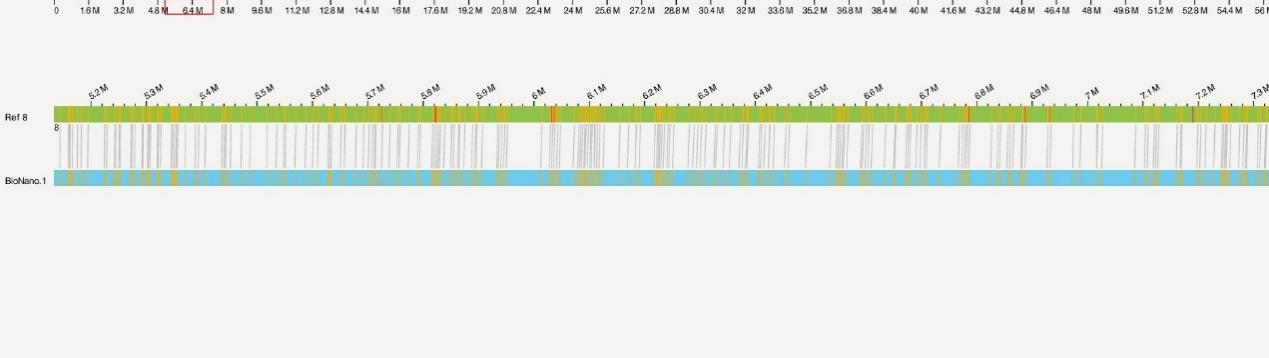

**Fig. S8** Bionano evidence for pseudo *S*-locus correctness of SC haplotype in *A.majus*. The top green panel represents reference cmap. The bottom represents bionano *de novo* assembly cmap. The grey lines between bars represent consensus restriction enzyme cutting sites.

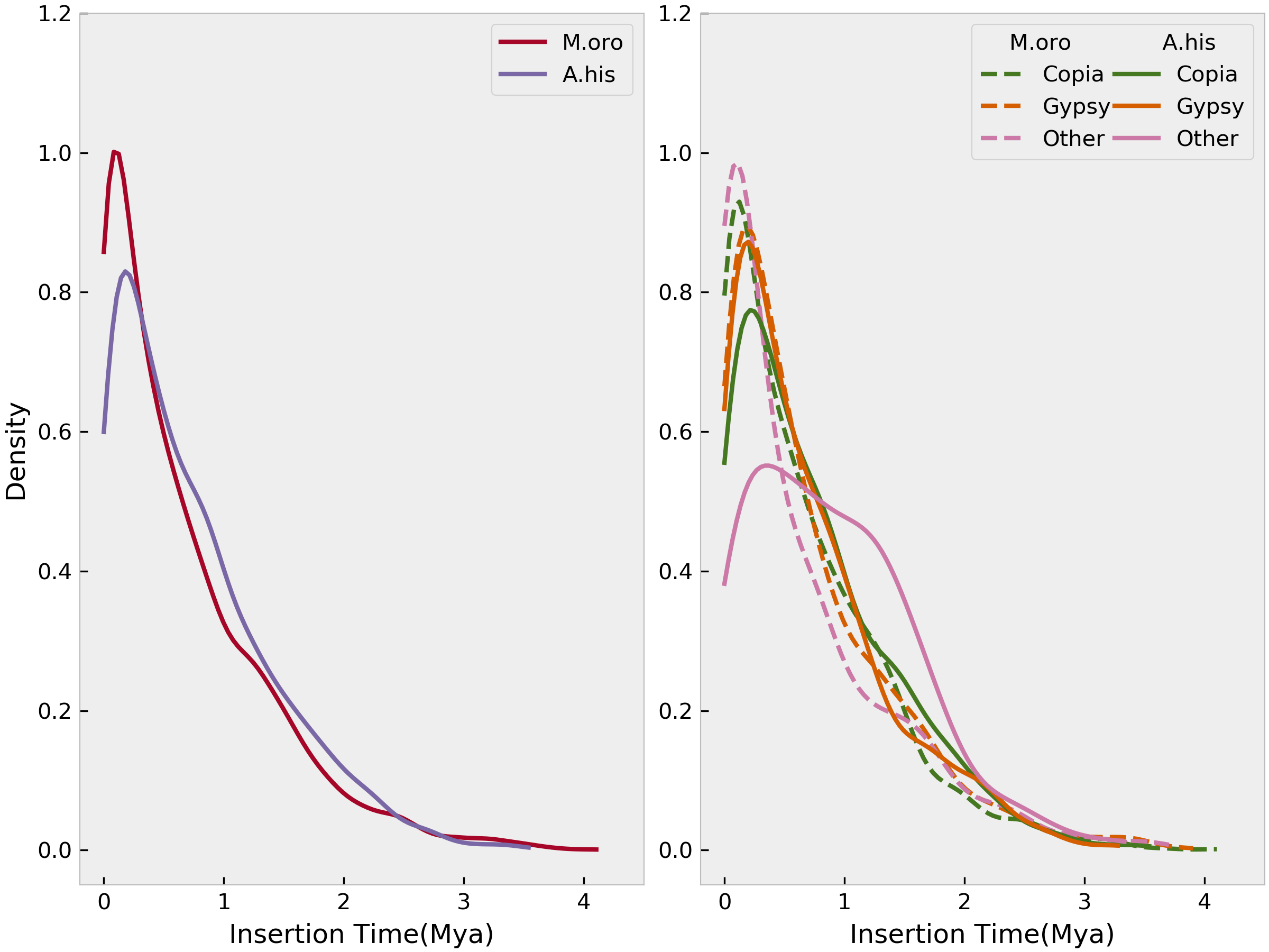

**Fig. S9** The density distribution of intact LTR-RTs insertion time. a) The density distribution of insertion time of intact LTR-RTs in self-compatible *M.orontium* and self-incompatible *A.hispanicum* genome. b) The density distribution of insertion time of intact Copia, Gypsy and other unclassified LTR-RTs in *M.orontium* and *A.hispanicum* genomes.

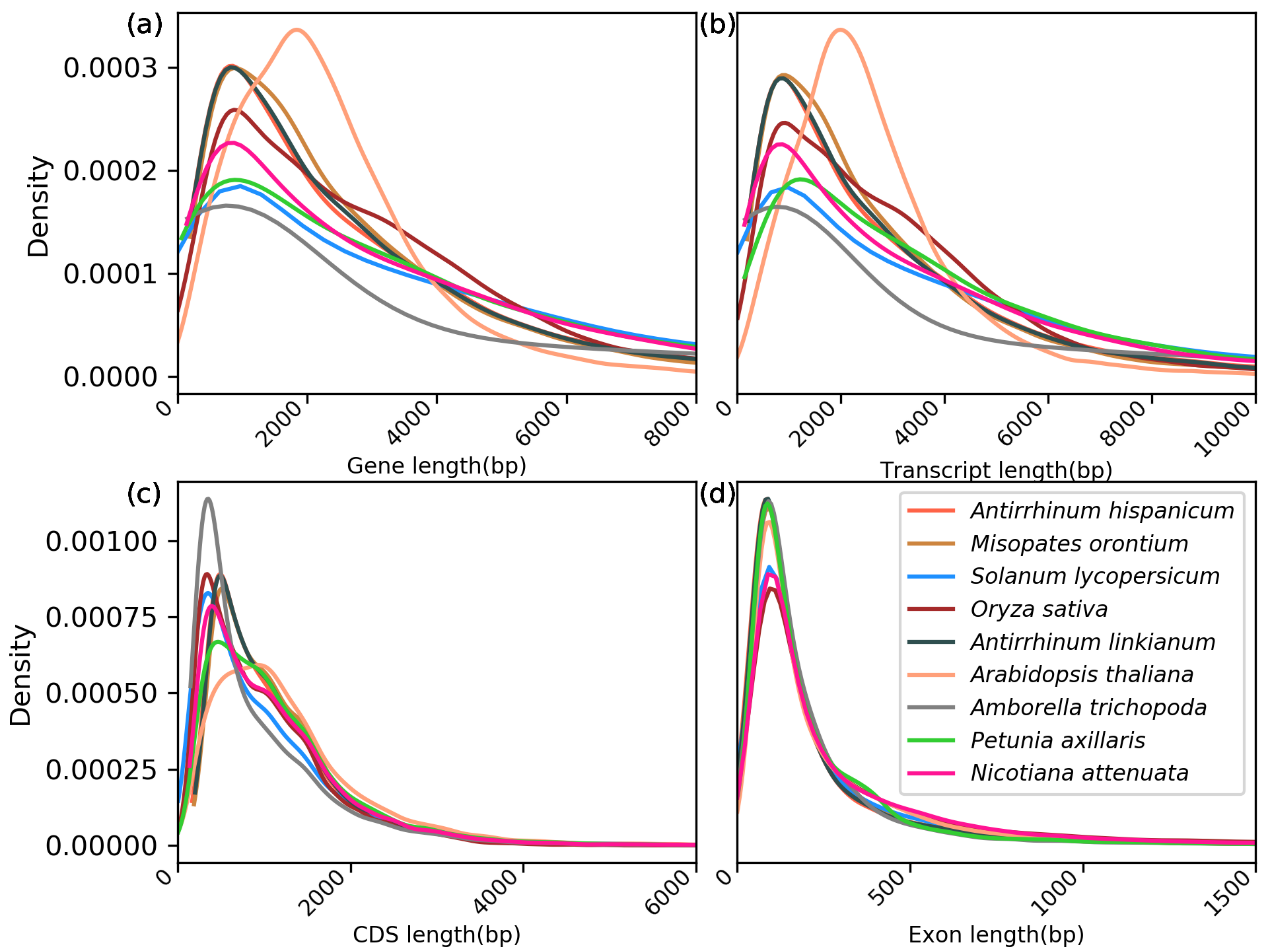

**Fig. S10** Comparison of gene length (a), transcript length (b), CDS length (c), and exon length (d) of *A.hispanicum*, *A.linkianum*, *M.orontium* and other six species. The x-axis represents length and the y-axis represents the density of genes. No unexpected differences were observed between *A.hispanicum* and *A.linkianum*. Other species, except *A.thaliana*, show similar genomic feature with snapdragon.

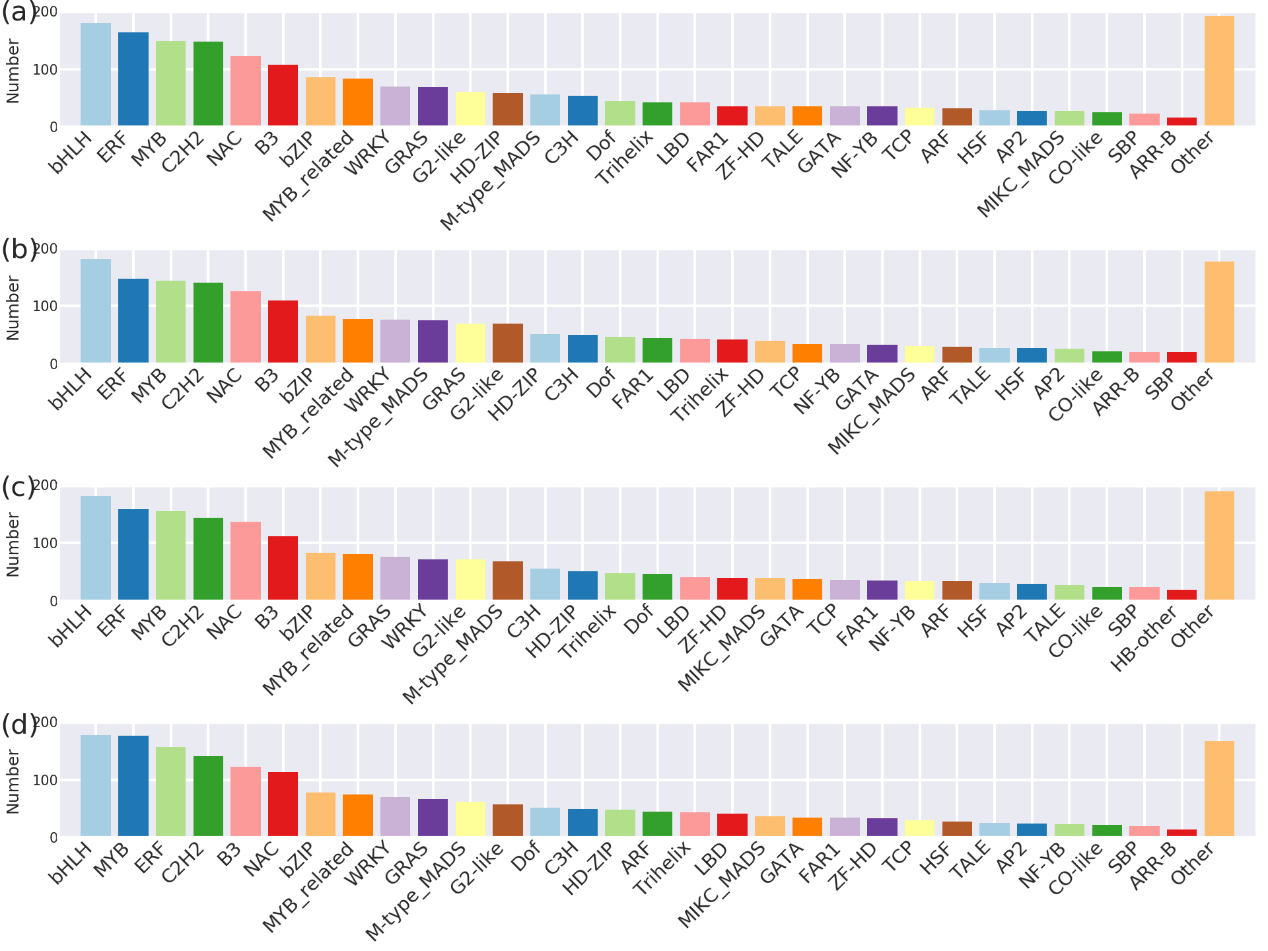

**Fig. S11** Families distribution of predicted transcription factors in *A.hispanicum* (a), *A.linkianum* (b), *A.majus*(c) and *M.orontium*(d) genome.

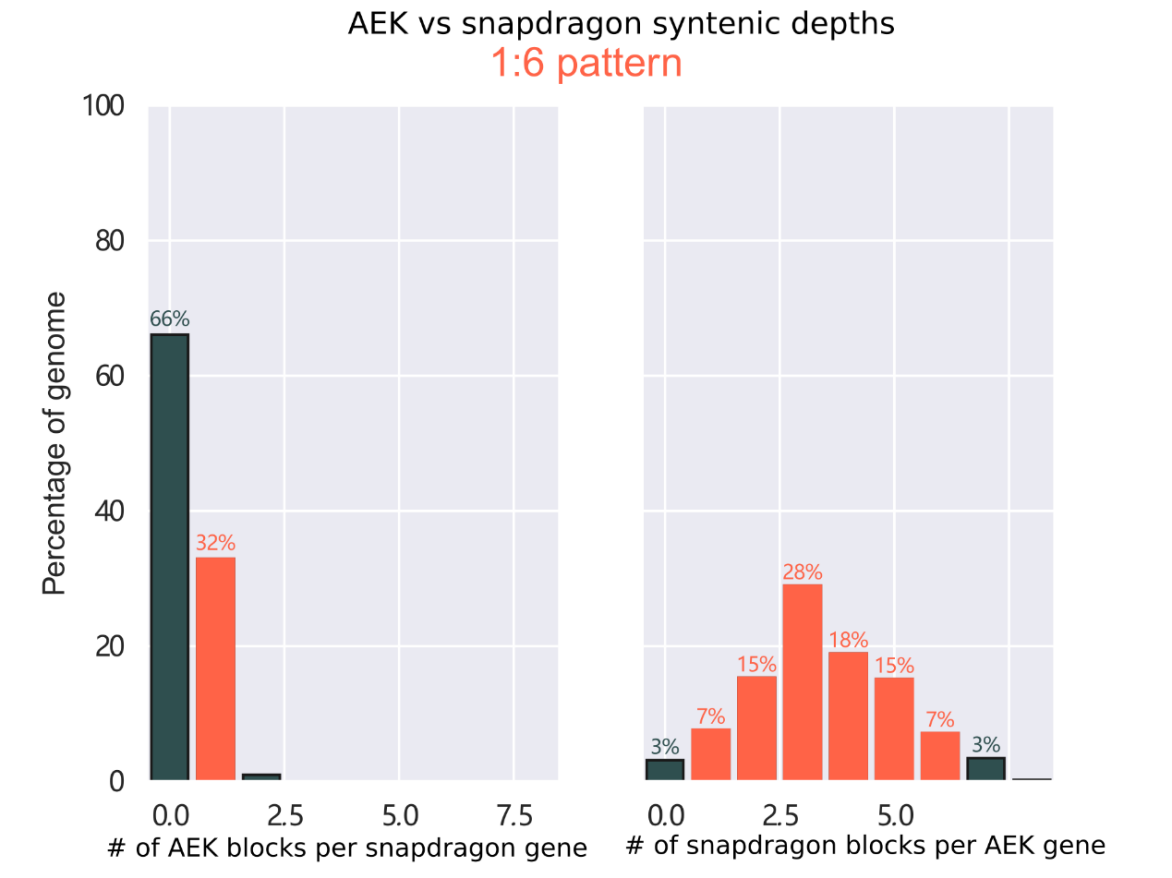

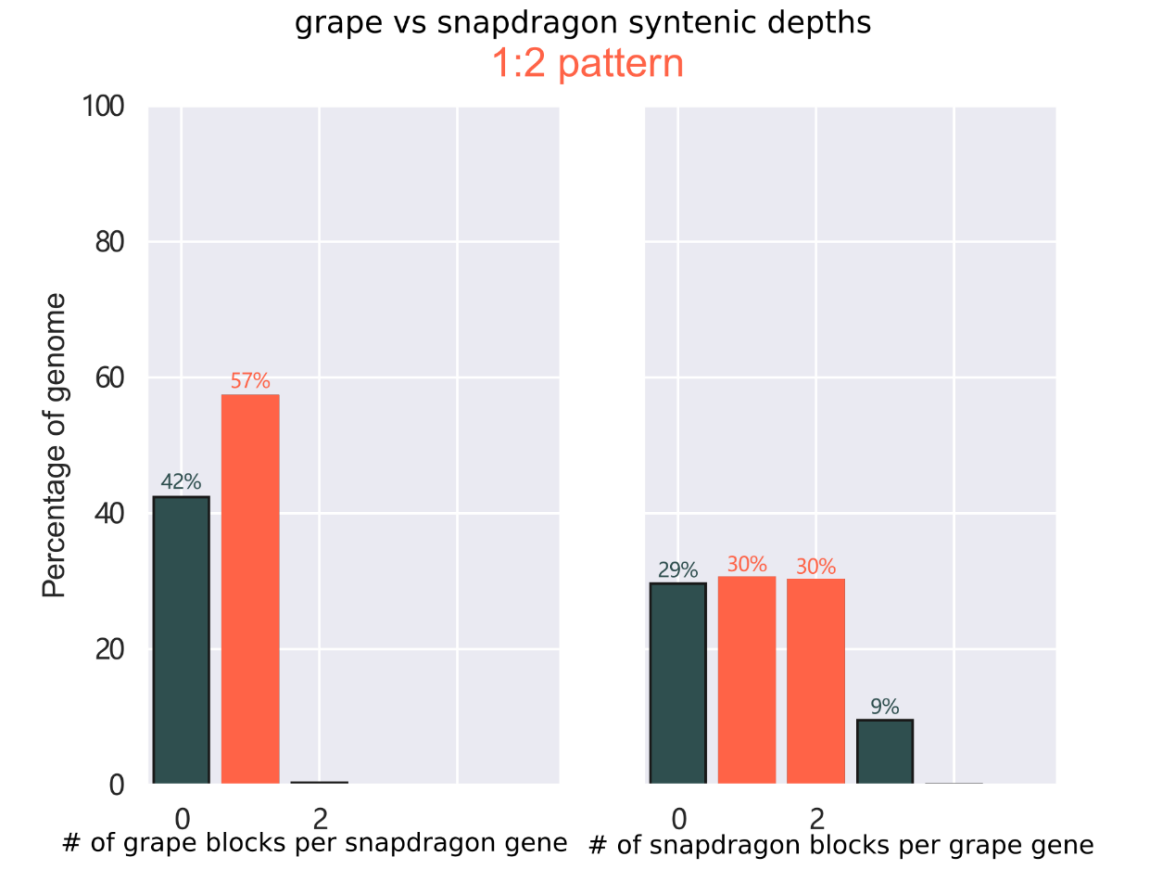

**Fig. S12** Ratio of syntenic depth between snapdragon and AEK, grape genome respectively. Syntenic blocks of snapdragon per AEK gene and syntenic blocks of AEK per snapdragon gene are shown indicating a clear 1:6 pattern.

| 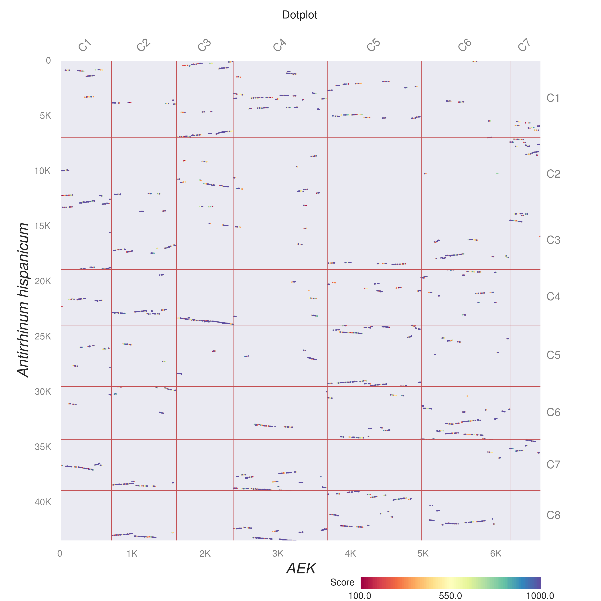 | 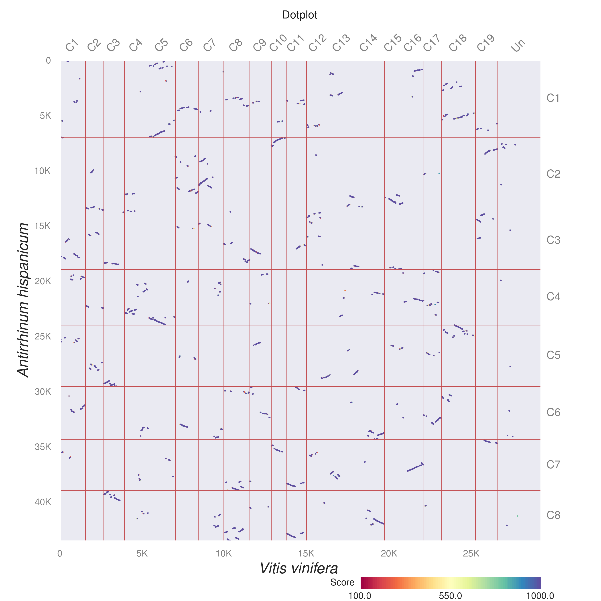 |
| --- | --- |
| 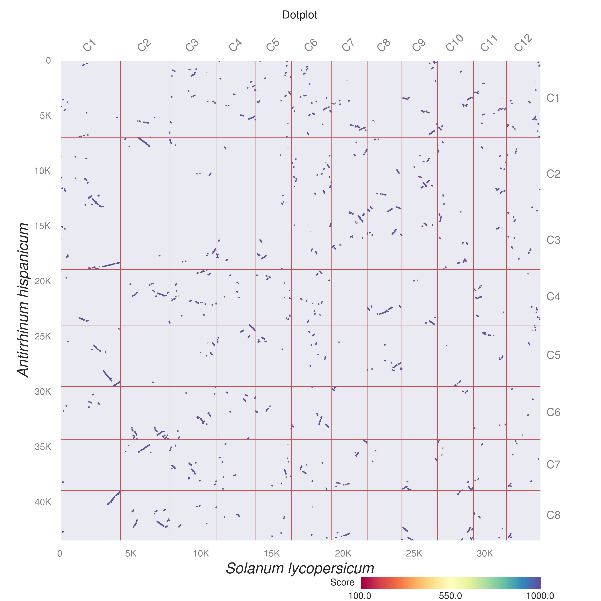 | 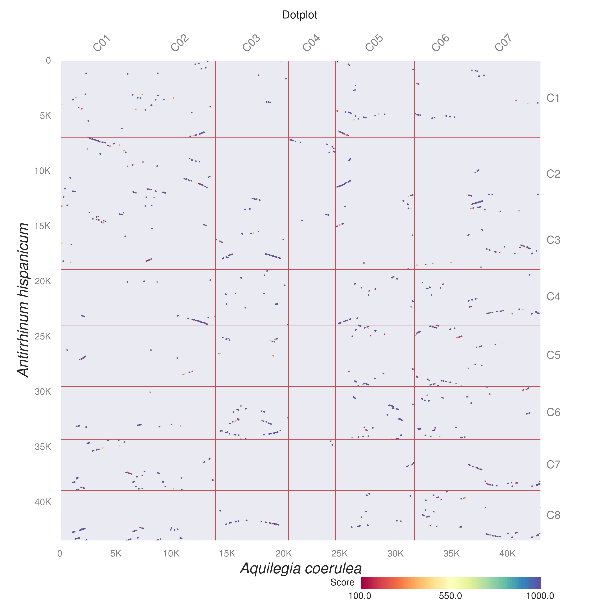 |
| 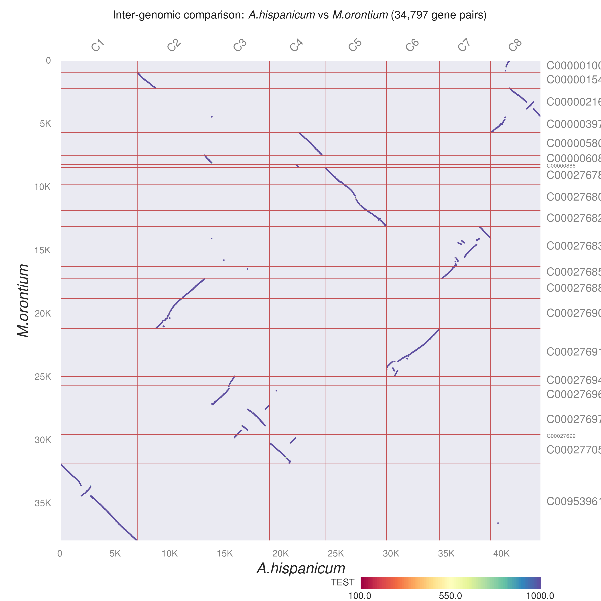 |  |

**Fig. S13** Inter- and Intra- genomic dot-plot for *Antirrhinum hispanicum* genome and other selected species including ancestral eudicot karyotype (AEK). Dot-plot based synteny visualization showed a clear 1:6 synteny depth pattern between AEK and *A.hispanicum* genome. Synteny plot between, *A.hispanicum* - *Vitis vinifera*, 1:2 pattern, *A.hispanicum - Solanum lycopersicum*, 2:3 pattern, *A.hispanicum - Aquilegia coerulea*, 1:3 pattern. Previous studies demonstrated that Aquilegia species experienced a WGD event after the divergence with grape.

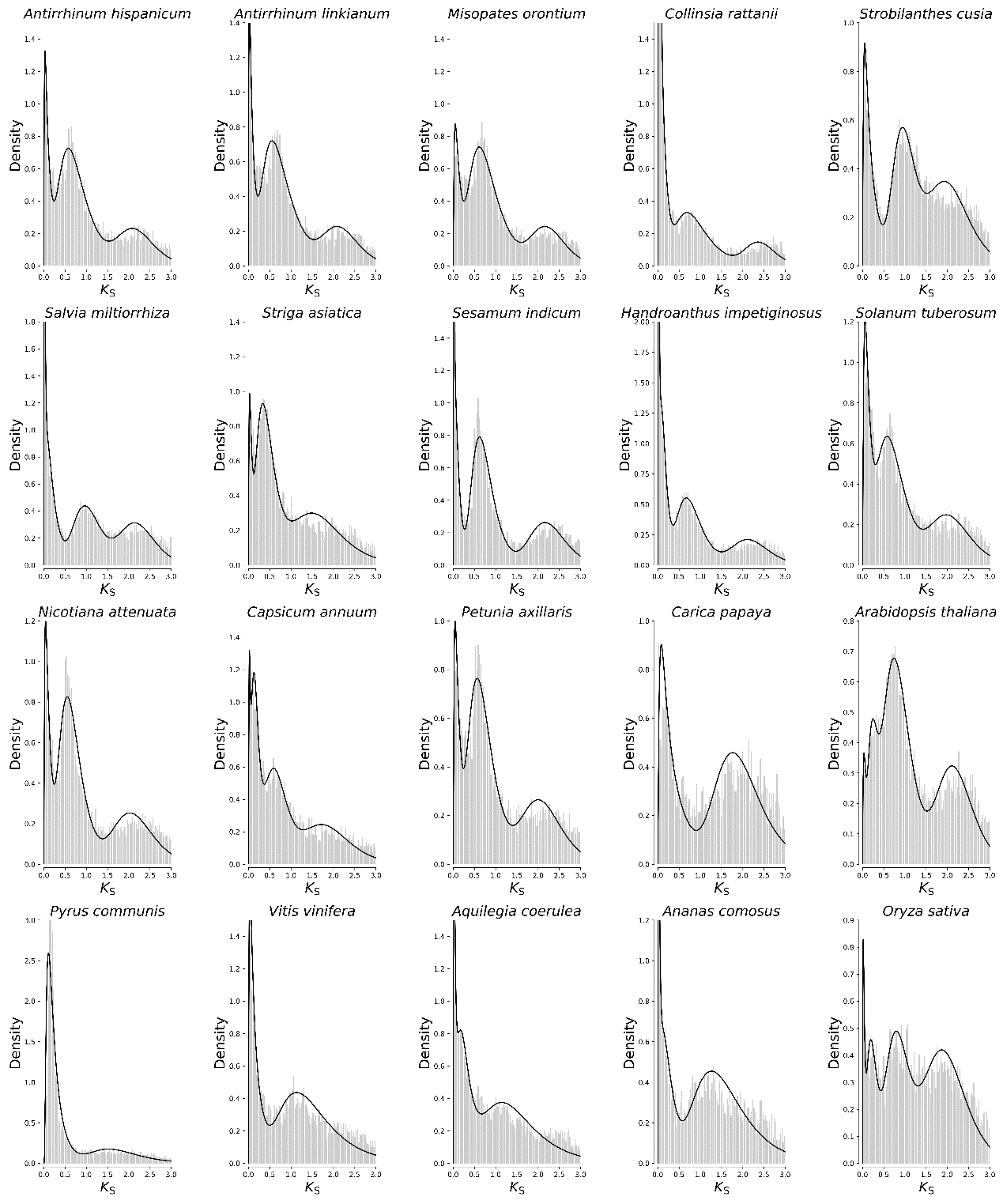

**Fig. S14** Density plot of *Ks* (synonymous substitution rate) distribution of paralogous gene pairs for selected plant species. The solid lines were fitted using Gaussian Mixture Model function.

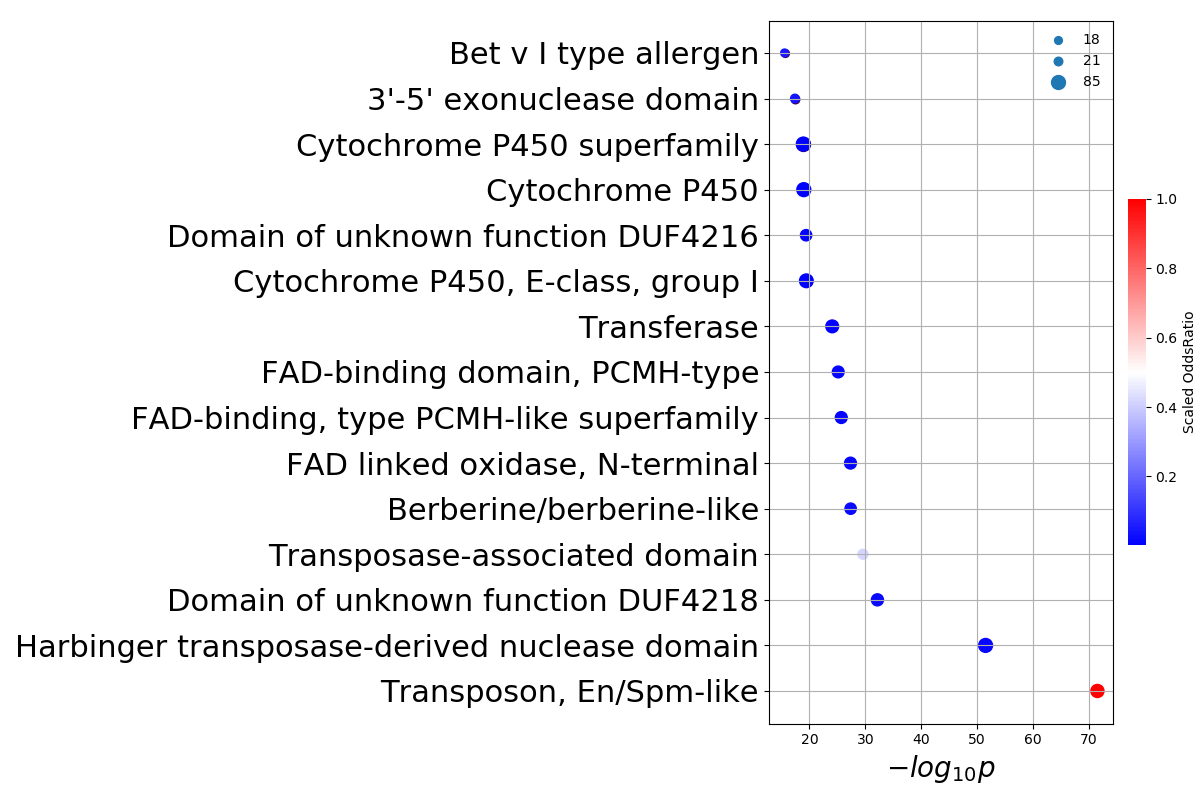

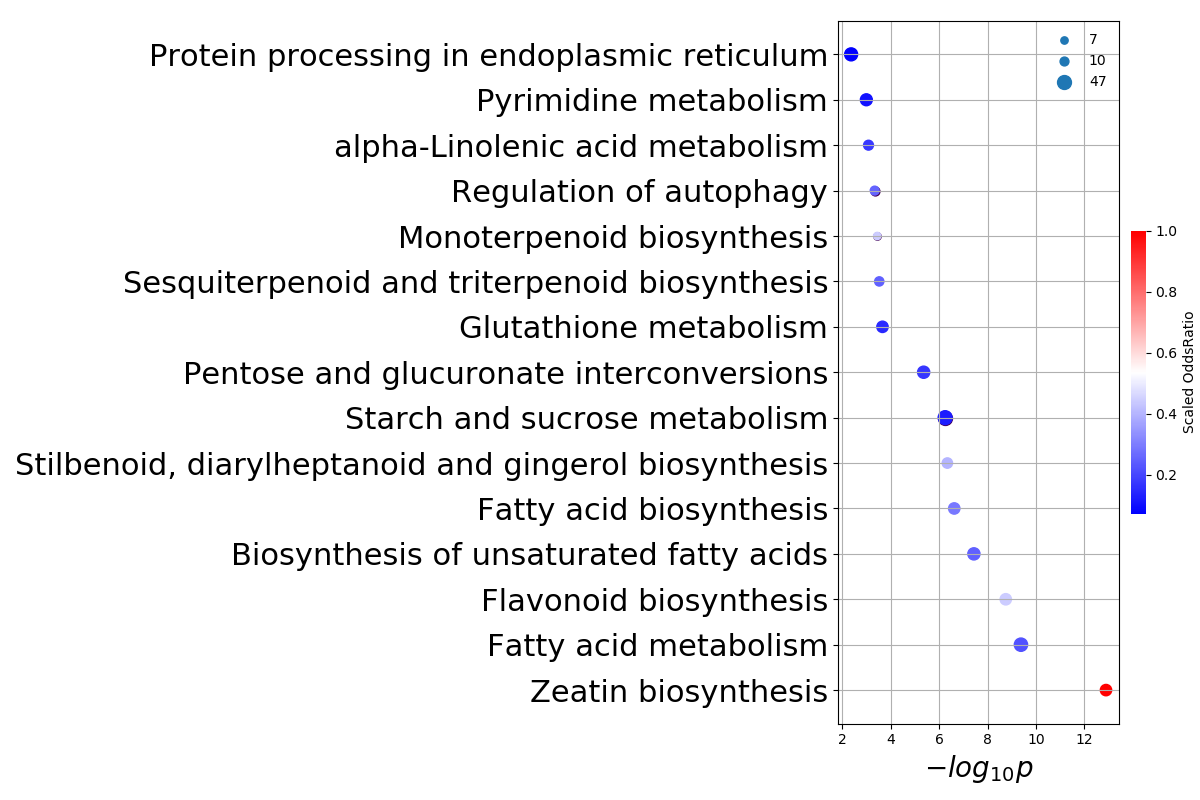

**Fig. S15** InterPro, and Kyoto Encyclopedia of Genes and Genomes (KEGG) enrichment analysis of *Antirrhinum* lineage gene members from expanded families. The color of circle represents the scaled odds ratio. The size of circle represents the gene number of the IPR terms or KEGG pathway. The p-value is corrected in hypergeometric test.

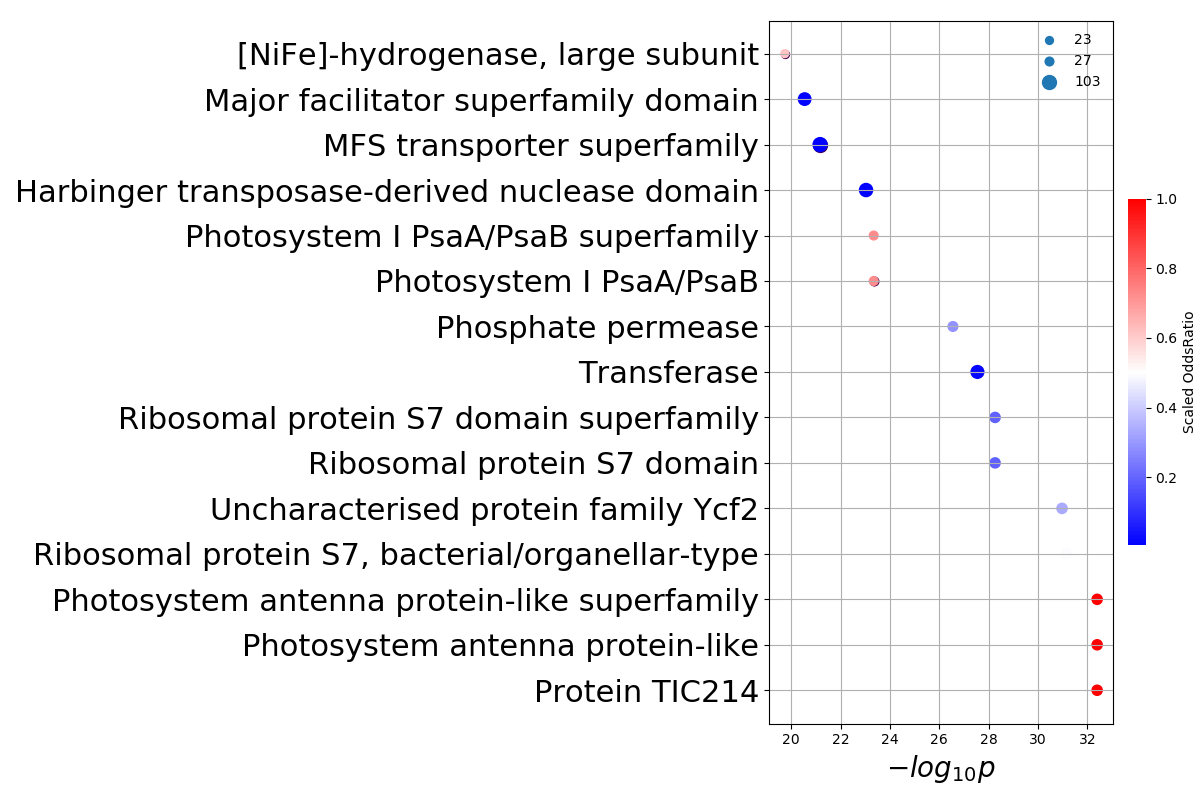

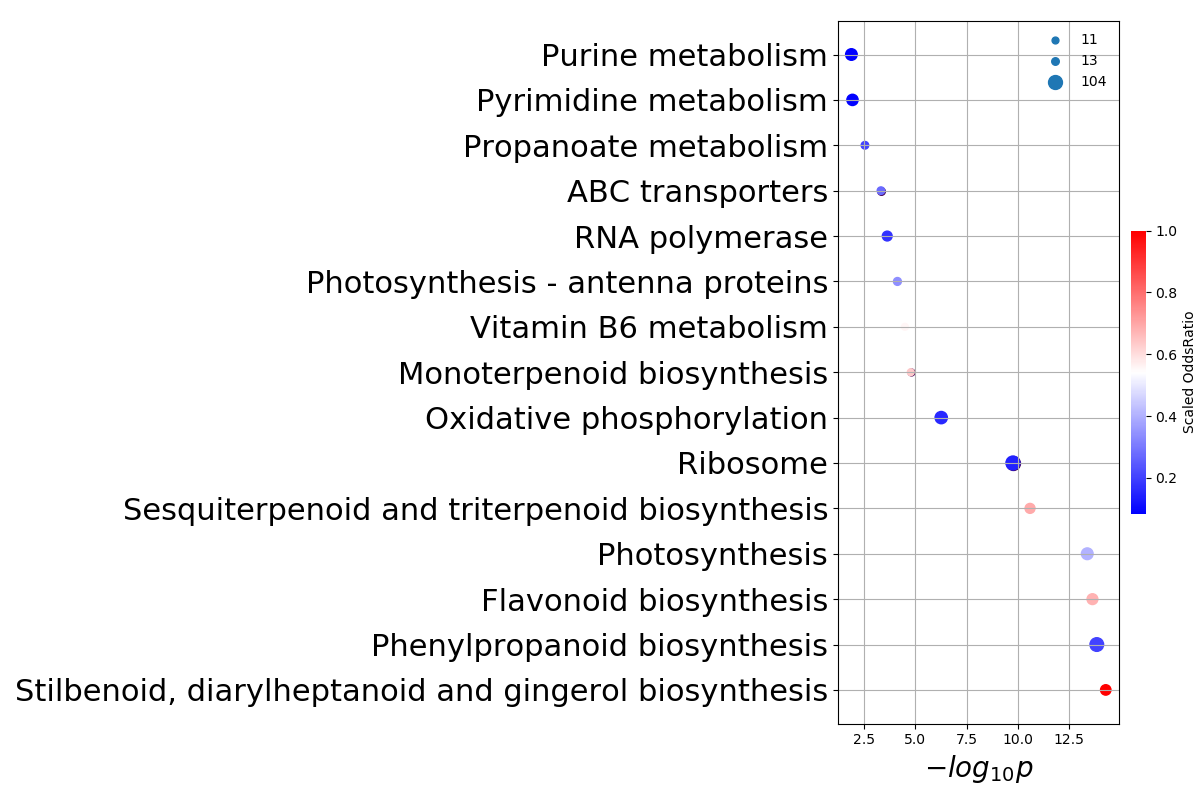

**Fig. S16** InterPro, and Kyoto Encyclopedia of Genes and Genomes (KEGG) enrichment analysis of *Misopates orontium* gene members from expanded families. The color of circle represents the scaled odds ratio. The size of circle represents the gene number of the IPR terms or KEGG pathway. The p-value is corrected in hypergeometric test.

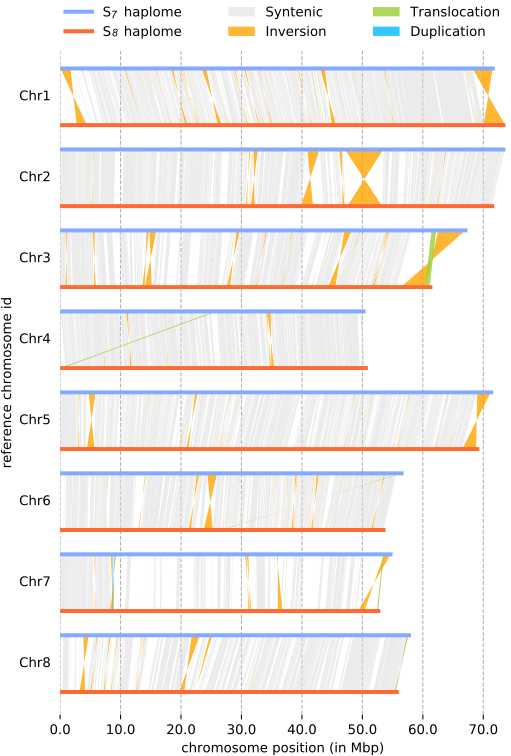

**Fig. S17** DNA alignment of two haploid genomes. Structural rearrangements are represented by colored lines (orange for inversions, green for translocations, blue for duplications) connecting regions of two homologous chromosomes (blue and red horizontal lines, respectively); syntenic regions are connected by gray lines.

| 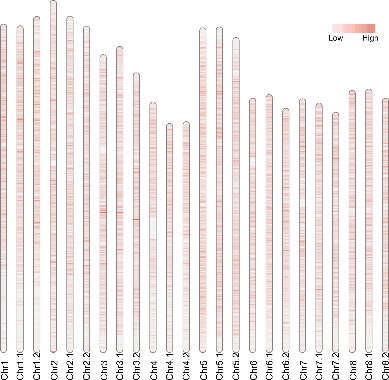 | 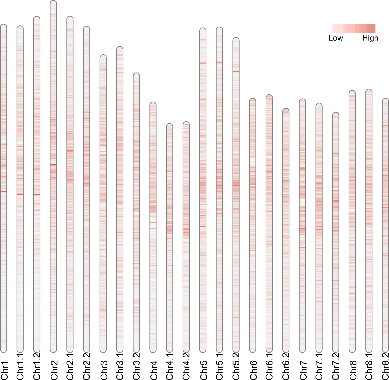 | 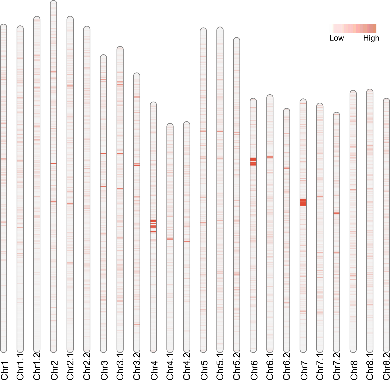 |
| --- | --- | --- |
| 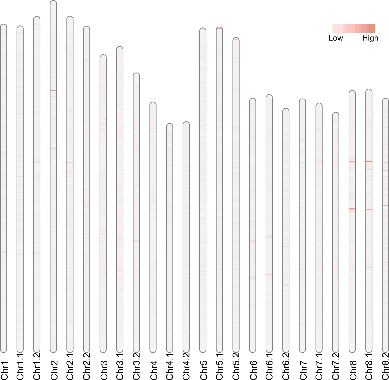 | 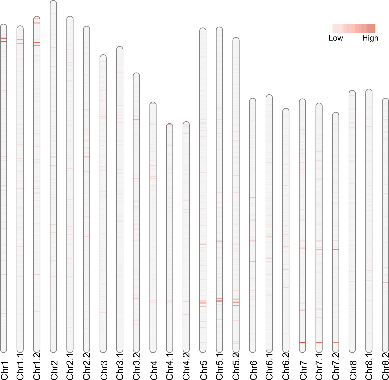 | 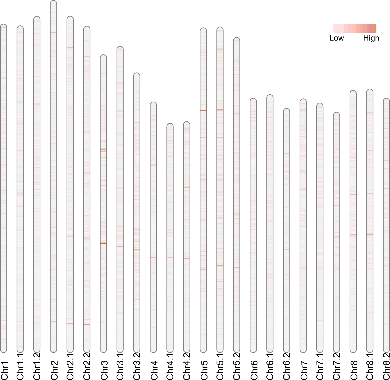 |
| 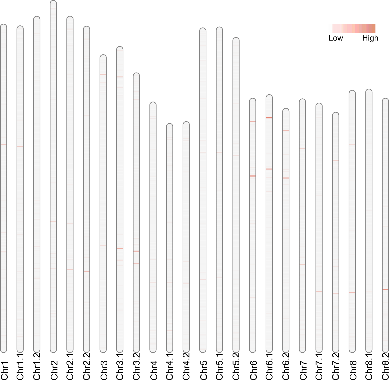 | 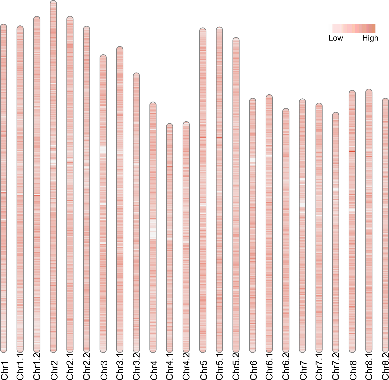 | 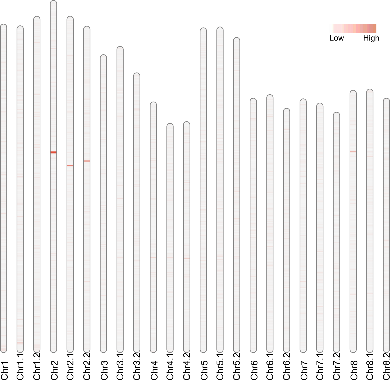 |

**Fig. S18** The distribution heatmap of repeat elements density across three haplomes. The density for each type of elements (Gypsy, Copia, EnSpms, hAT, Helitron, LINE (Long Interspersed Nuclear Element), SINE (Short Interspersed Nuclear Element), MITEs, and MuDRs) was caculated using a non-overlap sliding window approach with window size of 100kb. The ideogram labels ended with ‘.1’ are chromosomes of S_7_ haplome, with ‘.2’ are chromosomes of S_8_ haplome. The other ideograms denote S_C_ haplome of homozygous *A.majus*.

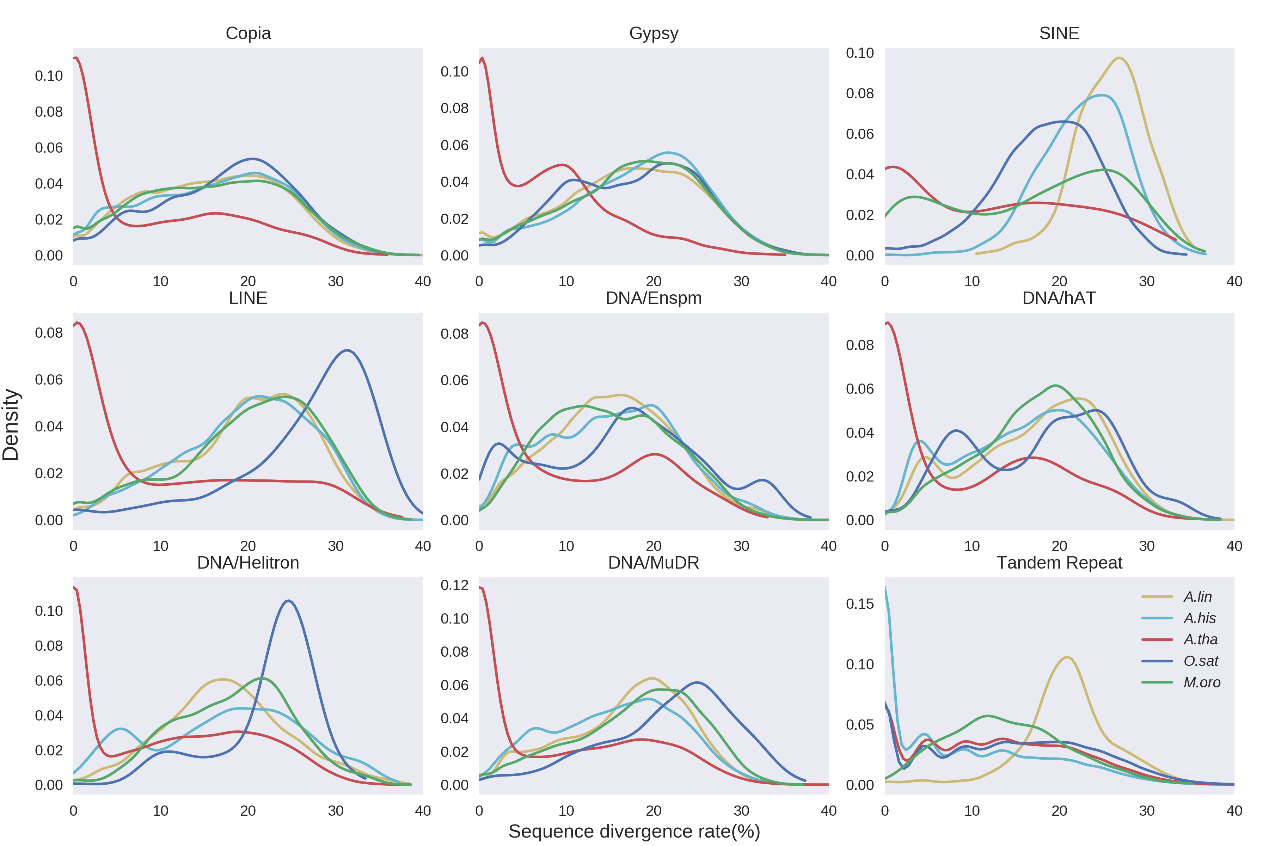

**Fig. S19** Denstity plot of divergence rates of different transposable element in different genomes.

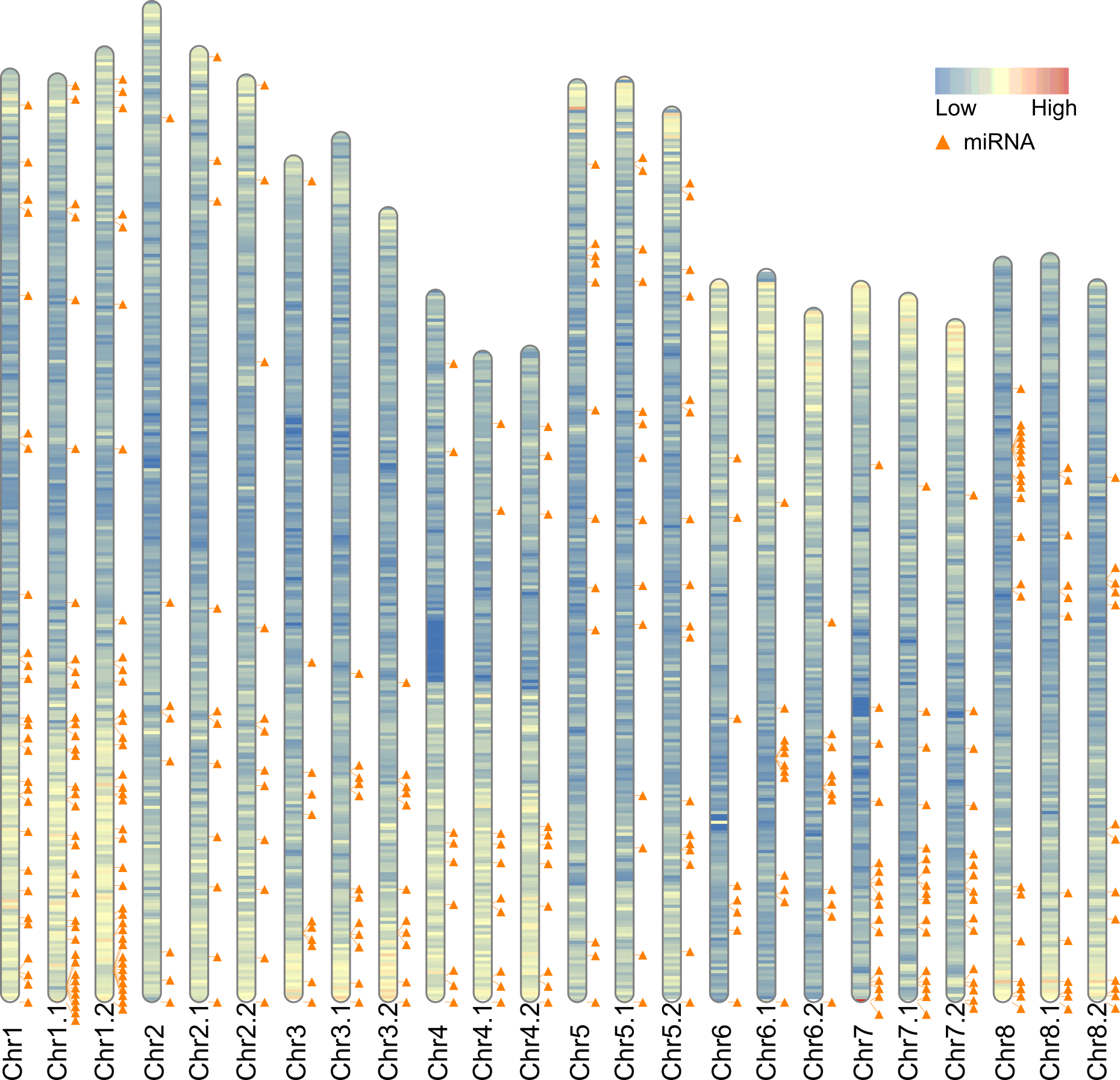

**Fig. S20** Gene density heatmap of 3 haplomes (S_7_, S_8_, and S_C_). The ideogram labels ended with ‘.1’ and ‘.2’ denote chromosomes of S_7_ haplome and S_8_ haplome of SI-Ah respectively. The other ideogram represent SC-Am S_C_ haplome. Chromosomes were binned into 100kb window to display the gene distribution of assembled chromosomes. Also, the miRNA location of 3 different haplotype genomes were marked along the chromosome using yellow triangle.

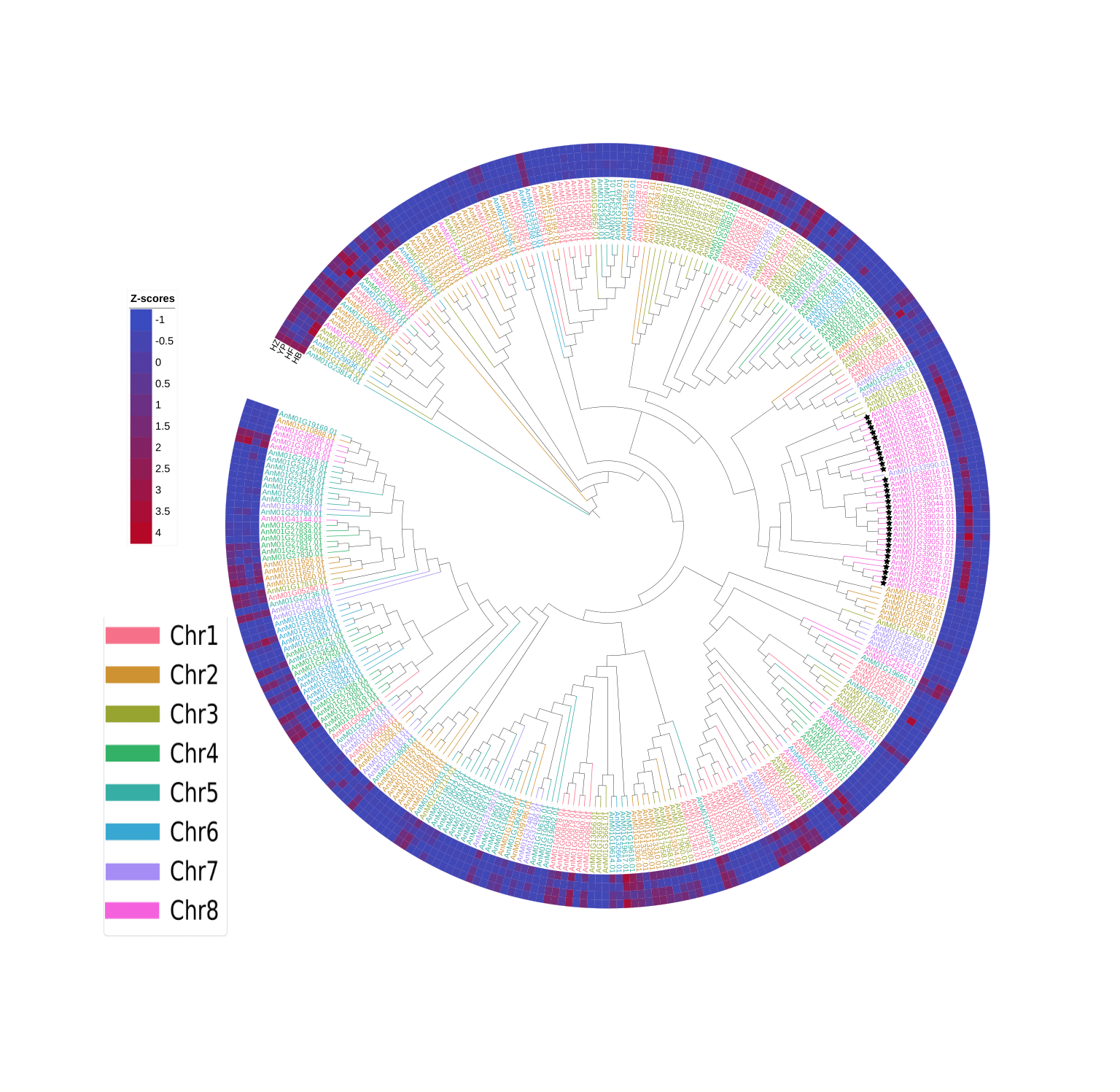

**Fig. S21** Phylogenetic tree reconstructed based on the amino acid sequences of all F-box genes from *A.majus* genome. The maximal likelihood tree was calculated using raxml-ng with 1000 bootstrap and GTR+G substitution model. The color of gene ID denoted the chromosome the gene within as indicated in the legend. Heatmap color represent Z-scores derived from RNA-seq expression data for each gene. SLF genes are marked with black star.

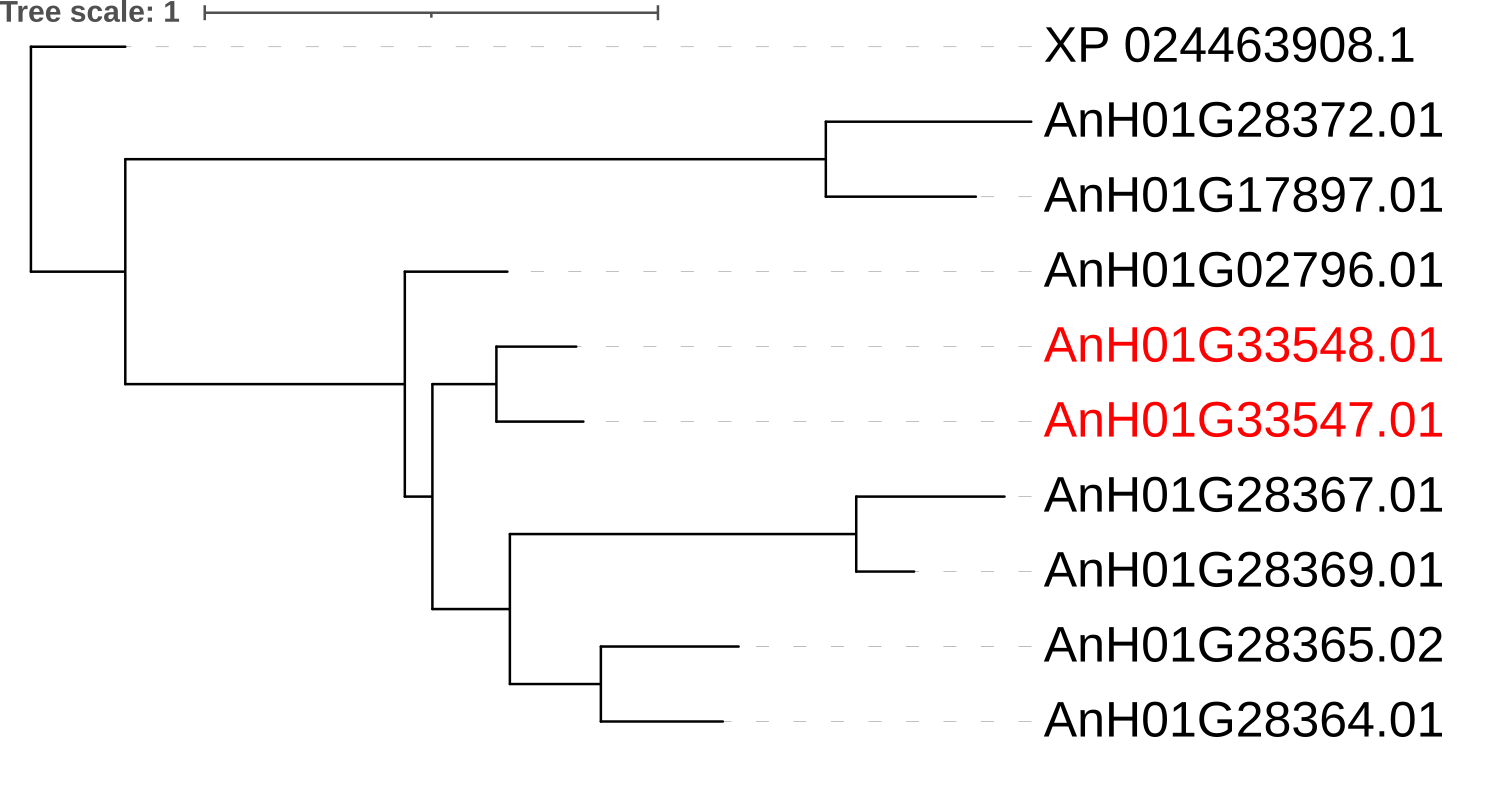

**Fig. S22** Phylogenetic tree reconstructed based on the amino acid sequences of all organ specific protein genes from *A.hispanicum* genome with XP_024463908.1 as outgroup, using raxml-ng.

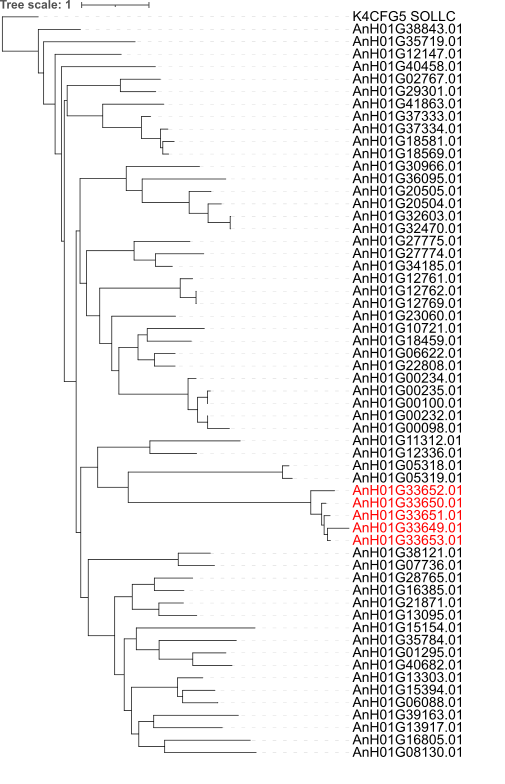

**Fig. S23** Phylogenetic tree reconstructed based on the amino acid sequences of all phytocyanin domain like genes from *A.hispanicum* genome with K4CFG5 as outgroup, using raxml-ng.

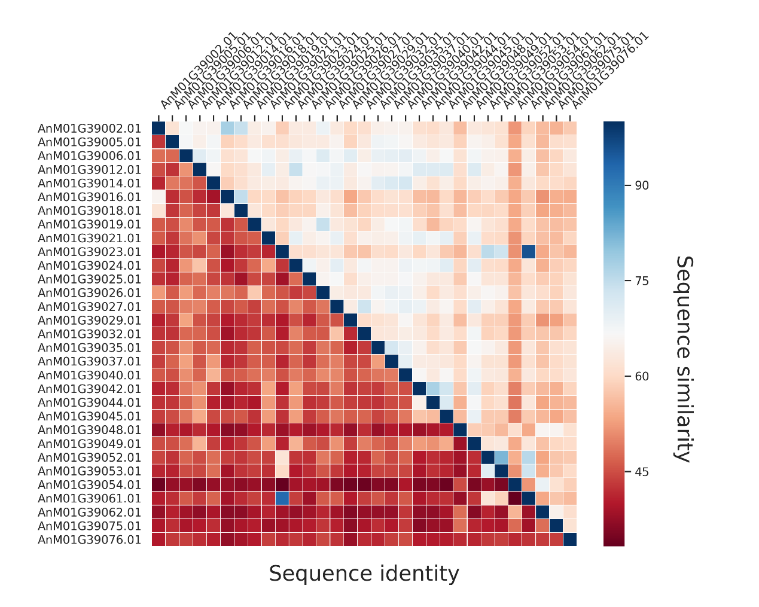

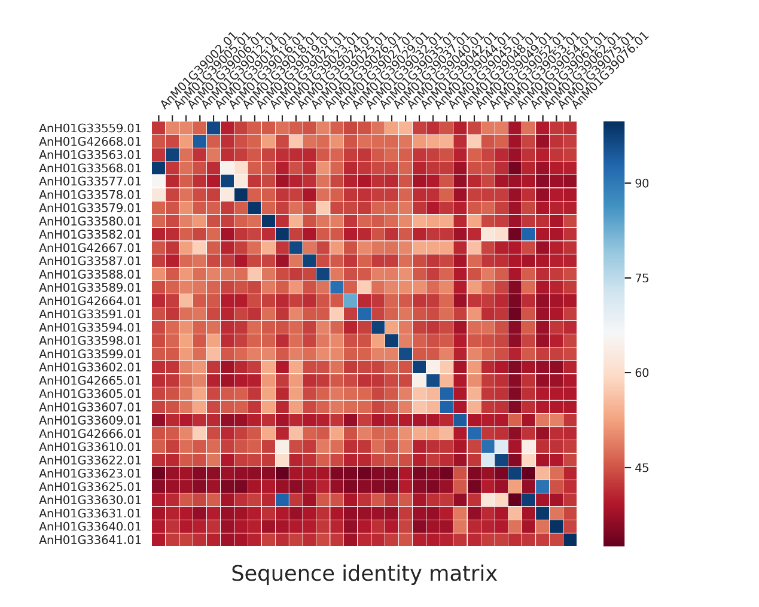

**Fig. S24** Heatmap showing sequence identity (lower left triangle) and sequence similarity (upper right triangle) between *SLFs* in SC-Am, and SI-Ah vs SC-Am. The genes in y-axis are ordered by position on the chromosome.

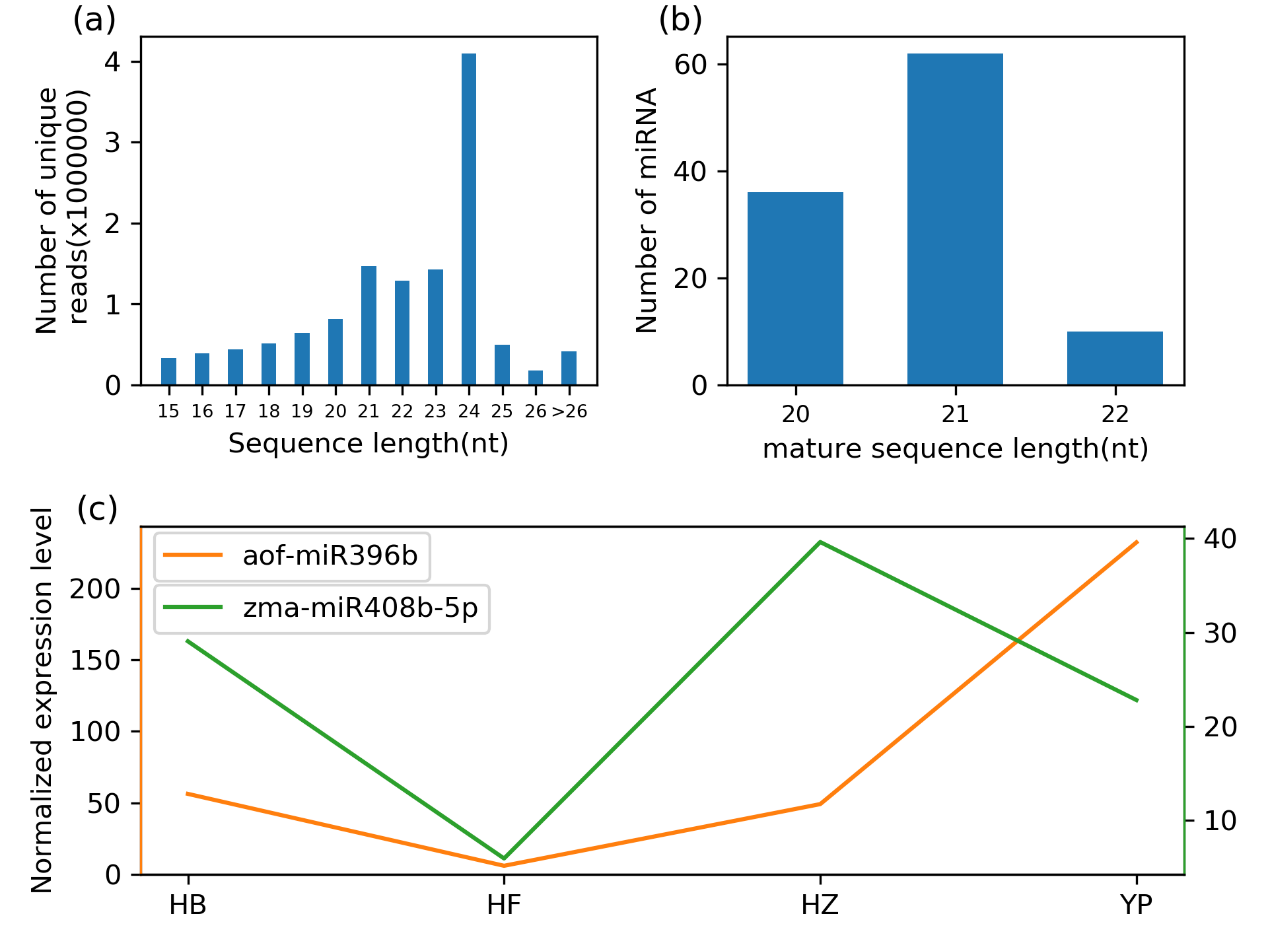

**Fig. S25 (a)** Length distribution of unique sRNA in four libraries of SI-Ah. Most of the generated reads were 24 and 21 nucleotides. **(b)** Count number of identified miRNAs in SI-Ah. **(c)** Expression level in four tissues of two miRNAs which may play a role in regulate expression of the MYB-related transcription factor, *AnH01G33501.01*.

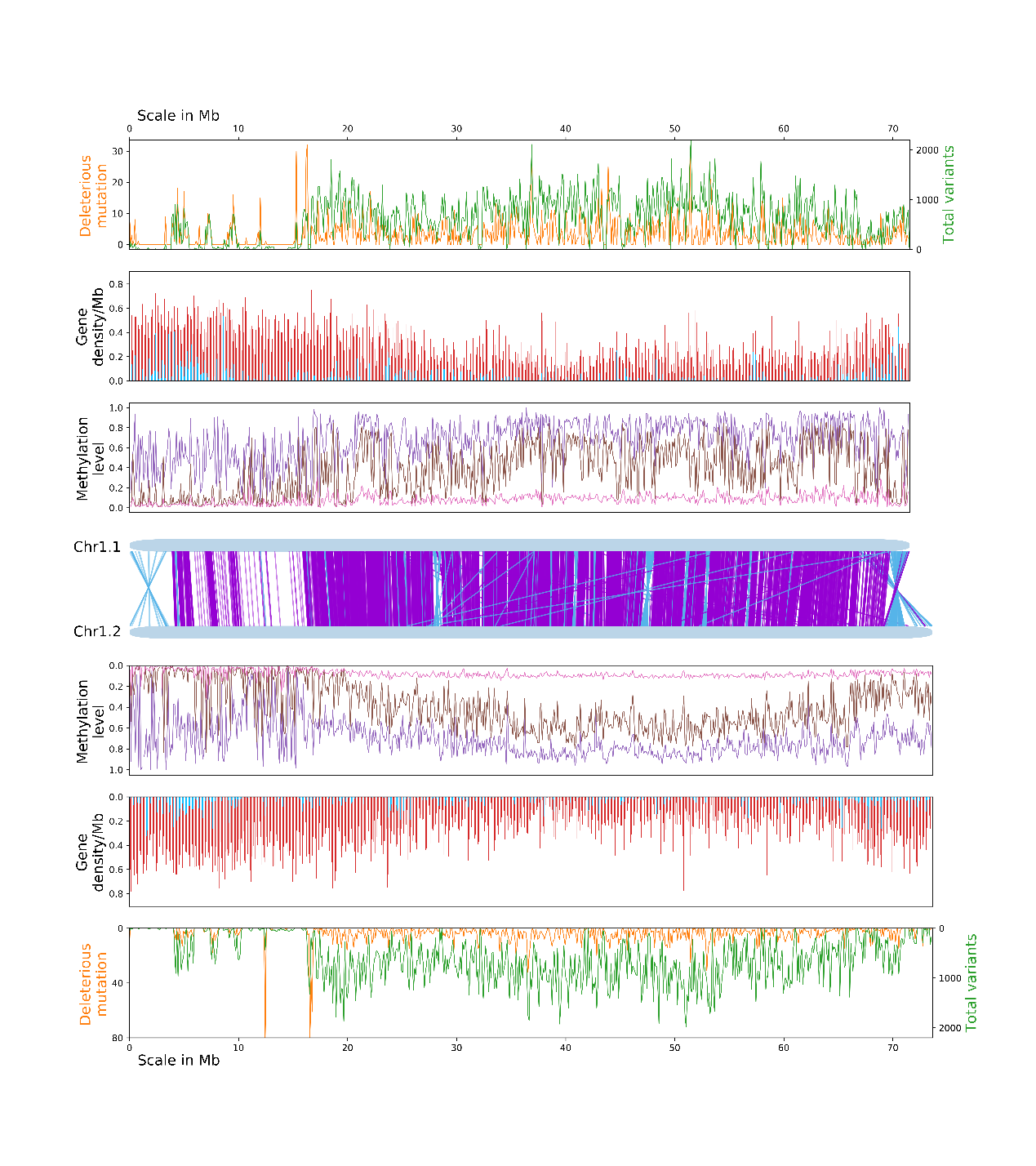

**Fig. S27** Alignment of SI-Ah chromosome 1 from two haplomes. The central light blue bars represent homologous chromosome 1 with the purple and dark blue lines indicating the forward and reverse alignments. The distribution of total variants(green), deleterious mutations (orange), annotated genes (pink), imbalanced expressed alleles (blue), methylation level of three contexts are arranged symmetrically for each haplotype. The methylation level of methylated sites in CpG, CHG and CHH contexts are indicated in purple, brown, and magenta respectively. All of the numbers were determined in 100 kb windows.

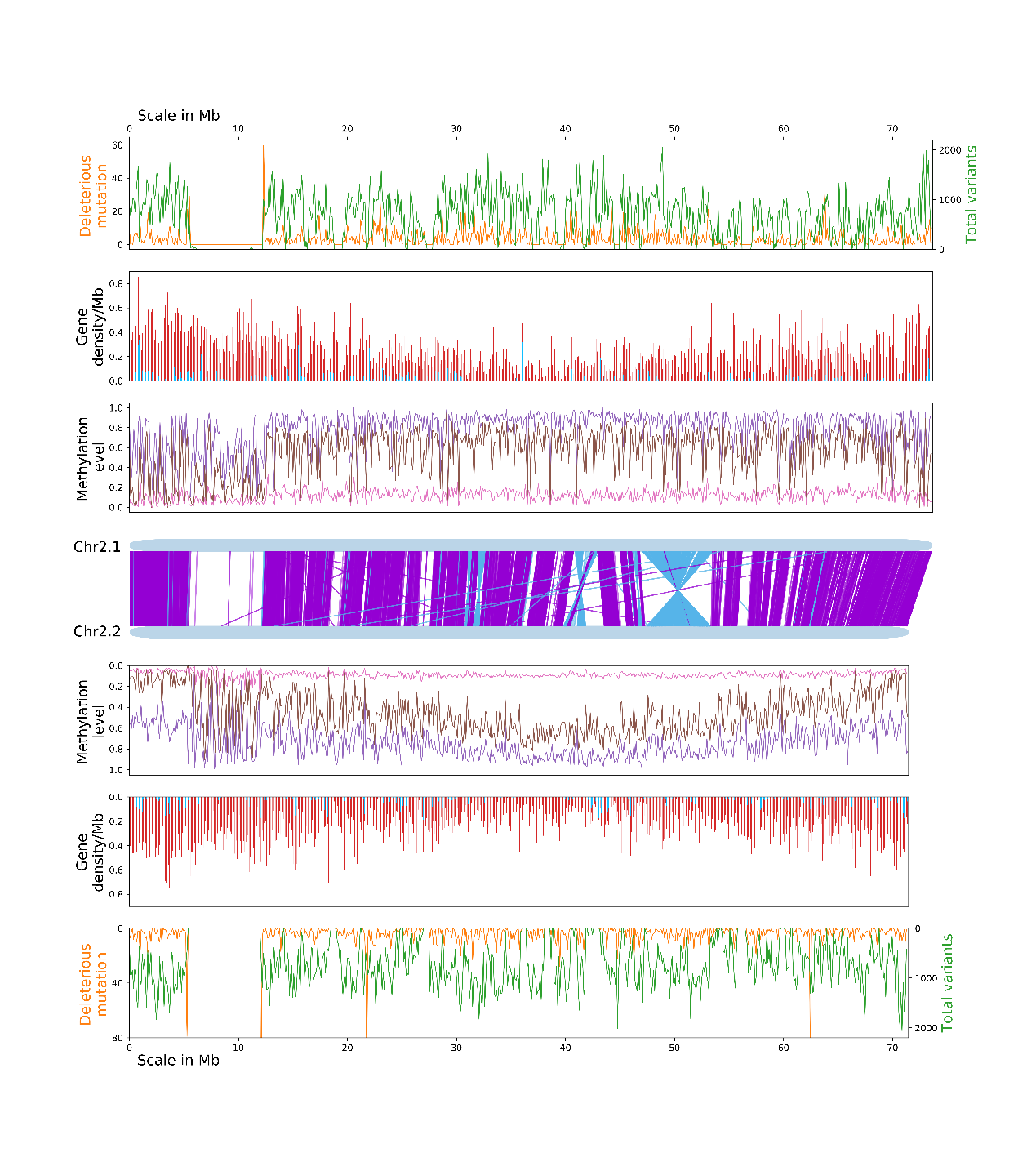

**Fig. S28** Alignment of SI-Ah chromosome 2 from two haplomes. The central light blue bars represent homologous chromosome 2 with the purple and dark blue lines indicating the forward and reverse alignments. The distribution of total variants(green), deleterious mutations (orange), annotated genes (pink), imbalanced expressed alleles (blue), methylation level of three contexts are arranged symmetrically for each haplotype. The methylation level of methylated sites in CpG, CHG and CHH contexts are indicated in purple, brown, and magenta respectively. All of the numbers were determined in 100 kb windows.

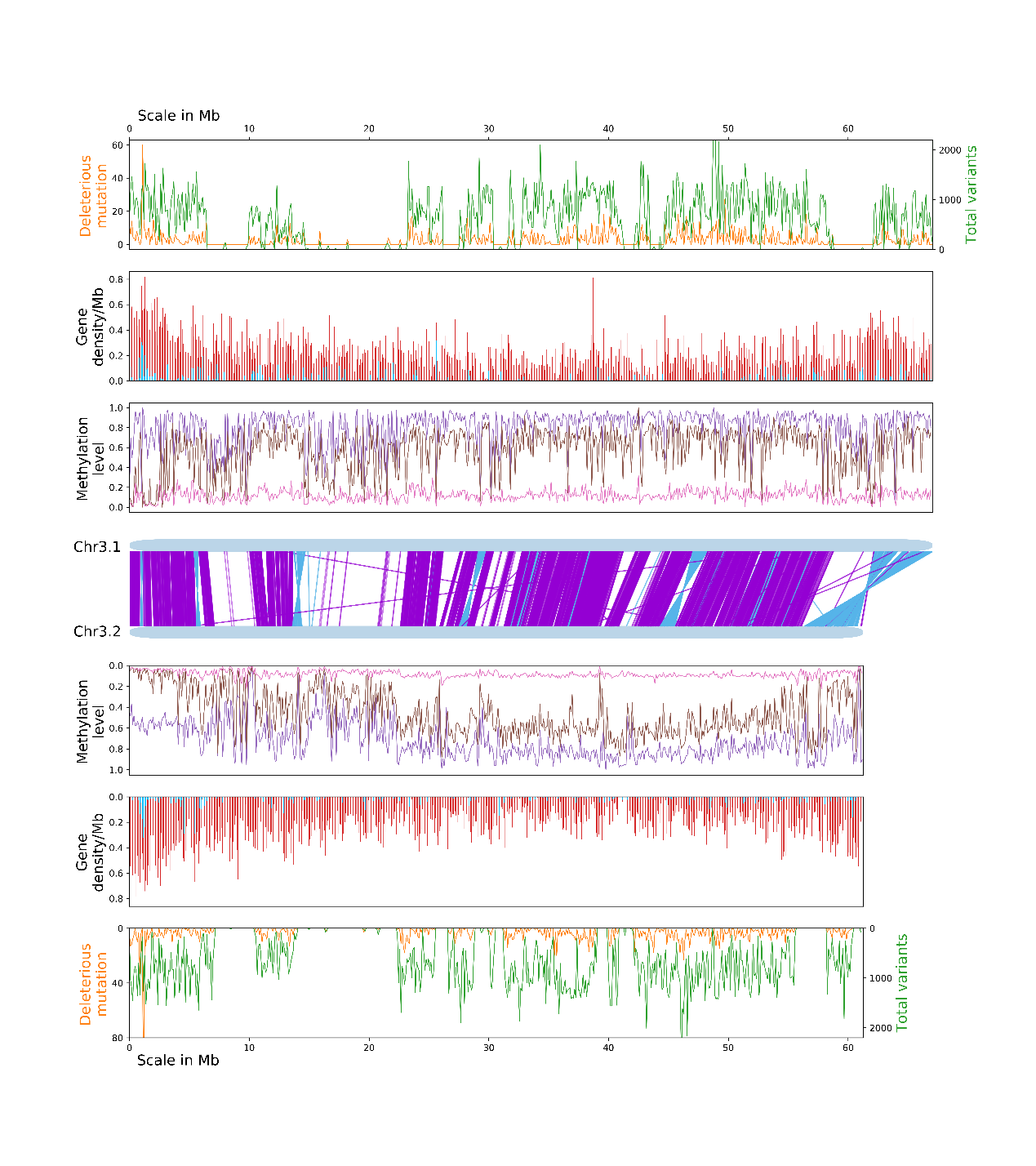

**Fig. S29** Alignment of SI-Ah chromosome 3 from two haplomes. The central light blue bars represent homologous chromosome 3 with the purple and dark blue lines indicating the forward and reverse alignments. The distribution of total variants(green), deleterious mutations (orange), annotated genes (pink), imbalanced expressed alleles (blue), methylation level of three contexts are arranged symmetrically for each haplotype. The methylation level of methylated sites in CpG, CHG and CHH contexts are indicated in purple, brown, and magenta respectively. All of the numbers were determined in 100 kb windows.

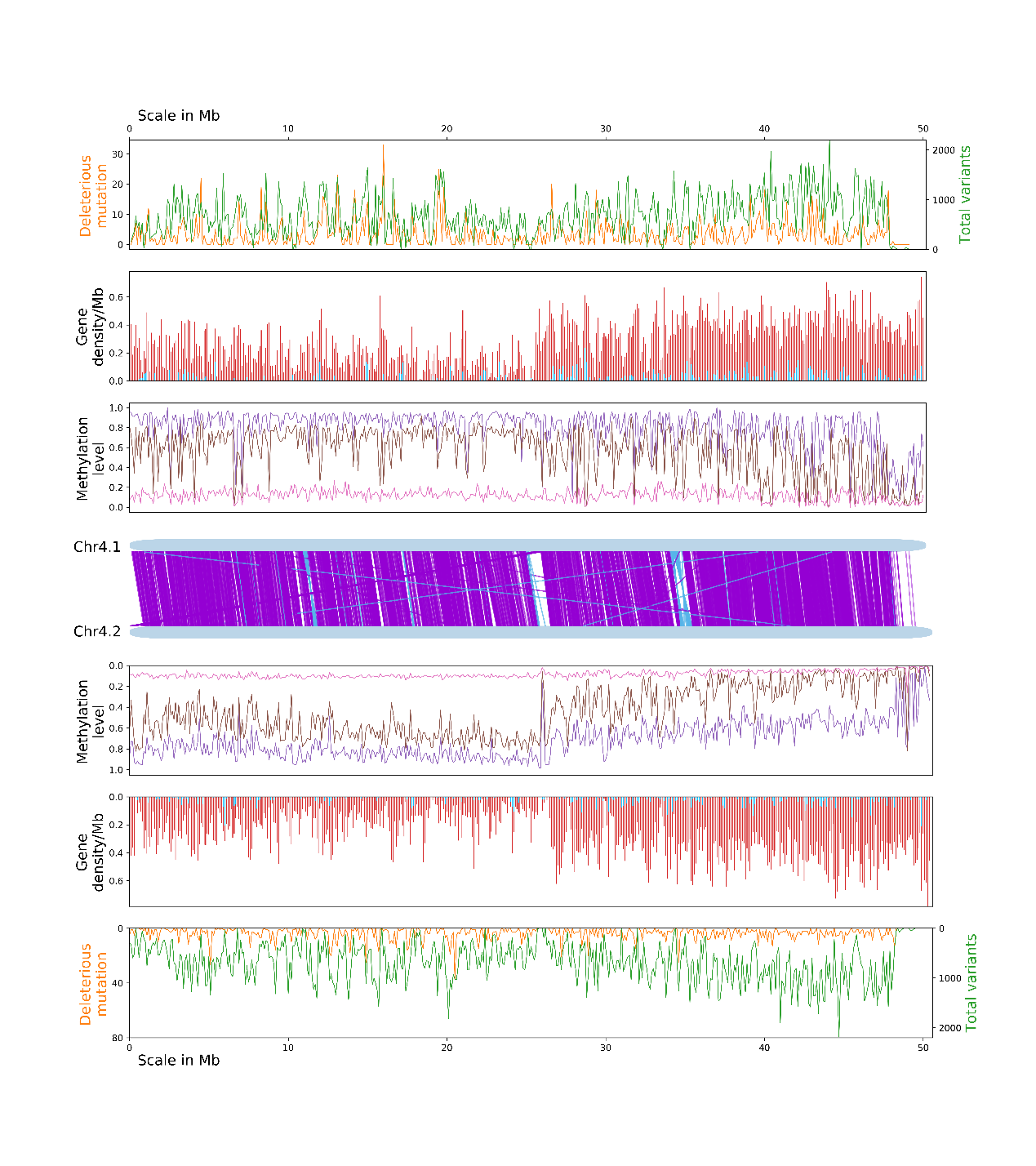

**Fig. S30** Alignment of SI-Ah chromosome 4 from two haplomes. The central light blue bars represent homologous chromosome 4 with the purple and dark blue lines indicating the forward and reverse alignments. The distribution of total variants(green), deleterious mutations (orange), annotated genes (pink), imbalanced expressed alleles (blue), methylation level of three contexts are arranged symmetrically for each haplotype. The methylation level of methylated sites in CpG, CHG and CHH contexts are indicated in purple, brown, and magenta respectively. All of the numbers were determined in 100 kb windows.

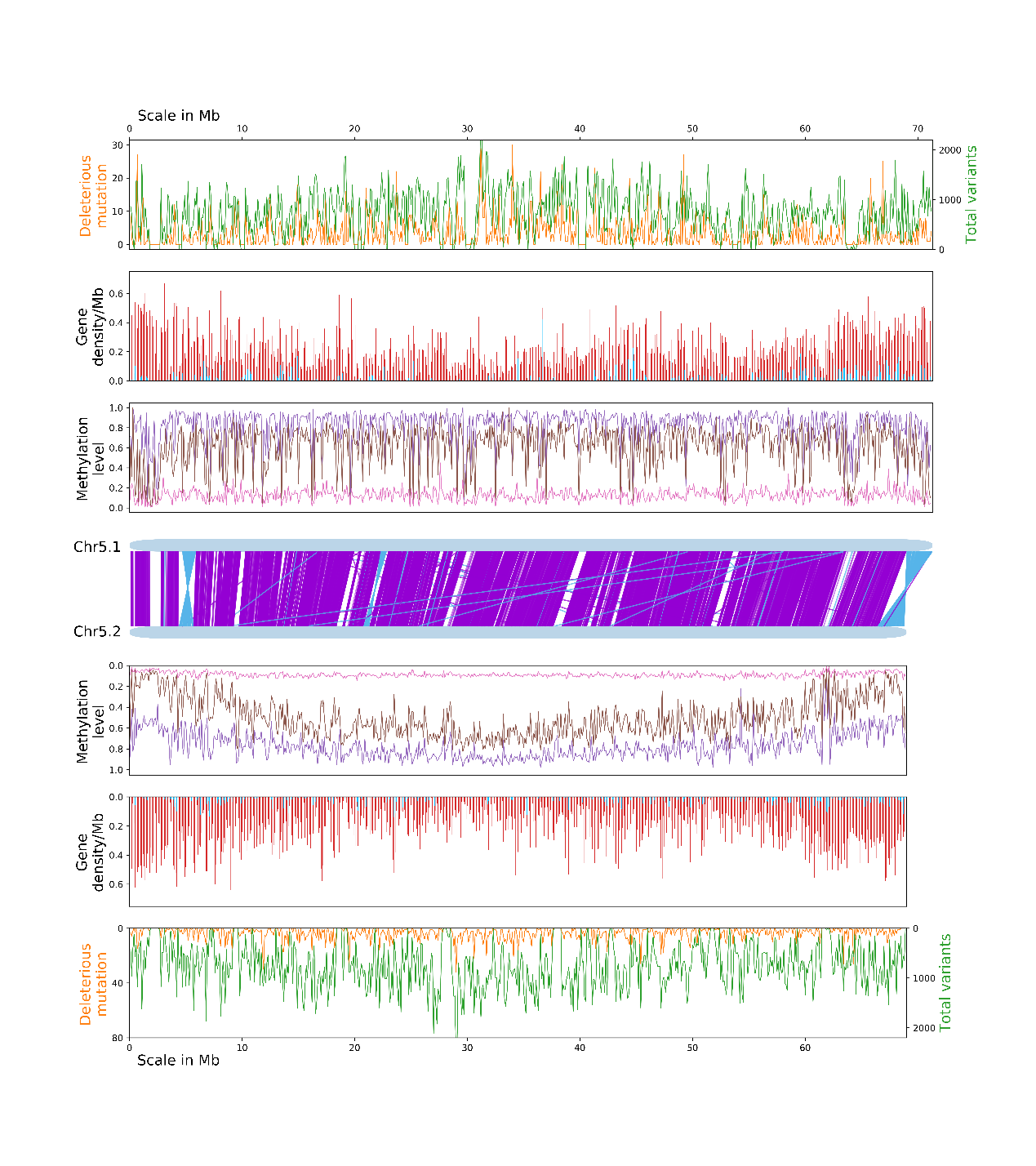

**Fig. S31** Alignment of SI-Ah chromosome 5 from two haplomes. The central light blue bars represent homologous chromosome 5 with the purple and dark blue lines indicating the forward and reverse alignments. The distribution of total variants(green), deleterious mutations (orange), annotated genes (pink), imbalanced expressed alleles (blue), methylation level of three contexts are arranged symmetrically for each haplotype. The methylation level of methylated sites in CpG, CHG and CHH contexts are indicated in purple, brown, and magenta respectively. All of the numbers were determined in 100 kb windows.

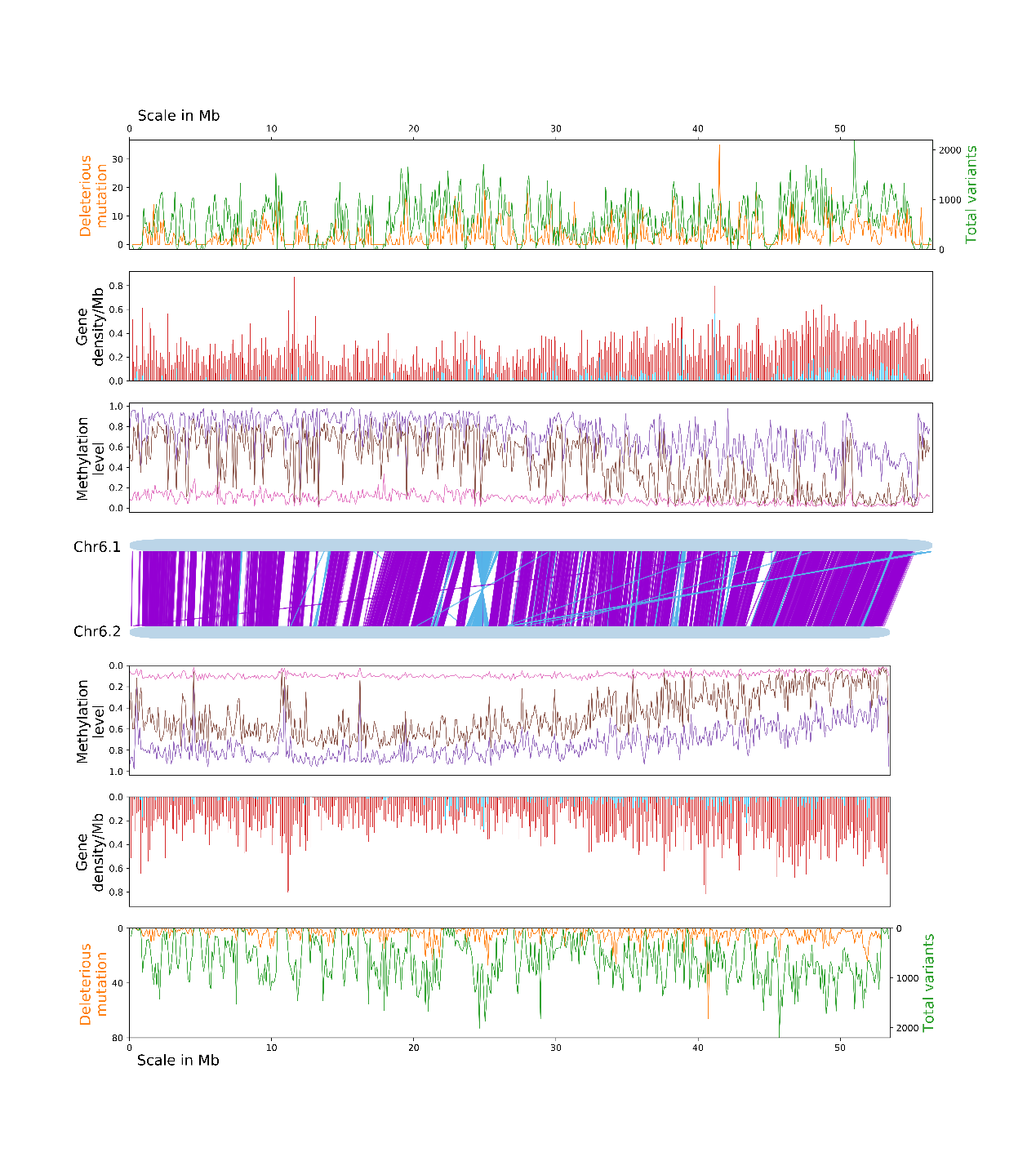

**Fig. S32** Alignment of SI-Ah chromosome 6 from two haplomes. The central light blue bars represent homologous chromosome 6 with the purple and dark blue lines indicating the forward and reverse alignments. The distribution of total variants(green), deleterious mutations (orange), annotated genes (pink), imbalanced expressed alleles (blue), methylation level of three contexts are arranged symmetrically for each haplotype. The methylation level of methylated sites in CpG, CHG and CHH contexts are indicated in purple, brown, and magenta respectively. All of the numbers were determined in 100 kb windows.

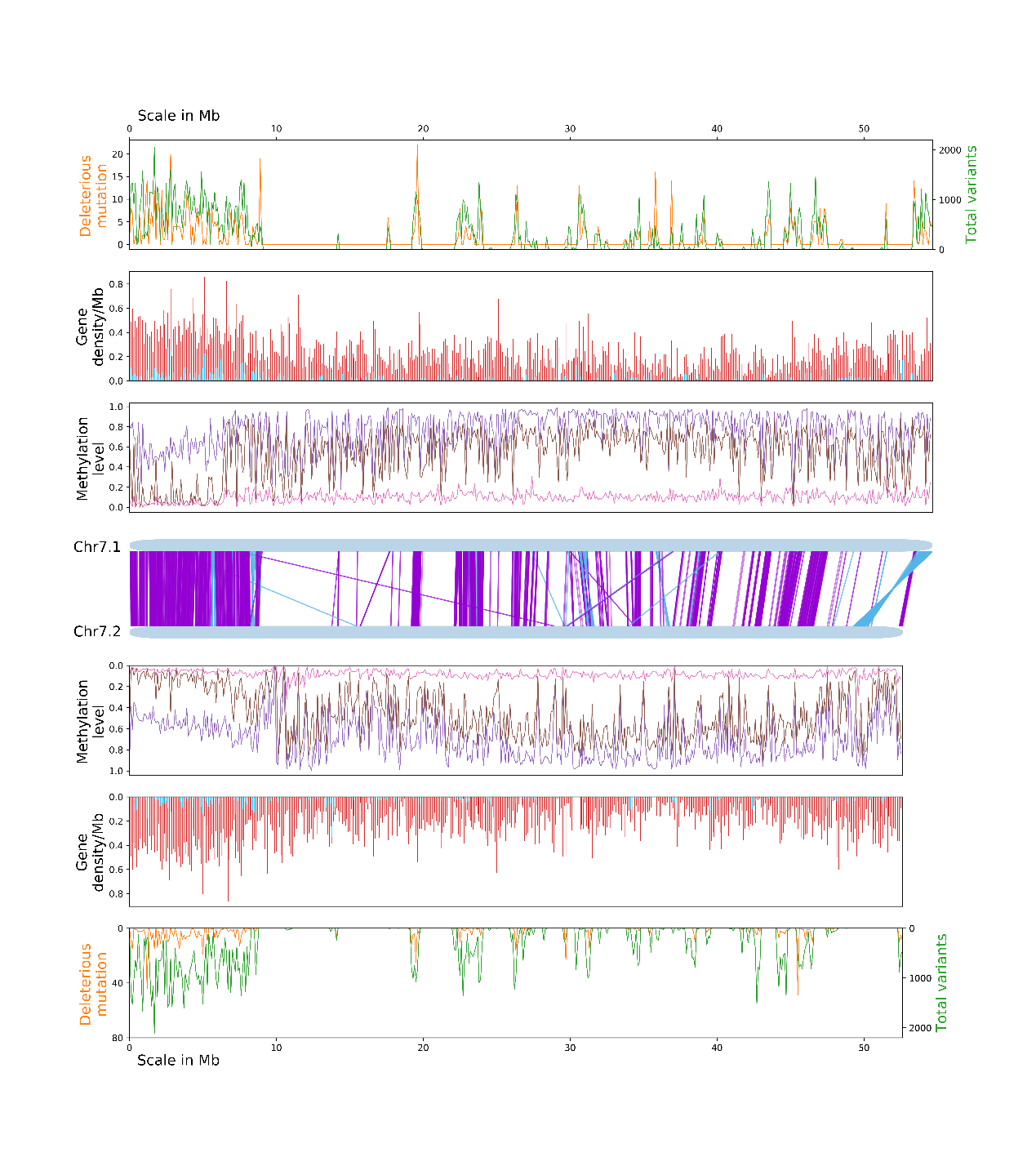

**Fig. S33** Alignment of SI-Ah chromosome 7 from two haplomes. The central light blue bars represent homologous chromosome 7 with the purple and dark blue lines indicating the forward and reverse alignments. The distribution of total variants(green), deleterious mutations (orange), annotated genes (pink), imbalanced expressed alleles (blue), methylation level of three contexts are arranged symmetrically for each haplotype. The methylation level of methylated sites in CpG, CHG and CHH contexts are indicated in purple, brown, and magenta respectively. All of the numbers were determined in 100 kb windows.

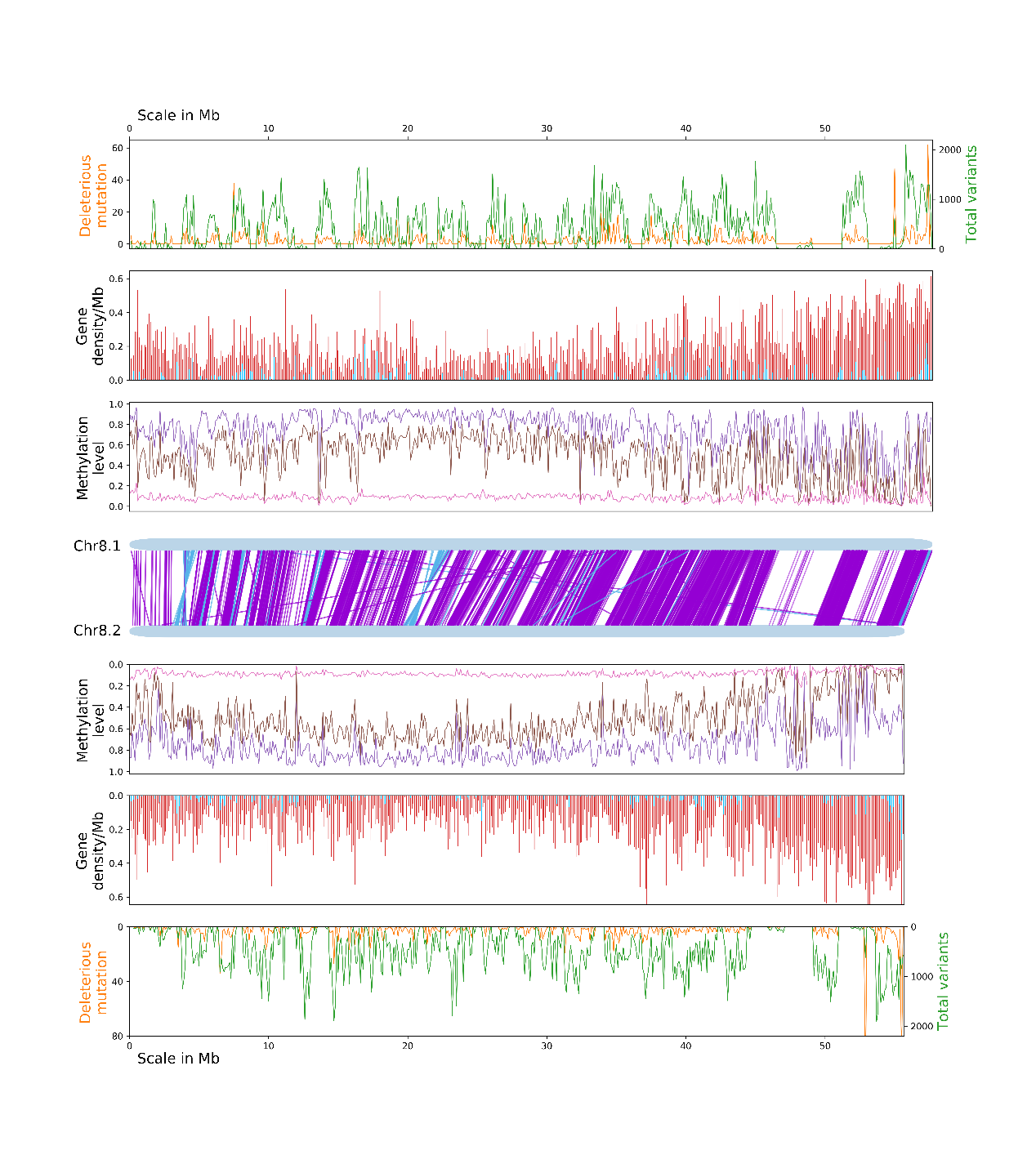

**Fig. S34** Alignment of SI-Ah chromosome 8 from two haplomes. The central light blue bars represent homologous chromosome 7 with the purple and dark blue lines indicating the forward and reverse alignments. The distribution of total variants(green), deleterious mutations (orange), annotated genes (pink), imbalanced expressed alleles (blue), methylation level of three contexts are arranged symmetrically for each haplotype. The methylation level of methylated sites in CpG, CHG and CHH contexts are indicated in purple, brown, and magenta respectively. All of the numbers were determined in 100 kb windows.

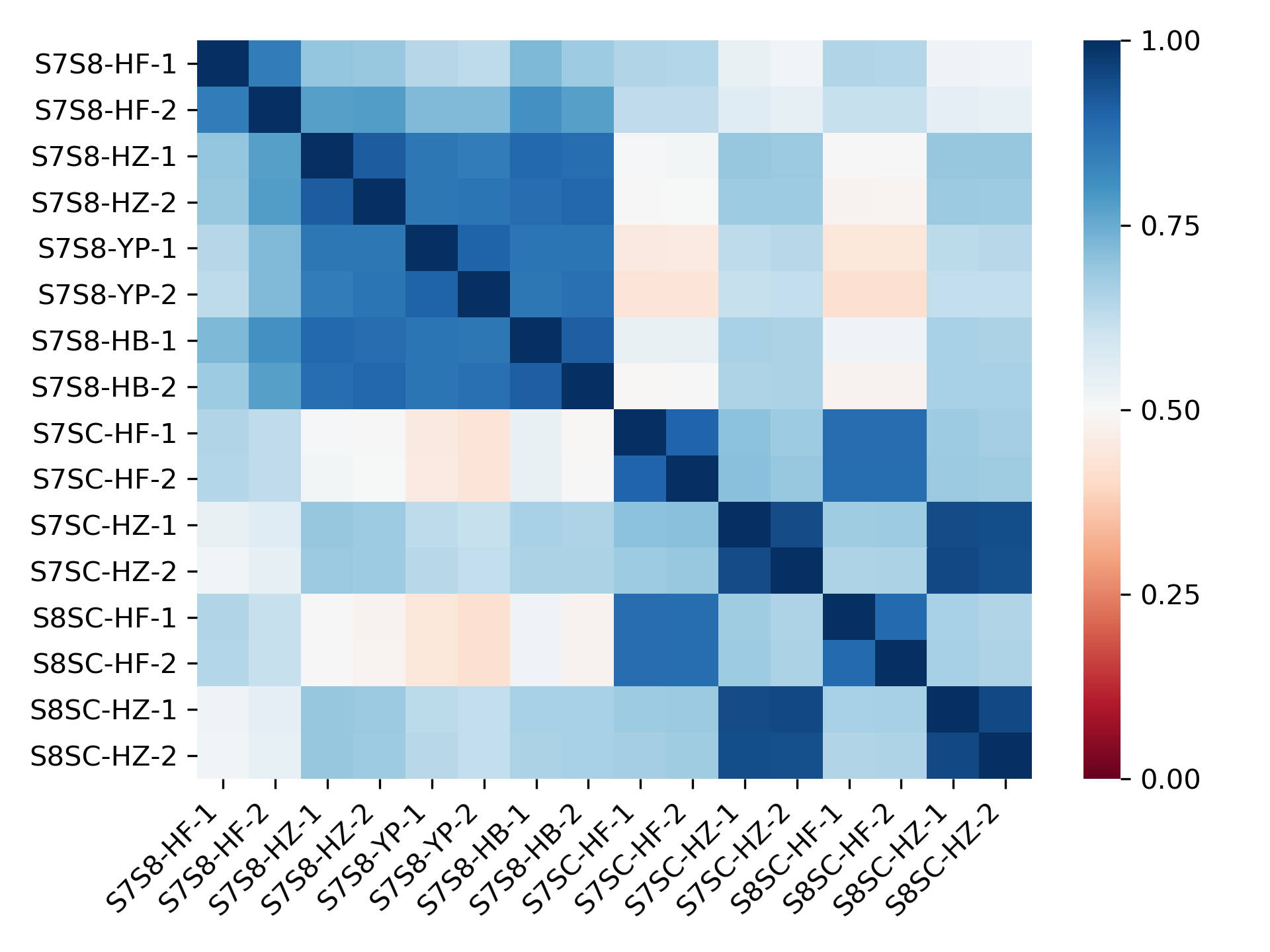

**Fig. S35** Spearman correlation heatmap of 16 RNA-seq samples. HB: petal, HF: pollen, YP: leaf, HZ: style.

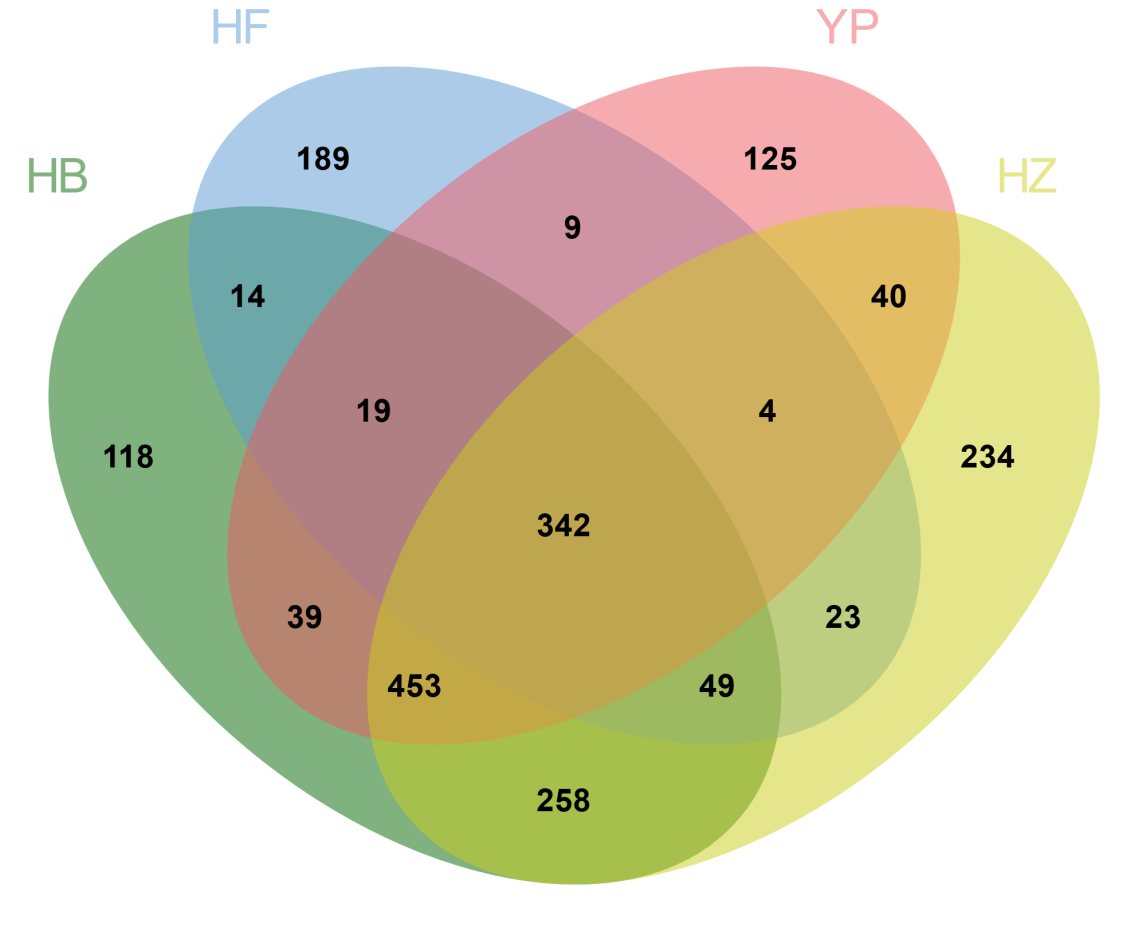

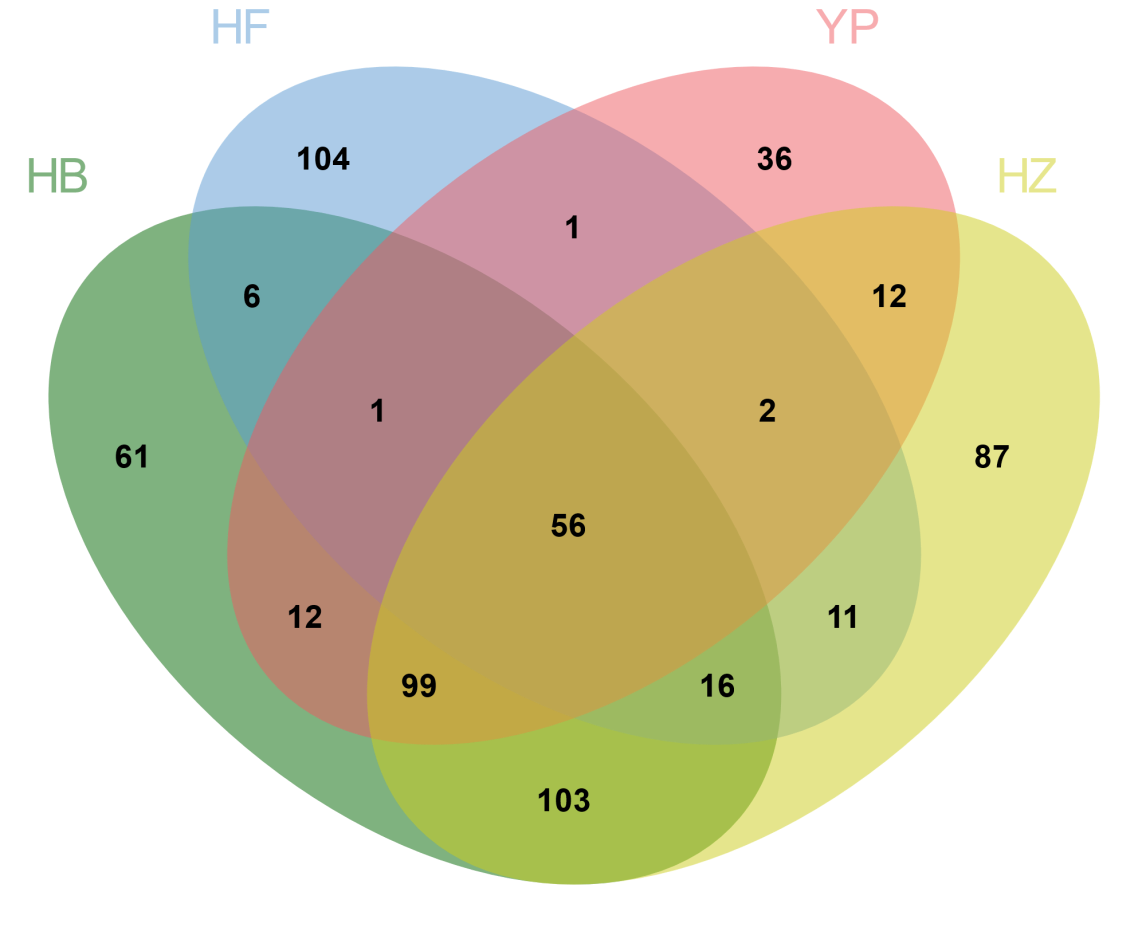

**Fig. S36** Venn diagram of imbalanced expressed gene pairs across four tissues (S7-dominant and S8-dominant). HB: petal, HF: pollen, YP: leaf, HZ: style.
